# Age-related declines in niche self-renewal factors controls testis aging and spermatogonial stem cell competition through Hairless, Imp, and Chinmo

**DOI:** 10.1101/2025.05.01.651651

**Authors:** Yu Zheng, Ya-Chien Lee, Yu-Ting Wang, Pin-Kuan Chiang, Shao-Lun Chang, Hwei-Jan Hsu, Li-Sung Hsu, Erika A. Bach, Chen-Yuan Tseng

## Abstract

Aging is associated with progressive tissue decline and shifts in stem cell clonality. The role of niche signals in driving these processes remains poorly understood. Using the *Drosophila* testis, we identify a regulatory axis in which age-related decline of niche signals (BMPs) lead to upregulation of the co-repressor Hairless, which downregulates the RNA-binding protein Imp in aged germline stem cells (GSCs). Reduced Imp causes loss of Chinmo, a key factor in GSC aging and competition. Reduced Chinmo causes ectopic Perlecan secretion which accumulates in the testis lumen and causes GSC loss. Aging of the testis is reversed by increasing BMPs in the niche, or by overexpressing Imp or depleting Hairless in GSCs. Furthermore, GSC clones with reduced Imp or increased Hairless are more competitive, expelling wild-type neighbors and monopolizing the niche. Thus, BMPs regulate testicular niche aging through the Hairless–Imp–Chinmo axis and “winning” GSCs usurp these aging mechanisms.

**Highlights:**

- Aged niche cells produce less BMPs, resulting in more Hairless (H) in aged GSCs
- Elevated H represses *Imp*, resulting in less Chinmo and in ectopic ECM secretion
- Aging is prevented by higher BMP in niche cells, or by higher Imp or lower H in GSCs
- GSCs with low *Imp* or high *H* exploit these aging mechanisms to colonize the GSC pool

## Introduction

Adult organs rely on tissue stem cells for lifelong maintenance. These stem cells typically reside in specialized microenvironments termed niches, which provide short-range signals essential for preserving stemness ^1^. However, as organisms age, niche functionality declines, resulting changes in secretion of soluble factors and in biophysical properties of the microenvironment ^2^. The age-dependent decline in niche function reduces both stem cell activity and overall stem cell numbers, leading to age-related organ dysfunction ^2,3^. In humans, the decline in fertility in older men is due in part to the dysfunction of Sertoli and Leydig cells—key components of the spermatogonial stem cell (SSC) niche ^4–9^. Mouse models similarly demonstrate age-dependent reductions in fertility and testis function, driven by decreased production of niche-derived signals such as glial cell-derived neurotrophic factor in older Sertoli cells ^10^. Consecutive transplantation of aged SSCs to the testes of young male mice maintains stem cell function for extended periods of time as the result of younger niches ^11^.

Competition among adult stem cells often occurs with aging and results from extrinsic and intrinsic events. The age-altered microenvironment can exert different selective forces on resident stem cells, and stem cells can sustain age-dependent mutations that could impart a selection advantage ^12^. Stem cell competition is implicated in a range of human pathologies. A well-known example is clonal hematopoiesis, where a small subset of hematopoietic stem cells (HSCs) expands to dominate the population in approximately 10% of older adults, markedly elevating the risk of leukemia ^13,14^. Age-related shifts in stem cell clonality have been observed in other adult somatic tissues and can underlie cancer risk ^15–25^. In the male germline, paternal age effect (PAE) disorders arise from *de novo* mutations in aging SSCs that confer a growth or competitive advantage, leading to clonal expansion and an increased proportion of mutant cells in the seminiferous tubules ^26^. These “selfish” mutations pose serious risks to offspring ^27^. Despite their clinical importance, *in vivo* models for PAE are limited, and the phenomenon of stem cell competition in the germline remains less explored than in the soma ^28^. How age-dependent changes in the niche microenvironment intersect with stem cell competitiveness remains largely uncharacterized.

The *Drosophila* testis has proven to be a powerful model for studying both SSC niche aging and SSC competition ^29–33^. Each testis houses a niche comprising approximately 12 post-mitotic cells, which support 8–10 SSCs (termed germline stem cells (GSCs) in flies) (**Fig. 1A**) ^34^. Niche cells produce the cytokine Upd, which activates the JAK/STAT pathway in GSCs and promotes adhesion to the niche but not GSC self-renewal *per se* ^35–38^. Niche cells produce less Upd declines in aged testes, contributing to decreased JAK/STAT signaling in older GSCs ^30,31^. Niche cells also produce Bone Morphogenetic Protein (BMP) ligands Decapentaplegic (Dpp) and Glass bottom boat (Gbb), which have been shown to be *bona fide* GSC self-renewal cues ^36,39–41^. Whether BMPs decline during aging in the testis niche and whether such a decline impacts age-related phenotypes in the testis remains unknown.

**Figure 1.**
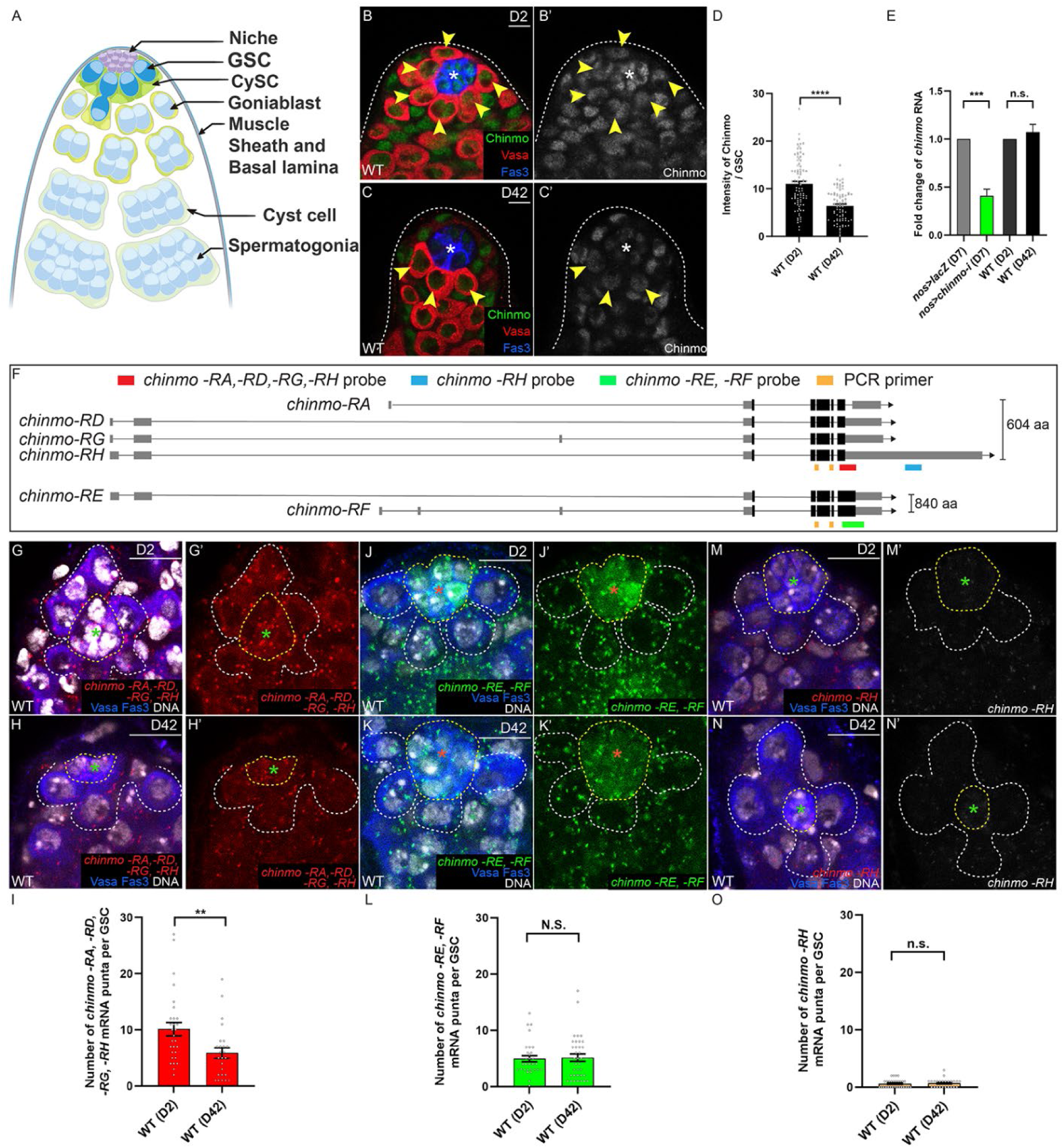
*chinmo* mRNA is post-transcriptionally regulated in GSCs during aging. (A) The adult *Drosophila* testis. A GSC makes direct contact with the niche and divides to produce a gonialblast (Gb), which undergoes transit-amplifying divisions to produce a 16-cell spermatogonia that undergoes meiosis and differentiates into sperm. CySCs divide to produce cyst cells, two of which envelope a Gb and its descendants. (B-D) Confocal images of WT testes in young (D2) (A) and aged (D42) (B) testes stained for. Chinmo (green). Arrowheads indicate GSCs. (C) Graph showing the relative intensity of Chinmo protein between D2 (A) and D42 (B) GSCs. (E) Graph showing the relative change of *chinmo* mRNA in *nos>lacZ* (control, gray) versus *nos>chinmo-i* (green) GSCs at D7 and the relative change of *chinmo* mRNA in D2 and D42 GSCs. (F) Schematic of the six *chinmo* mRNA transcript (*-RA*, *-RD*, *-RE*, *-RF, -RG*, *-RH*). Exons (gray boxes), introns (lines), and coding regions (black boxes) are shown. *chinmo-RA*, *-RD*, *-RG*, and *-RH* encode a 604 aa protein, whereas *chinmo-RE* and *-RF* encode an 840 aa protein. The orange boxes indicates the PCR primers spanning all *chinmo* transcripts. HCR probes for *chinmo* isoforms are shown in red for *-RA, -RD, -RG, -RH*, in blue for *-RH*, and in green for *-RE, -RF*. (G-O) Confocal images of HCR-FISH for *chinmo* mRNA in D2 (G, J, M) or D42 (H, K, N) in WT testes. Dashed white and yellow lines indicate GSCs and niche cells, respectively. Arrowheads indicate GSCs. (G-I) Probe for *chinmo-RA, -RD, -RG*, and *-RH* transcripts (red) in F (D2) and G (D42), quantified in I. (J-L) Probe for *chinmo-RE* and *-RF* transcripts (green) in I (D2) and J (D42), quantified in L. (M-O) Probes for *chinmo-RH* transcripts (red) in L (D2) and M (D42), quantified in O. Asterisks in indicate the niche. DAPI labels DNA; Vasa labels germ cells; Fas3 labels niche cells. Scale bars = 10 μm. In D, E, I, L, and O, error bars represent SEM. ** *P* ≤ 0.01; *** *P* ≤ 0.001; **** *P* ≤ 0.0001; n.s. indicates no significance, as determined by Student’s t-test (see STAR Methods).

We previously reported that levels of the transcription factor Chinmo in germline stem cells (GSCs) regulate aging of the testis stem cell niche ^33^. During aging, GSCs exhibit a progressive and significant decline in Chinmo expression. As Chinmo levels decrease, expression of its target genes—including those encoding the secreted extracellular matrix (ECM) protein Perlecan (Pcan) and the ECM-binding protein Dystroglycan (Dg)—significantly increases. These age-dependent transcriptional changes drive: (1) the formation of an ECM structure around the niche, termed the “moat”; (2) a substantial accumulation of Dg at the GSC–niche interface; and (3) a marked reduction in the number of GSCs. Notably, maintaining robust Chinmo levels in GSCs during adulthood can rejuvenate the aging testis stem cell niche.

In the same study, we also found that GSC clones homozygous mutant for *chinmo* exhibit a striking competitive advantage over wild-type (WT) neighbors, ultimately monopolizing the entire germline and leading to biased transmission of the chinmo-mutant allele ^33^. Mechanistically, this advantage stems from hijacking the aging-associated niche remodeling process: *chinmo*-mutant GSCs ectopically secrete Pcan into the testis lumen, promoting moat formation. The moat impairs the ability of neighboring WT GSCs to adhere to the niche, compromising their maintenance. Simultaneously, *chinmo*-null GSCs upregulate Dg, allowing them to anchor themselves more effectively within the remodeled niche. However, the upstream factors that regulate Chinmo expression during aging have remained unknown, limiting our understanding of both testis niche aging and GSC competition.

Here, we show that Chinmo mRNA levels in GSCs are positively regulated by the RNA-binding protein (RBP) IGF-II mRNA-binding protein (Imp). We find that *Imp* mRNA and protein levels are robust in young GSCs but decline significantly with age. Importantly, sustaining Imp expression during adulthood preserves Chinmo levels and prevents testis aging phenotypes. Further analyses reveal that *Imp* is transcriptionally downregulated by the co-repressor Hairless (H), as *Imp* mRNA levels increase when H is depleted in aged GSCs. Manipulating H levels in GSCs modulates aging: loss of H alleviates aging, whereas overexpression of H accelerates it. Notably, H expression in GSCs is itself repressed by Bone Morphogenetic Protein (BMP) signals, specifically Dpp and Gbb, secreted by the niche. As BMP production declines with age, H levels rise in GSCs, leading to reduced Imp and subsequently Chinmo, ultimately driving ectopic Pcan secretion and niche remodeling. Thus, the age-dependent decline of niche-derived BMP signals converges on the H–Imp–Chinmo axis to regulate aging of the testis stem cell niche.

Finally, we demonstrate that GSC clones with lower Imp or higher H levels exhibit enhanced competitiveness, displacing neighboring WT GSCs from the niche similarly to chinmo-mutant GSCs. Our findings reveal that competitive GSCs exploit the age-dependent intrinsic and extrinsic changes regulating Chinmo expression, leading to stem cell niche disruption and eventual monoclonality of the GSC lineage.

## Results

### Chinmo is post-transcriptionally regulated in GSCs during aging

We previously reported that Chinmo protein in GSCs significantly declines during adulthood (**Fig. 1B-D**) ^33^. Four *chinmo* mRNA isoforms (*chinmo*-*RA,-RD,-RG,-RH*) encode a 604 amino acid (aa) protein, and two isoforms (*chinmo-RE* and *-RF*) encode 840 aa protein (**Fig. 1F**). We established that *chinmo* is not transcriptionally regulated in GSCs during aging (**Fig. 1E**, third and fourth bars). This assay is robust because germline depletion of *chinmo* using an RNAi transgene targeting all isoforms and the GSC-specific driver *nos-Gal4* (*nos>chinmo-i*) caused a significant decline in *chinmo* mRNA abundance compared to control *nos>lacZ* testes (**Fig. 1E**, first and second bars), Thus, *chinmo* appears to be regulated post-transcriptionally in GSCs.

To confirm this, we used hybridization chain reaction (HCR) fluorescence *in situ* hybridization (FISH) to determine *chinmo* mRNA isoform expression patterns. We used three probes: one for *chinmo-RA, -RD, -RG,* and *-RH*; a second for *chinmo-RE* and *-RF*; and a third for *chinmo-RH* (**Fig. 1F**). We found that *chinmo-RA, -RD, -RG,* and *-RH* isoforms are robustly expressed in GSCs from young 2-day-old (D2) males and decline significantly in GSCs from aged 42-day-old (D42) males (**Fig. 1G-I**). *chinmo-RE* and *-RF* are moderately expressed in GSCs regardless of age (**Fig. 1J-L**), while *chinmo-RH* is expressed at very low levels in GSCs regardless of age (**Fig. 1M-O**). *chinmo-RH* is the only isoform known to be targeted by microRNAs (miRs) (i.e., *mir-let7*) through its 6 kb extended *3’UTR* ^42^. The fact that *chinmo-RH* is not regulated by aging in GSCs suggests that miRNAs are not responsible for *chinmo* transcript downregulation during aging. Indeed, Chinmo protein was not altered in GSCs deficient in miRNA processing components *Dcr-1*, *AGO1* or *drosha* or in *mir-let7* (**Fig. S1A-C, E-I**). Additionally, *AGO1*-mutant GSCs cannot be recovered at 14 days post-clone induction (dpci) (**Fig. S1D** and **Table S1 #6**), as previously reported ^43^, indicating that they are hypocompetitive. Finally, although Chinmo is a JAK/STAT target gene in other cells ^44^, STAT-deficient GSCs still express Chinmo protein, indicating that Chinmo is not regulated by the JAK/STAT in male GSCs (**Fig. S1J-M**). Taken together, these results indicate that in male GSCs *chinmo-RA*, *-RD* and *-RG* are downregulated during aging through a process that does not involve miRs.

### Imp is required for GSC homeostasis during aging by promoting Chinmo protein expression

The RBP Imp promotes Chinmo expression in neuroblast lineages and binds *chinmo-RD* mRNA in S2 cells ^45,46^. We found that *Imp* mRNA is significantly decreased both in number and size of puncta in GSCs from aging D28 males compared to young D2 males (**Fig. 2A-D**). Imp-specific antibodies and protein traps revealed that Imp is robustly expressed in GSCs in WT young D2 males and significantly declines in GSCs from WT aged D42 males (**Fig. 2E-L** and **S2A-C**). As expected, Imp levels in niche cells declined during aging (**Fig. 2A-C** and **S2A, B, D, E**) ^31^. To test whether sustained expression of Imp can augment Chinmo expression in aged GSCs, we over-expressed Imp in GSCs throughout adulthood using *nos-Gal4*. Exogenous Imp significantly increased Imp protein in aged GSCs but not in aged niche cells (**Fig. S2D-G**) and significantly increased Chinmo protein in aged GSCs (**Fig. 2M-O**). Furthermore, depletion of *Imp* using two validated RNAi lines (**Fig. S3A-D**), caused a significant decline in Chinmo protein compared to age-matched controls (**Fig. S3E-H**). These results indicate that Imp promotes Chinmo expression in GSCs during homeostasis. We previously showed that the number of GSCs declines during adulthood and that GSC number in aged D42 males could be significantly restored by over-expressing Chinmo in GSCs ^33^. We now show that sustained Imp expression in GSCs (*nos>Imp*) throughout adulthood also restored GSC numbers in aged males (*nos>lacZ*) (**Fig. 2P**). By contrast, the depletion of Imp further reduced the total GSC number in aged D42 males (**Fig. S3I**). These results show that Imp regulates the number of GSCs during aging by promoting Chinmo expression.

**Figure 2.**
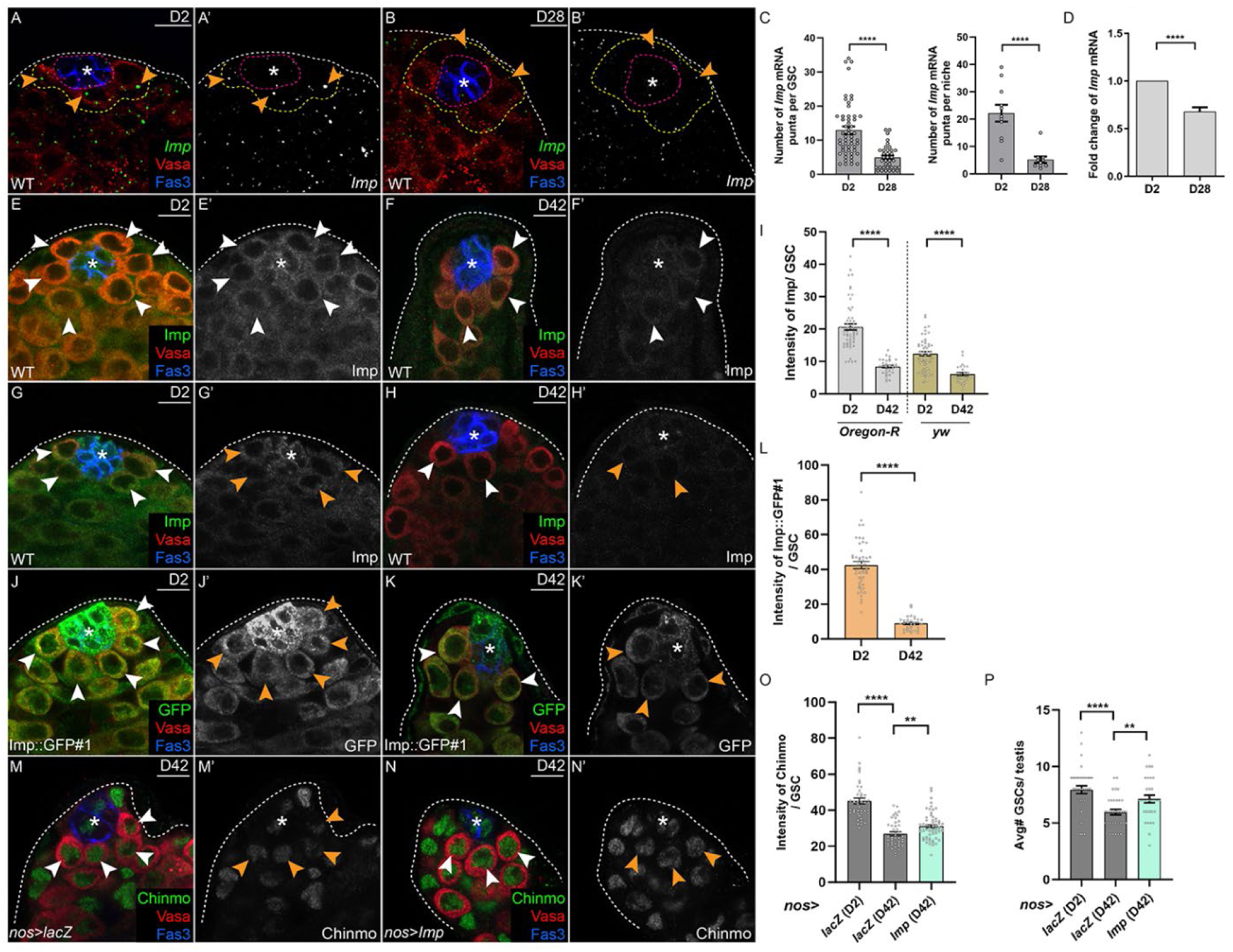
Imp positively regulates Chinmo protein expression in GSCs during aging. (A-C) Confocal images of RNA-FISH for *Imp* mRNA (green) in young (D2) testes (A) and middle-aged (D28) testes (B). The yellow and blue dashed lines outline GSCs and niche cells, respectively. Graph in C shows the average number of *Imp* mRNA puncta per GSC (left) and per niche cell (right). (D) qRT-PCR of *Imp* mRNA in D2 and D28 WT testes. (E–I) Confocal images of Imp protein (green) in WT D2 (E, G) and D42 (F, H) testes, quantified in I. (J-L) Confocal images of Imp::GFP protein trap #1 (green) in D2 (J) and D42 (K) testes, quantified in L. (M-O) Confocal images of Chinmo (green) in D42 *nos>lacZ* (M) or D42 *nos>Imp* (N) testes, quantified in O. (P) Graph showing the number of GSCs in D2 or D42 *nos>lacZ* (gray bars) or D42 *nos>Imp* (light blue bars) testes. In E, F, G, H, J, K, M, and N, arrowheads indicate GSCs, and asterisks indicate the niche. Vasa labels germ cells; Fas3 labels niche cells. Scale bars = 10 μm. In C, D, I, L, O, and P, error bars represent SEM. ** *P* ≤ 0.01; **** *P* ≤ 0.0001; n.s. indicates no significance, as assessed by Student’s t-test.

### Imp regulates age-dependent niche remodeling via Chinmo

Our prior work revealed that testes from aged D42 WT males had ectopic ECM accumulation around the niche (**Fig. 3A, C, E**) and that GSCs in these aged testes upregulated the ECM-binding protein Dg at the GSC-niche interface to remain in the altered niche (**Fig. 3F, H**) ^33^. We also demonstrated that the ectopic ECM and Dg in older testes were reversed by sustained Chinmo expression in GSCs ^33^. We now show that sustained Imp expression in GSCs throughout adulthood significantly suppressed the ectopic ECM (**Fig. 3B, D, E**) and ectopic Dg at the GSC-niche interface in aged testes (**Fig. 3G, H**).

**Figure 3.**
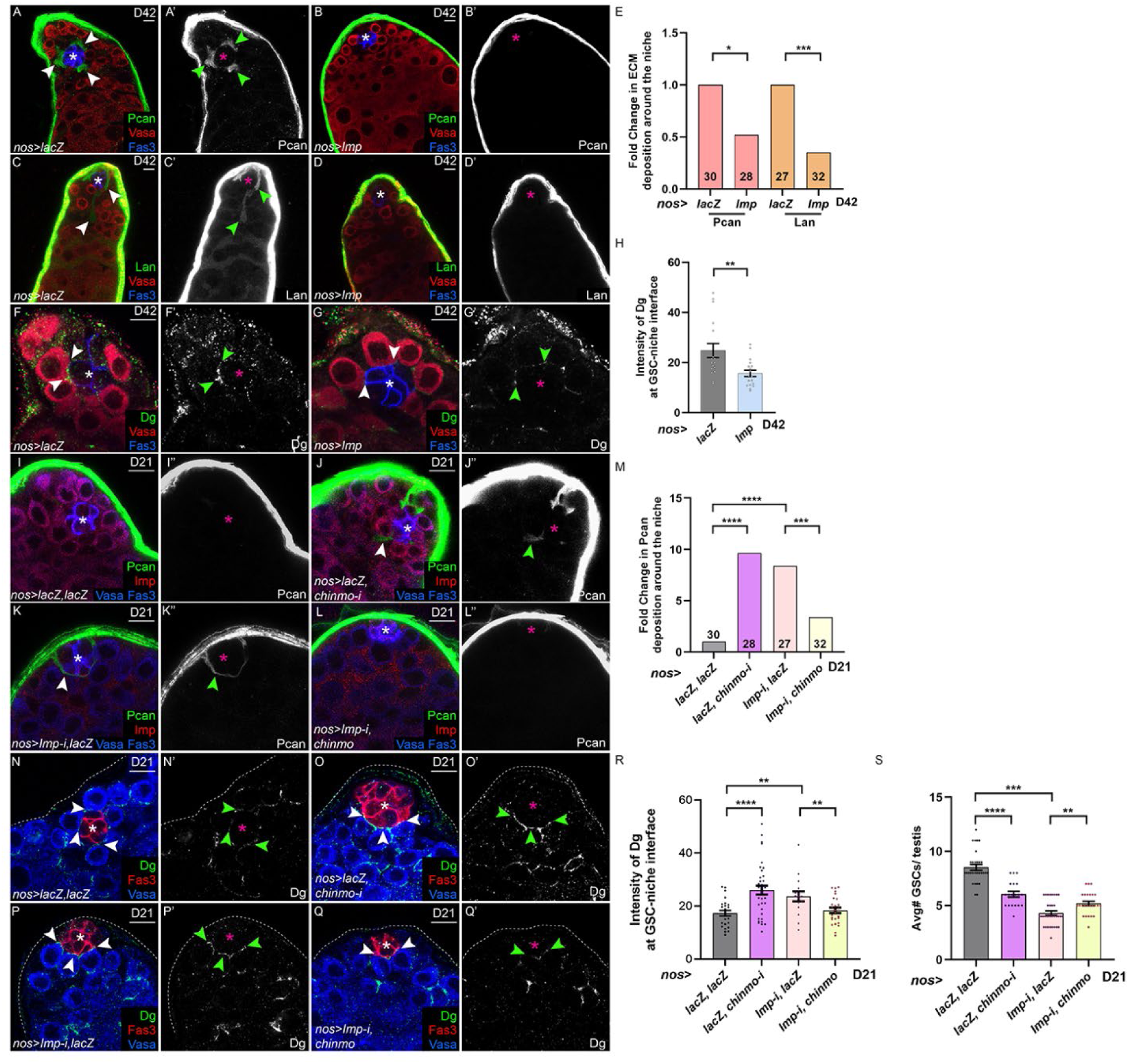
Imp controls aging of the testis stem cell niche via Chinmo. (A–H) Confocal images showing Pcan (green in A, B), Lan (green in C, D), or Dg (green in F, G) proteins in D42 *nos>lacZ* (A, C, F) or D42 *nos>Imp* (B, D, G) testes, quantified in E (for Pcan and Lan) and H (for Dg). Arrowheads in A, C mark ectopic Pcan or Lan surrounding the niche. Sample numbers for each genotype were indicated in each bar. In F, G, arrowheads show increased Dg at the GSC–niche interface in *nos>lacZ* (F) compared to *nos>Imp* (G). (I–L) Confocal images of Pcan (green) in D21 testes in *nos>lacZ, lacZ* (I), *nos>lacZ, chinmo-i* (J), *nos>Imp-i, lacZ* (K), and *nos>Imp-i, chinmo* (L). Arrowheads in (J, K) indicate ectopic Pcan around the niche. (M) Graph showing the fold change in Pcan surrounding the niche in D21 testes of the genotypes shown in (I–L). (N–R) Confocal images of Dg (green) in D21 testes of the genotypes shown in (I–L), quantified in R. Arrows indicate Dg the GSC–niche interface. (S) Graph showing that the average number of GSCs in the genotypes shown in (I–L). Asterisks in indicate the niche. Vasa labels germ cells; Fas3 labels niche cells. Scale bars = 10 μm. In H, R and S, error bars represent SEM. * *P* ≤ 0.05; ** *P* ≤ 0.01; *** *P* ≤ 0.001; **** *P* ≤ 0.0001 as assessed by Student’s t-test in H, R, S, and by χ^2^ test in E and M.

To determine whether the age-dependent effects of Imp occurred through Chinmo, we examined ECM deposition and Dg localization in GSCs that were depleted for *Imp* and that over-expressed Chinmo. We reasoned that if Chinmo functioned downstream of Imp, then sustained Chinmo will rescue the effects of Imp depletion. For this experiment, we chose a reproductive middle age time point of 21 days (D21) because few testes from control males (*nos>lacZ, lacZ*) have ectopic ECM in the testis lumen or ectopic Dg (**Fig. 3I, M, N, R**). Testes from D21 males with GSC depletion of *chinmo* (*nos>lacZ, chinmo-i*) or GSC depletion of Imp (*nos>Imp-i, lacZ*) have a significant increase in luminal ECM and in Dg at the GSC-niche interface (**Fig. 3J, K, M, O, P, R**); they also have a significant drop in the number of GSCs (**Fig. 3S**). Concomitant over-expression of Chinmo and depletion of Imp significantly reduced the ectopic ECM and Dg and significantly increased the average total number of GSCs (**Fig. 3L, M, Q-S**). These results indicate that Imp prevents stem cell niche aging through Chinmo.

### H negatively regulates Imp in GSCs

Our data indicate that aged GSCs have reduced *Imp* mRNA (**Fig. 2A-D**). Consistent with this, factors known to regulate *Imp* expression in other tissues (i.e., miRNAs, the RBP Syncrip (Syp), and Chinmo ^31,45,46^) did not impact Imp levels in male GSCs (**Fig. S4**). To identify factors that control *Imp* in GSCs, we performed an RNAi screen of transcriptional regulators known to genetically interact with *Imp* per Flybase and to be expressed in early germ cells per Fly Cell Atlas: the transcription factors Ets21C and p53, and the co-repressor H ^47–50^. We depleted these factors individually in GSCs during adulthood and examined Imp protein. Only germline depletion of H using two independent RNAi lines significantly altered Imp protein in GSCs (**Fig. S5A-G**). Since Imp protein levels increase upon germline H depletion, H is a negative regulator of Imp expression in GSCs.

### H expression in GSCs increases during aging

To assess whether H expression in GSCs is modulated during aging, we assessed the activity of the *H* gene using the *H^CR60128-TG4.1^*-GAL4 line crossed to *UAS-mCD8-GFP*, encoding a membrane-tethered GFP. We monitored *H* expression in young D2 males and aging D28 males because there is a significant decline in the average total number of GSCs in WT testes at D28 (**Fig. 4H**) ^33^. GSCs from young testes have low *H* expression as indicated by low levels of membrane GFP (**Fig. 4A**). By contrast, GSCs from D28 males have robust *H* expression (**Fig. 4B**) that was similar to the positive control of membrane GFP driven by *nos-Gal4* (**Fig. 4C**). We also found a significant increase in *H* transcripts and in protein during aging (**Fig. 4D-G**).

**Figure 4.**
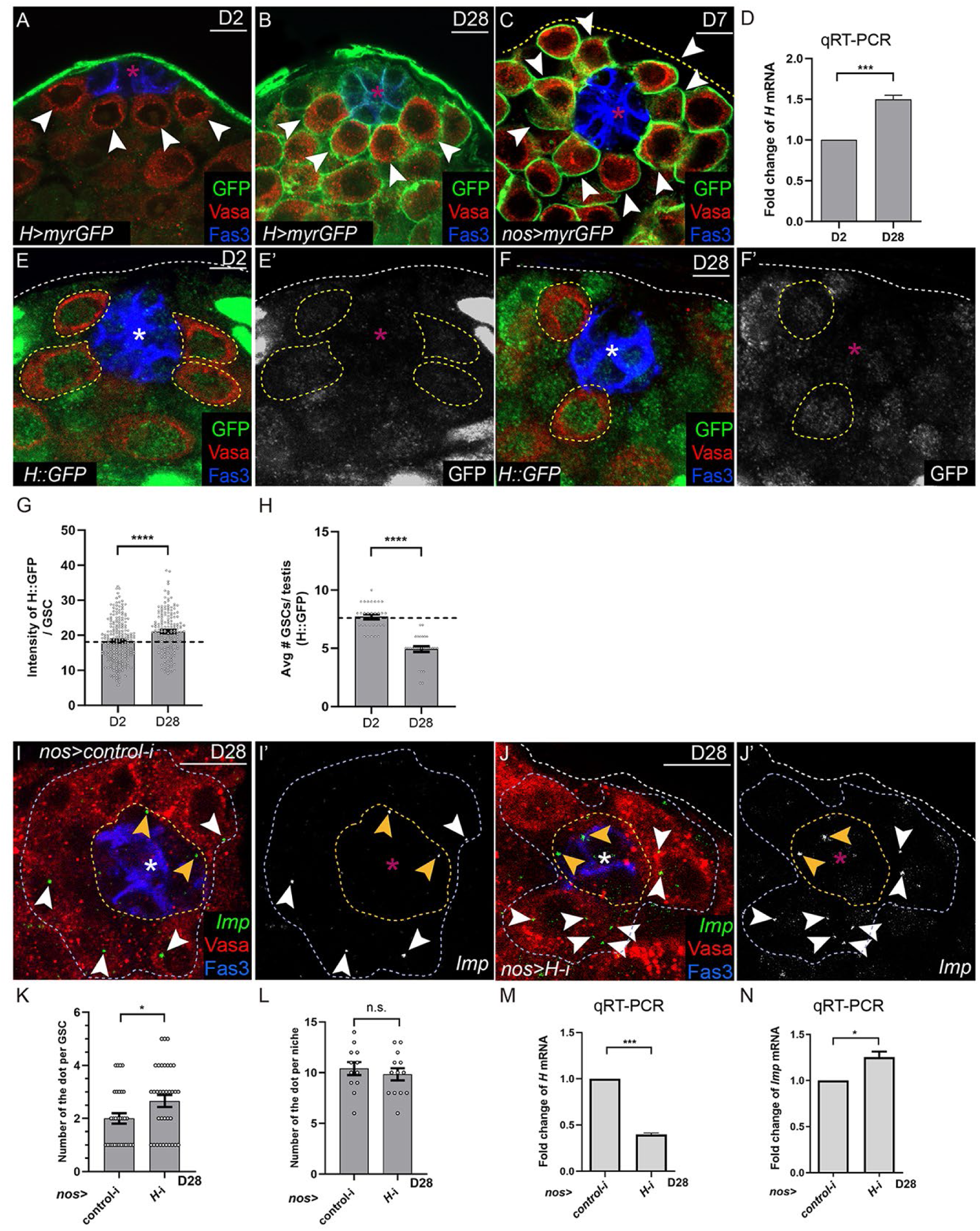
H is upregulated during aging and suppresses *Imp*. (A–C) Confocal images of myristoylated GFP (myrGFP, green) in D2 (A) and D28 (B) *H>myrGFP* testes, and *nos>myrGFP* testes at D7 (C). Arrowheads indicate GSCs. (D) Graph showing the relative fold change of *H* mRNA in D2 and D28 testes, measured by qRT-PCR. (E–H) Confocal images of H::GFP (green) in D2 (E) and D28 (F) testes, quantified in G. Graph in H shows the average number of GSCs in D2 and D28 H::GFP testes. Dashed lines in E, F indicate GSCs. (I–L) RNA-FISH for *Imp* mRNA (green) in D28 *nos>control-i* (I) or *nos>H-i* (J) testes, quantified in K for GSCs and in L for niche cells. Blue and yellow dashed lines circumscribes the niche and GSCs, respectively. Arrowheads indicate *Imp* mRNA puncta. (M, N) qRT-PCR analysis of *H* (M) or *Imp* (N) transcripts in D28 *nos>control-i* or *nos>H-i* testes. Asterisks in indicate the niche. Vasa labels germ cells; Fas3 labels niche cells. Scale bars = 10 μm. In G, H, K, L, M, and N, error bars represent SEM. * *P* ≤ 0.05; *** *P* ≤ 0.001; **** *P* ≤ 0.0001 as assessed by Student’s t-test.

### *Imp* is transcriptionally regulated by H in GSCs

Our results demonstrate that Imp and H exhibit opposite expression patterns in adult male GSCs. Since H can decrease transcription of target genes ^51^, we determined whether H could downregulate *Imp* transcripts. *Imp* mRNA is detected in GSCs in D28 control testes and is significantly increased in H-depleted GSCs at this time point (**Fig. 4I-K, M, N**). As expected, *Imp* levels in niche cells across the two genotypes were unchanged at D28 (**Fig. 4L**). These data indicate that H transcriptionally regulates *Imp* in GSCs during homeostasis.

### Self-renewal cues regulate aging of the testis stem cell niche

Because niche-derived soluble factors can decline with age ^31^, we investigated whether reduced levels of the BMPs Dpp and Gbb might underlie GSC aging. Dpp and Gbb are well-established self-renewal cues for GSC maintenance ^36,39–41^, but their role in testis stem cell niche aging is not known. While we observed robust expression of both proteins in the niche cells of young D2 testes, their levels were significantly diminished in D28 testes (**Fig. 5A–F**). We next asked whether maintaining high levels of these ligands during adulthood would suppress the age-dependent increase in H in GSCs. We overexpressed *UAS* transgenes encoding either Dpp::GFP or Gbb::GFP in niche cells with the niche-specific driver *upd*-*Gal4* and examined H::GFP in GSCs at D14 and D28. In both *upd>Dpp::GFP* and *upd>Gbb::GFP* testes, H::GFP was significantly reduced in GSCs compared to controls (**Fig. 5G-J**). We evaluated whether sustained BMP levels in niche cells would mitigate aging phenotypes at D28. In *upd>Dpp::GFP* and *upd>Gbb::GFP* testes at D28, Chinmo was significantly elevated in GSCs, while Dg was significantly reduced (**Fig. 5K, L**). Additionally, Pcan accumulation around the niche was substantially reduced, and there were significantly more GSCs compared relative to age-matched controls (**Fig. 5M, N**). Taken together, these findings indicate that H levels in GSCs are negatively regulated by niche-derived BMPs and that sustaining robust BMP expression in niche cells prevents aging-phenotypes.

**Figure 5.**
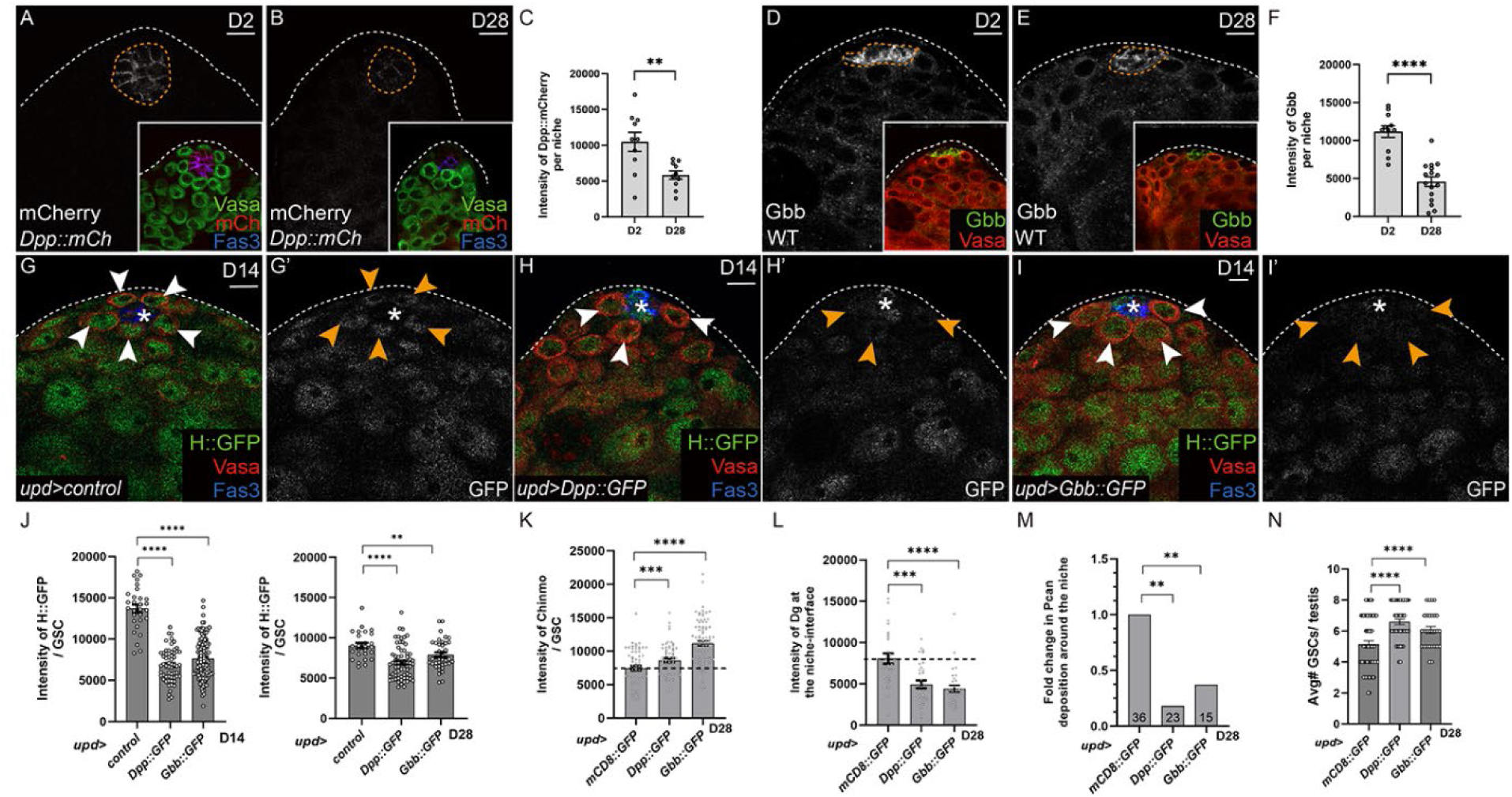
Overexpression of BMPs suppresses H and age-related phenotypes of the testis. (A-F) Confocal images of Dpp::mCherry (abbreviated Dpp::mCh, white, A and B) and Gbb (white, D and E) in D2 (A, D) and D28 (B, E) testes, quantified in C for Dpp::mCherry and in G for Gbb. Orange dashed lines indicate the level of Dpp::mCherry and Gbb in niche cells. (G–H) Confocal images of H::GFP (green) in D14 testes in *upd>+* (control, G), *upd>Dpp::GFP* (H), or *upd>Gbb::GFP* (I). (J) Graph showing intensity of H::GFP per GSC in D14 (left) vs D28 (right) testes. (K-N) Graph showing intensity of Chinmo (K) per GSC and Dg (L) at GSC-niche interface, the fold change in Pcan surrounding the niche (M), and the average total number of GSCs (N) in D28 testes in *upd>mCD8-GFP*, *upd>Dpp::GFP*, or *upd>Gbb::GFP*. Asterisks in indicate the niche. Vasa labels germ cells; Fas3 labels niche cells. Scale bars = 10 μm. In C, F, J, K, L and N, error bars represent SEM. * *P* ≤ 0.05; ** *P* ≤ 0.01; *** *P* ≤ 0.001; **** *P* ≤ 0.0001 as assessed by Student’s t-test for C, F, J, K, L and N, and by χ^2^ test in M.

### H acts in GSCs to promote aging of the testis stem cell niche

Since BMPs repress H expression in GSCs, we hypothesized that GSC depletion of H would ameliorate age-dependent phenotypes of the testis stem cell niche. Consistent with this model, GSC depletion of H for 28 days significantly increased Chinmo protein (**Fig. 6A-C**), significantly decreased ectopic Pcan in the testis lumen and ectopic Dg at the GSC-niche interface (**Fig. 6D-I**) and significantly restored the average total number of GSCs compared to the aged-match control (**Fig. 6J**). Similar results were observed with another independent *H*-RNAi line (**Fig. S5H-Q**). Furthermore, we found that H over-expression in GSCs accelerated aging of the testis stem cell niche. Over-expression of H using three independent *UAS* lines (*H^EY03696^* = *H^EY^, H^EP-211^* = *H^EP^, H^GS11776^* = *H^GS^*) in GSCs caused a significant decline in Imp and Chinmo proteins, resulting in significantly more luminal Pcan and Dg at the GSC-niche interface, and impaired GSC homeostasis (**Fig. 6K-W**). Thus, both loss and gain of function experiments indicate that H levels in GSCs regulate aging of the testis stem cell niche.

**Figure 6.**
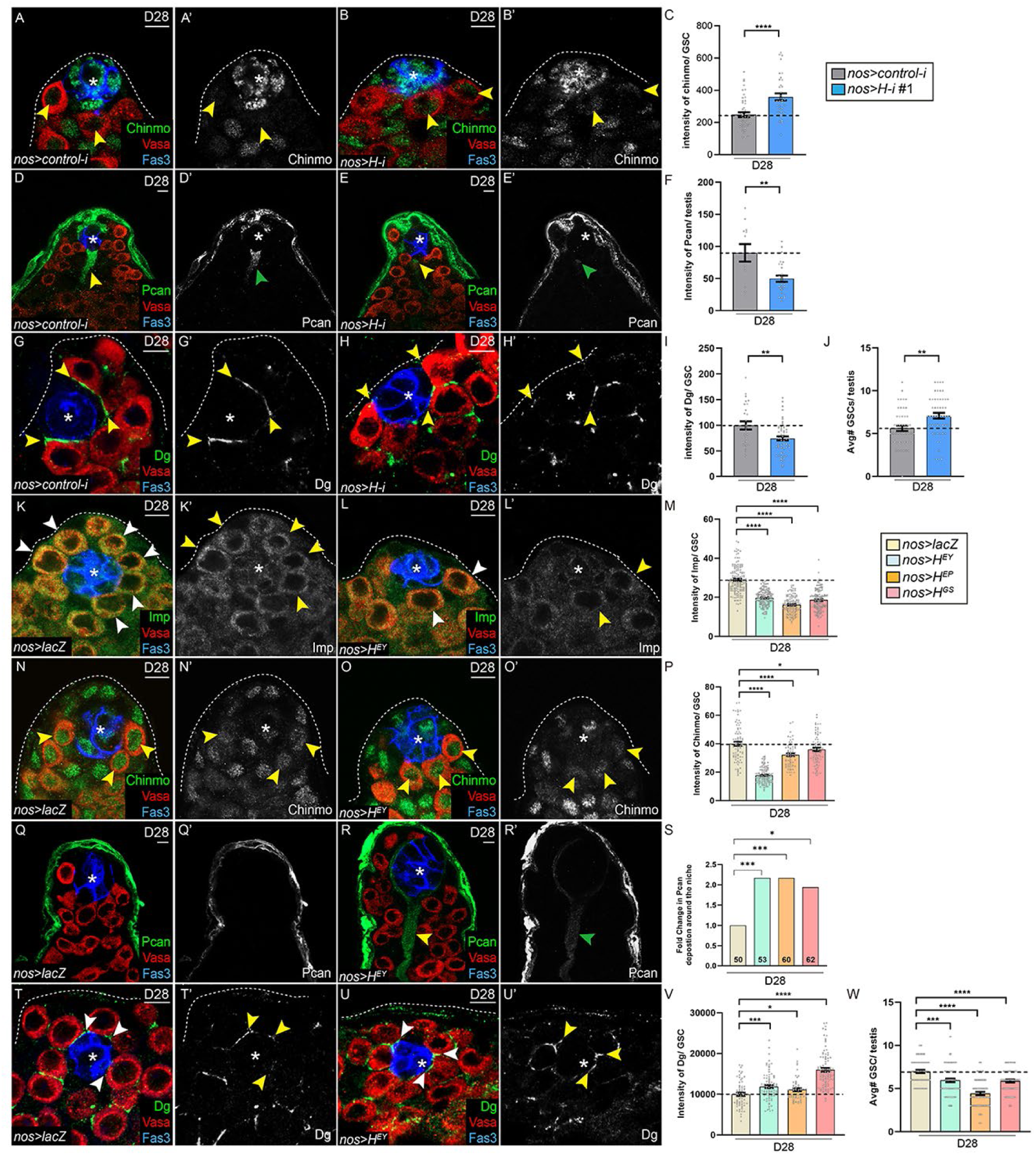
Hairless negatively regulates aging of the testis stem cell niche. (A-J) Confocal image of Chinmo (green, A, B), Pcan (green, D, E) or Dg (green, G, H) in D28 *nos>control-i* (A, D, G) or *nos>H-i* testes (B, E, H), quantified in C, F, I, respectively. Arrowheads indicate Chinmo in GSCs (A, B); ectopic Pcan surrounding the niche (D, E), or Dg at the GSC-niche interface (G, H). Graph in J shows the average number of GSCs in D28 indicated genotype testes. (K-W) Confocal image of Imp (green, K, L), Chinmo (green, N, O), Pcan (green, Q, R), or Dg (green, T, U) in D28 *nos>lacZ* (K, N, Q, T) or *nos>H^EY^* testes (L, O, R, U), quantified in M, P, S, V, respectively. Arrowheads indicate Imp in GSCs (K, L), Chinmo in GSCs (N, O), ectopic Pcan in the testis lumen (R), or Dg at the GSC-niche interface (T, U). Graph in W shows the average number of GSCs in D28 indicated genotype testes. Asterisks in indicate the niche. Vasa labels germ cells; Fas3 labels niche cells. Scale bar = 10 μM In C, F, I, J, M, P, V, and W, error bars represent SEM. * *P* ≤ 0.05; ** *P* ≤ 0.01; *** *P* ≤ 0.001; **** *P* ≤ 0.0001 as assessed by Student’s t-test in C, F, I, J, M, P, V, and W, and by χ^2^ test in S.

### H functions independently of Notch (N) signaling in GSCs

N target genes are repressed by a complex of co-repressors (H, Groucho and dCtBP) that recruit the transcription factor Suppressor of Hairless (Su(H)) ^51,52^. Although we observe low levels of Su(H)::GFP in GSCs, we did not detect any N transcriptional activity in the adult testis using established reporters, even when we depleted H from GSCs (**Fig. S6A-E, H-K**). As expected, N activity was observed in polar cells of the *Drosophila* ovary, indicating the efficacy of the N reporters in adult gonads (**Fig. S6F, G**) ^53^. Furthermore, ectopic activation of N using the dominant-active NICD transgene in GSCs for 28 days of adulthood did not alter Imp protein levels or the average total number of GSCs (**Fig. S6L-O**), which is consistent with our model that H has N-independent functions in male GSCs (**Fig. S6P**).

### *Imp*-deficient or H-overexpressing GSC clones dominate their niche by evicting non-mutant neighbors

We previously demonstrated that *chinmo*-mutant GSC clones had a competitive advantage over their non-mutant sibling GSCs by co-opting an aging mechanism (decline of Chinmo protein in GSCs during aging) ^33^. Next, we wanted to test whether *Imp*-mutant or H-over-expressing GSC clones exhibit similar behavior. *Imp* is located on the X chromosome and *Imp* mutations are lethal in males, precluding mosaic analysis. Instead, we induced MARCM GSC clones that over-expressed a validated *Imp-RNAi* transgene and GFP or control clones that expressed GFP ^54^. We also employed the GSC-specific *nos* flip-out methodology (*nos-FRT-STOP-SV40-polyA-FRT-gal4-VP16-nos 3′UTR*) ^55^ to generate *Imp*-mutant GSCs clones with a second validated RNAi transgene and or H-overexpressing clones with two independent *UAS* lines *H^EP^*and *H^GS^* (**Fig. 7A**). In both MARCM and *nos* flip-out techniques, the clones are GFP-positive and the non-mutant sibling GSCs are GFP-negative (**Figs. 7B-E**; **S7A,B**). As expected, *Imp*-*i* and H-overexpressing GSC clones have significantly lower Imp and Chinmo protein **(Figs. 7E, F, L, M**) and significantly increased ECM in the testis lumen (**Figs. 7G-K, O**; **S7C-E**) at 14 dpci compared to control clones. Additionally, H-overexpressing GSC clones have significantly higher Dg at the GSC-niche interface compared to control clones (**Fig. 7N**).

**Figure 7.**
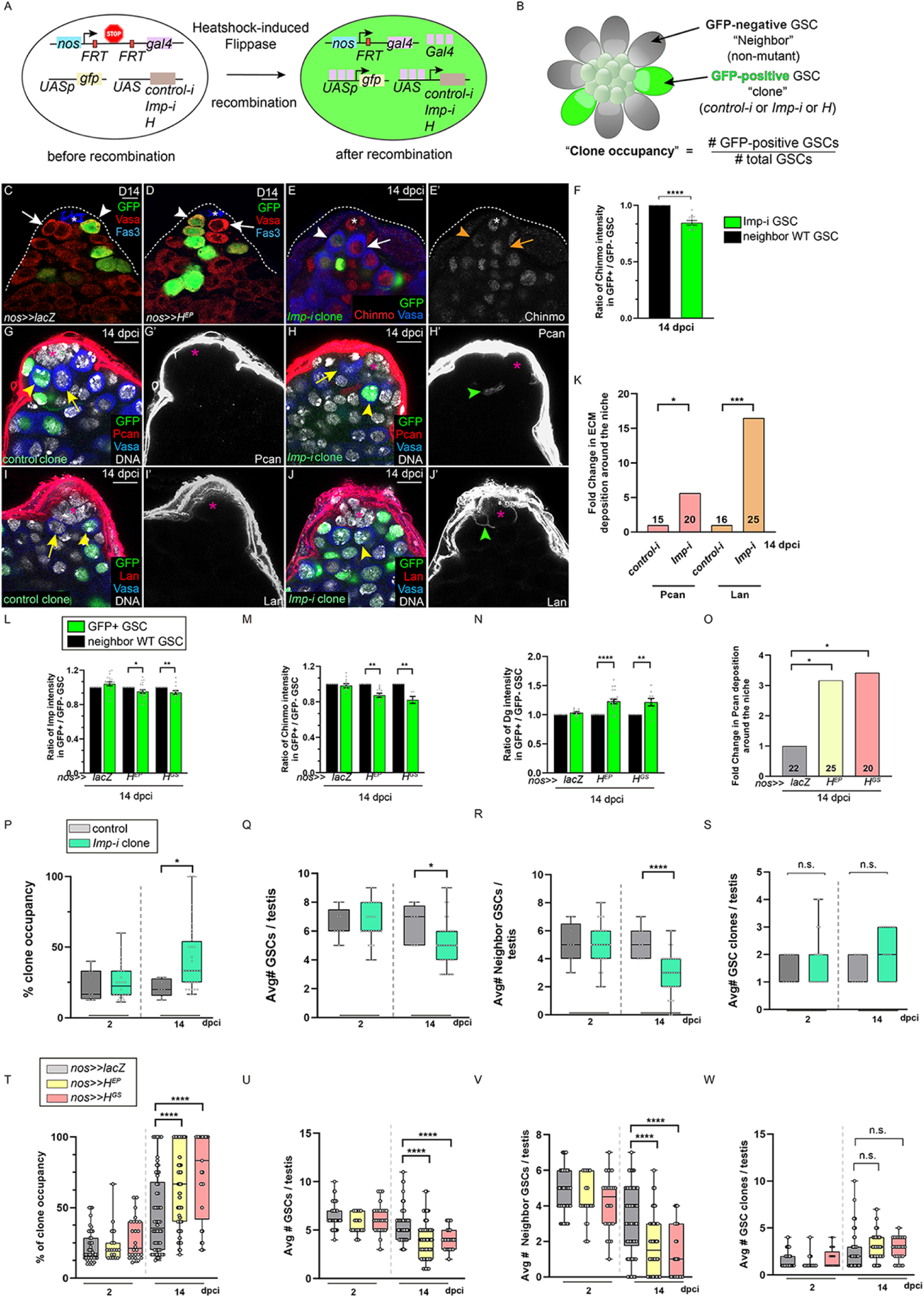
*Imp*-deficient and H-overexpressing GSC clones gain a competitive advantage for niche occupancy and evict non-mutant neighbors. (A) Schematic of clone induction using the *nos* flip-out technique. The *nos* promoter drives the expression of a cassette containing two *FRT* (Flippase Recognition Target) sites flanking a stop codon and the *gal4* coding sequence on the same chromosome. A *UASp* promoter controls *gfp*, and either *UAS-lacZ* (control) or *UAS-H* is also present in the GSC. Heat-shock induction of Flippase (FLP) excises the intervening sequence between the *FRT* sites, removing the stop codon and allowing Gal4 expression. As a result, Gal4 activates GFP and either LacZ or H from the *UASp* or *UAS* promoter, leading to positively-labeled clones. (B) Diagram illustrating the clone occupancy assay. In this testis, eight GSCs surround the niche. Two GSCs are clones (either control or *Imp^-/-^*) and are GFP-positive (green). The non-mutant neighbor GSCs are GFP-negative. The occupancy of each clone is calculated by dividing the number of GFP-positive GSCs by the total number of GSCs in each testis. In this example, the clone occupancy is 25% (2 GSC clones/8 total GSCs). (C, D) Confocal images of testes with GFP-positive *nos>>lacZ* GSC clones (C, arrowhead) or *nos>>H* GSC clones (D, arrowhead) at 14 days post-clone induction (dpci). Arrows indicate non-mutant neighboring GSCs. (E-F) Confocal image of a testis harboring a GFP-positive *Imp^-/-^* GSC clone (arrowhead) at 14 dpci. The arrow indicates a GFP-negative non-mutant GSC neighbor. Graph in F shows the ratio of Chinmo protein intensity in the GSC clone vs that in the non-mutant GSC for control clones (black bar) and *Imp*-mutant clones (green bar). (G-K) Confocal images of Pcan (red, G, H) and Lan (red, I, J) expression in the testis lumen with control GSC clones (G, I) or *Imp^-/-^* GSC clones (H, J), quantified in K. Green arrowheads mark ectopic Pcan (H) or ectopic Lan (J) near the niche. The total number of analyzed testes are indicated in each bar. (L-O) Graph showing the ratio of Imp (L), Chinmo (M) or Dg (N) protein at the niche-GSC interface in GSC clones (*i.e.*, *nos>>lacZ*, *nos>>H^EP^*, or *nos>H^GS^* GSC clones (green bar)) to neighbor GSCs (black bar) at 14 dpci. (O) Graph quantifying the fold change Pcan surrounding the niche when *nos*>>*lacZ* GSC clones (first bar)*, nos>>H^EP^* GSC clones (second bar), or nos>>*H^GS^* GSC clones (third bar) are present. The number of testes analyzed are indicated in each bar. (P-S) Box-and-whisker plots depicting (P) clone occupancy, (Q) average total number of GSCs, (R) average number of non-mutant GSC neighbors, and (S) average number of GSC clones in testes containing control (gray bars) or *Imp^-/-^* (green bars) GSC clones at 2 and 14 dpci. (T-W) Box and whisker plots showing clone occupancy (T), average total number of GSCs (U), average number of non-mutant GSC neighbors (V), and average numbers of GSC clones (W) in testes with control *nos*>>*lacZ* GSC clones (gray bars), with *nos*>>*H^EP^* GSC clones (yellow bar), or with *nos*>>*H^GS^*GSC clones (pink bar) at 2 and 14 dpci. Asterisks in indicate the niche. DAPI labels DNA; Vasa labels germ cells; Fas3 labels niche cells. Scale bars = 10 μm. In F, L-N, P-W, error bars represent SEM. * *P* ≤ 0.05, ** *P* ≤ 0.01, *** *P* ≤ 0.001, **** *P* ≤ 0.0001 by Student’s t-test for F, L-N, P-W and by χ^2^ test for K, O.

To examine GSC competitiveness, we quantified the percent of *Imp-i* and H-over-expressing clones composing the GSC pool, termed “clone occupancy” ^33^, at 2 and 14 dpci (**Fig. 7B**). At 2 dpci, *Imp-i* or H-overexpressing GSC clones and their appropriate control clones occupied the niche in equal proportions (**Fig. 7P**; **Table S2 #1** and **#3**; **Fig. 7T**; **Table S4 #1, #3** and **#5**). By 14 dpci, *Imp-i* or H-overexpressing GSC clones occupied a majority of the GSC pool compared to control clones (**Fig. 7P**; 44.38 ± 5.38% GSCs in *Imp-i* compared to 21.12 ± 1.98% GSCs in control, *P* ≤ 0.0199; see also **Table S2 #4** vs **#2**; **Fig. 7T**; 63.92 ± 2.9% GSCs in *H^EP^* and 73.6 ± 6.05% in *H^GS^* compared to 41.86 ± 3.67% GSCs in control, *P* ≤ 0.0001; see also **Table S4, #6** vs **#2** and **#4** vs **#2**).

We previously showed that *chinmo*-mutant GSCs cause a significant drop in the average total number of GSCs by evicting of their non-mutant GSC neighbors ^33^. Here, we find that *Imp-i* or H-over-expressing GSC clones similarly affect the GSC pool by causing a net loss in the average total number of GSCs at 14 dpci (**Fig. 7Q**; 5.06 ± 0.37 GSCs in *Imp-i*, compared to 6.50 ± 0.43 GSCs in control, *P* ≤ 0.02; see also **Table S2 #8** vs **#6**; **Fig. 7U**; 3.85 ± 0.19 GSCs in *H^EP^* and 4.08 ± 0.22 GSCs in *H^GS^* compared to 5.17 ± 0.19 GSCs in control, *P* ≤ 0.0001; see also **Table S4 #12** vs **#8** and **#10** vs **#8**). Additionally, by 14 dpci non-mutant neighbor GSCs were precipitously lost from the niche in the presence of *Imp-i* or H-overexpressing GSC clones but not control GSC clones (**Fig. 7R**; 2.91 ± 0.33 non-mutant neighbor GSCs in *Imp-i* compared to 5.13 ± 0.37 in control, *P* ≤ 0.0001; see also **Table S2 #12** vs **#10**; **Fig. 7V**; 1.67 ± 0.16 non-mutant neighbor GSCs in *H^EP^*and 1.22 ± 0.29 in *H^GS^* compared to 3.18 ± 0.25 in control, *P* ≤ 0.0001; see also **Table S4 #18** vs **#14, #16** vs **#14**). There was no statistical difference between the number of control, *Imp-i*, or H-overexpressing GSC clones at 14 dpci (**Fig. 7S**; 1.38 ± 0.17 versus 1.92 ± 0.30, control and *Imp-i* clones, respectively, *P* ≤ 0.1191; see also **Table S2 #16** vs **#14**; **Fig. 7W**; 2.42 ± 0.12 GSCs in *H^EP^* and 2.76± 0.24 in *H^GS^* compared to 2.04 ± 0.17 GSCs in control, *P* ≤ 0.25 and *P* ≤ 0.54, respectively; see also **Table S4 #24** vs **#20** and **#22** vs **#20**). These data indicate that control, *Imp-i* or H-overexpressing GSC clones expand to a similar extent, consistent our prior work on *chinmo-*mutant GSC clones. They also highlight that the competitiveness of *Imp-i* or H-overexpressing GSC clones derives from their non-autonomous effects on non-mutant neighbors. Additionally, we obtained similar results on all of these parameters using an additional, validated *Imp-i* transgene (**Fig. S7F-I**; **Tables S1, #11-14** and **S3**) ^28^. Taken together, these results indicate that competitive GSCs exert their competitive advantage by expelling neighbor GSCs..

## Discussion

### Aging regulates the BMP/H/Imp/Chinmo axis in GSC testis niche

Our study reveals how age-dependent changes in niche cells of the *Drosophila* testis drive aging of the niche microenvironment. We demonstrate that declining BMP levels in niche cells lead to the upregulation of the co-repressor H in GSCs. In turn, H represses expression of the RBP Imp, thereby reducing levels of the key aging and competition factor Chinmo. Age-dependent phenotypes are caused by reduced BMP in niche cells, increased H in GSCs or reduced Imp/Chinmo in GSC, ultimately causing a decline in the GSC pool. Moreover, clonal analyses indicate that GSCs lacking Imp or harboring robust H gain a competitive advantage in niche residency over WT GSC neighbors. Altogether, our data link the age-related decline in self-renewal cues to a cell-intrinsic H/Imp/Chinmo mechanism that drives GSC aging and competition.

Our results suggest that Chinmo levels in GSCs regulate competition among WT GSCs during aging, ultimately contributing to the significant decline in GSC numbers in aged testes. Although Chinmo levels decrease overall with age, there is substantial variability in Chinmo expression between individual GSCs (**Fig. 1D**). In a youthful environment, high Chinmo levels repress expression of Pcan and Dg. However, as GSCs age and Chinmo levels decline, Pcan and Dg become upregulated. Secreted Pcan accumulates around the niche, forming a progressively expanding “moat” that alters the niche architecture. Within this altered environment, GSCs with the lowest Chinmo—and consequently the highest Dg—are best equipped to remain in the niche. In contrast, GSCs with relatively higher Chinmo and lower Dg are displaced and differentiate. Thus, our findings suggest that among WT GSCs, those with the lowest Chinmo levels “win” the competition for niche occupancy during aging.

### BMP production declines during aging

Although the JAK-STAT ligand Upd also declines with age in the testis niche, Chinmo is not regulated by STAT signaling in GSCs (**Fig. S1J-M**) ^30,31^. Instead, we show that Chinmo is positively regulated by the level BMPs in niche cells. Our data now provide direct evidence that BMP production in testicular niche cells wanes over time, and this local reduction is sufficient to cause age-dependent phenotypes in the niche microenvironment and in GSCs via Chinmo. Thus, BMPs appear to be the principal extrinsic cues orchestrating testis aging. Interestingly, a similar role for niche-derived BMPs has been shown for aging of the *Drosophila* ovary ^56^. BMP signaling is highly conserved, and studies in vertebrate models have demonstrated that significant alterations in BMP signaling regulate aging of several tissues ^57–59^.

We find that overexpression of either Gbb or Dpp in testicular niche cells throughout adulthood suppresses age-related changes in H, Chinmo, Pcan, Dg, and in GSC homeostasis. Thus, both ligands can mitigate niche deterioration when artificially elevated (**Fig. 5K–N**). Prior studies indicate that Gbb is more dominant under normal conditions, as *gbb* loss-of-function mutants display severe GSC loss, whereas *dpp* hypomorphic mutants are less affected ^39^. Further work is needed to determine whether Gbb and Dpp act as homodimers or heterodimers to regulate H, Imp, and Chinmo in aging GSCs.

### H is a Notch-independent regulator of aging

We found that H exerts a Notch-independent effect on adult GSCs by repressing *Imp* transcription. Although H is best known for antagonizing the Notch pathway, we did not detect changes in the expression of Notch transcriptional reporters when manipulating H in GSCs (**Fig. S6H-K**). Prior work has shown additional roles for H beyond direct Notch inhibition, including modulation of EGFR signaling to promote vein development and regulation of apoptosis or cell growth through diverse pathways ^47,60^. Our findings place H downstream of BMP signaling, as age-associated increases in H reduce Chinmo levels via repression of Imp. This indicates a broader function for H in stem cell competency and aging that extends beyond Notch. Several questions remain. For instance, it is not yet clear whether a specific threshold of H is needed to repress *Imp* or precisely how BMPs regulate H. Mothers against decapentaplegic (Mad) is the best-known transcriptional effector of BMP signaling in *Drosophila*: it becomes phosphorylated by BMP receptors upon BMP ligand binding on the GSC and subsequently represses *bam*, a key differentiation gene in female GSCs ^61^. Investigating potential binding sites for the Mad within the regulatory elements of *H* may clarify how external signals converge on the *H* locus to control GSC aging and competitive dominance. It is tempting to speculate that transcriptional changes in observed aged tissues might be caused by age-related increases in co-repressors ^2^.

### Parallels with mammalian Imp orthologs

Our findings parallel those from a recent study on the mammalian ortholog of Imp, IGF2BP1/2/3, in HSC aging ^62^. In mice, IGF2BP2 is highly expressed in young HSCs but becomes significantly downregulated with age, contributing to reduced stem cell function, decreased metabolism-related gene expression, and age-related phenotypes. These parallels in Imp/IGF2BP function across evolutionarily distant systems underscore the conserved role of this family of proteins in stem cell maintenance and the aging process. However, whereas the mammalian study highlights a predominantly intrinsic role of IGF2BP2 in HSC aging, our work shows how niche-dependent factors regulate Imp in *Drosophila* GSCs. This suggests that environmental cues influence Imp/IGF2BP activity in stem cells. Given the well-documented effects of aging on niches in both invertebrates and vertebrates, it is plausible that extrinsic regulatory pathways also control IGF2BP2 levels or function in mammalian HSCs. Unraveling these mechanisms could clarify how systemic cues contribute to stem cell aging in higher organisms.

### Limitations of the study

While our data suggest that Dpp and Gbb produced by niche cells are sufficient to modulate H expression in GSCs, we cannot exclude the possibility that additional cues from other nearby cells, such as CySCs that also secrete Dpp or Gbb ^36^ and contribute to regulating H in GSCs. Further tissue-specific knockdown or over-expression studies would be necessary to clarify this point. In addition, we cannot conclude whether Dpp and Gbb form homodimers or heterodimers when activating BMP signaling in GSCs. Determining the precise ligand composition *in vivo* would require endogenously tagged alleles in mutant backgrounds designed to distinguish between BMP ligand dimeric forms. Another limitation is the temporal resolution of our experiments. While we examined changes at certain age intervals, the dynamic progression of niche remodeling and GSC competition between these time points is not fully captured. A more continuous sampling of various time points during adulthood using long-term *ex vivo* live images would provide a clearer view of the intermediate steps in these processes.

## RESOURCE AVAILABILITY

### Lead contact

Further information and requests for resources and reagents should be directed to and will be fulfilled by the Lead Contact, Chen Yuan Tseng.

### Materials availability

Materials used in this study are available upon request to the lead contact, Chen-Yuan Tseng

### Data and code availability

- Any additional information required to reanalyze the data reported in this paper is available from the lead contact upon request.
- All data reported in this paper will be shared by the lead contact upon request.

## Acknowledgments

We thank all members of the Bach and Tseng laboratories for their insightful discussions and technical support. We are grateful to Dr. Jeremy J.-W. Chen (College of Medicine, National Chung Hsing University) for assistance with the confocal microscope (LSM980), and we thank the Instrument Center at National Chung Hsing University for help with the LSM900 confocal microscope. We thank P. Rangan, N. Sokol, T. Volk, S. Baumgartner, L. Weaver, C. Desplan, T. Lee, S. Bray, Bloomington *Drosophila* Stock Center (BDSC), Vienna *Drosophila* Resource Center (VDRC) and Kyoto *Drosophila* Stock Center for antibodies and fly stocks. The BDSC is supported by a grant from the Office of the Director of the NIH (P40OD018537). We are grateful to FlyBase, supported by a grant from the National Human Genome Research Institute at NIH (U41 HG000739). Work in the Bach lab is supported by grants from the NIH (R01GM085075, R35GM156624-01, R03HD090422). Work in the Tseng lab is supported by the Ministry of Science and Technology of Taiwan (MOST 111-2311-B-005-011-MY3), Taiwan and National Chung Hsing University and Chun Shan Medical University (NCHU-CSMU 11204) and start-up funds from the Institute of Life Science, NCHU.

## Author contributions

Conceptualization, C.Y.T. and E.A.B.; Investigation, Y.Z, Y.C.L., Y.T.W., P.K.C., S.L.C., L.S.H., H.J.H. and C.Y.T;. Manuscript writing. C.Y.T. and E.A.B.; Funding, C.Y.T. and E.A.B.; Supervision, C.Y.T. and E.A.B.

## STAR★Methods

### Key Resource Table

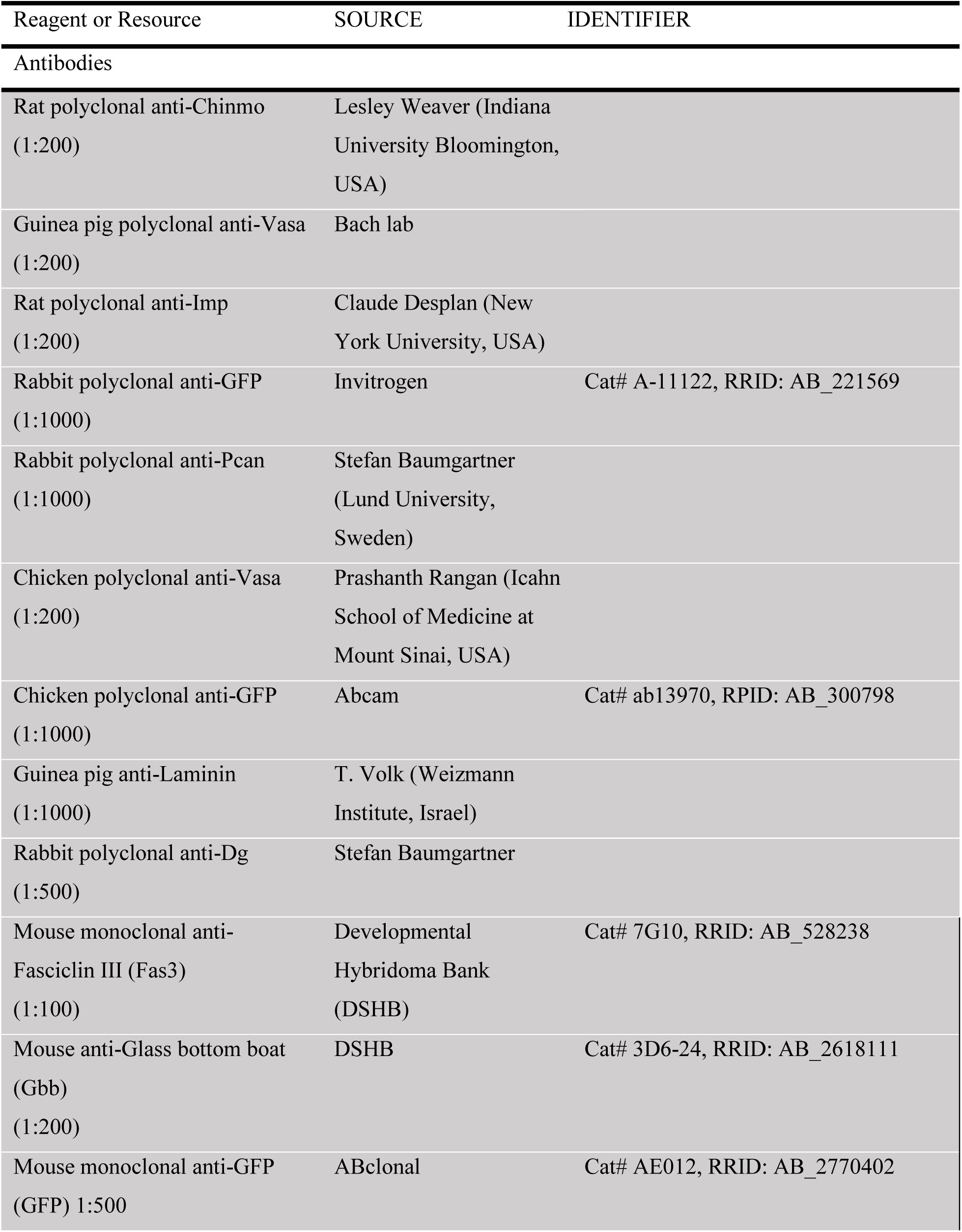
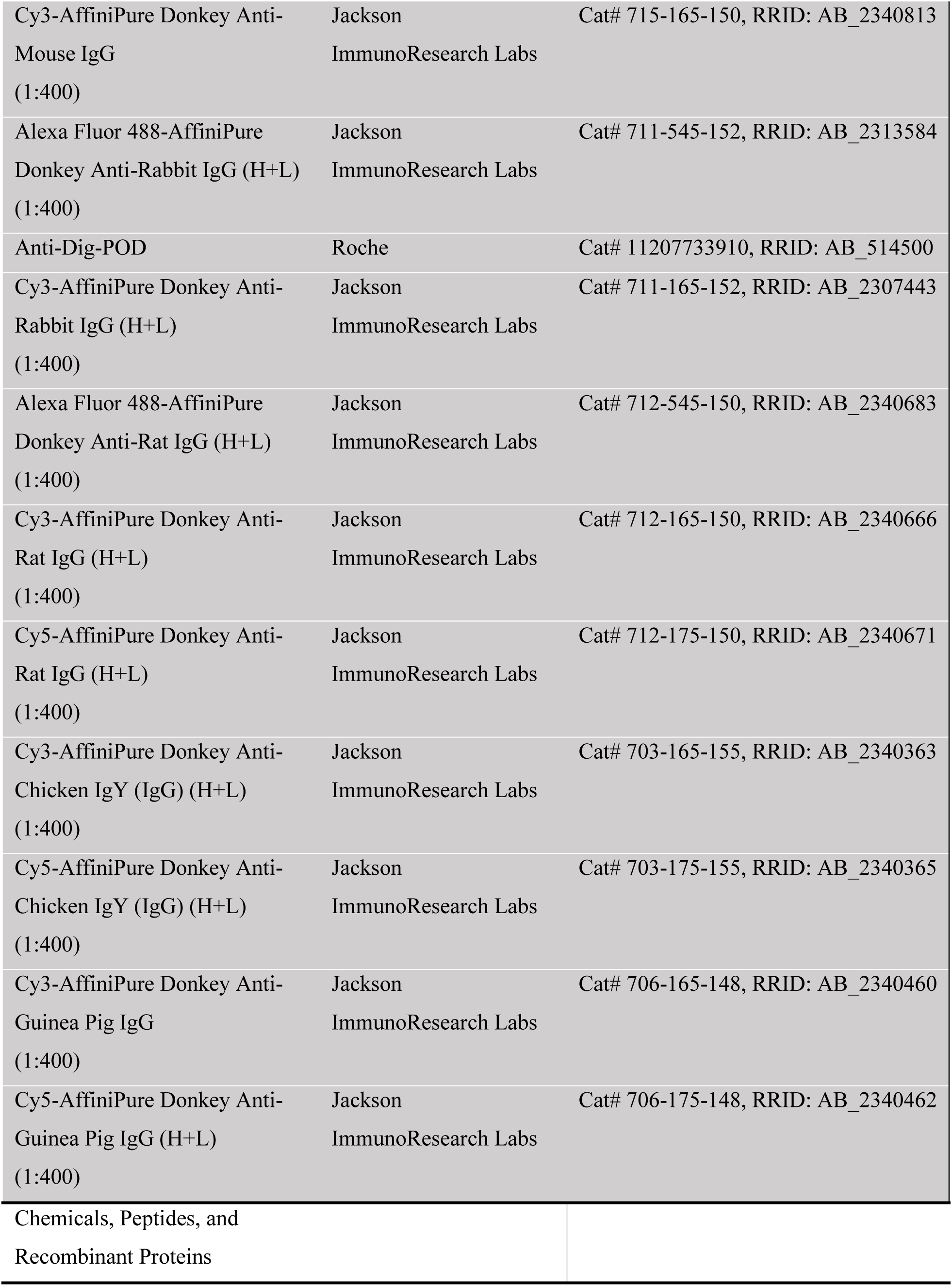
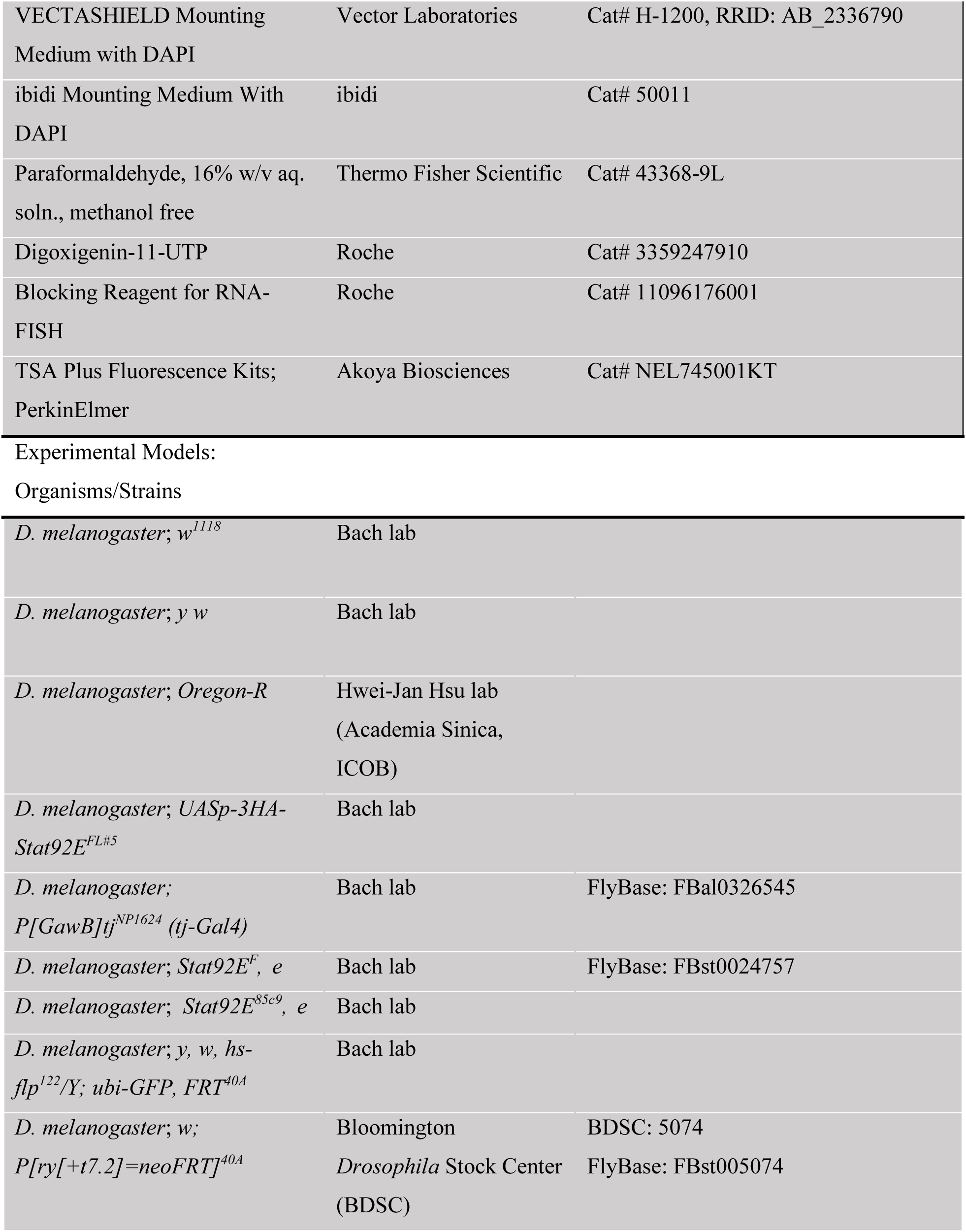
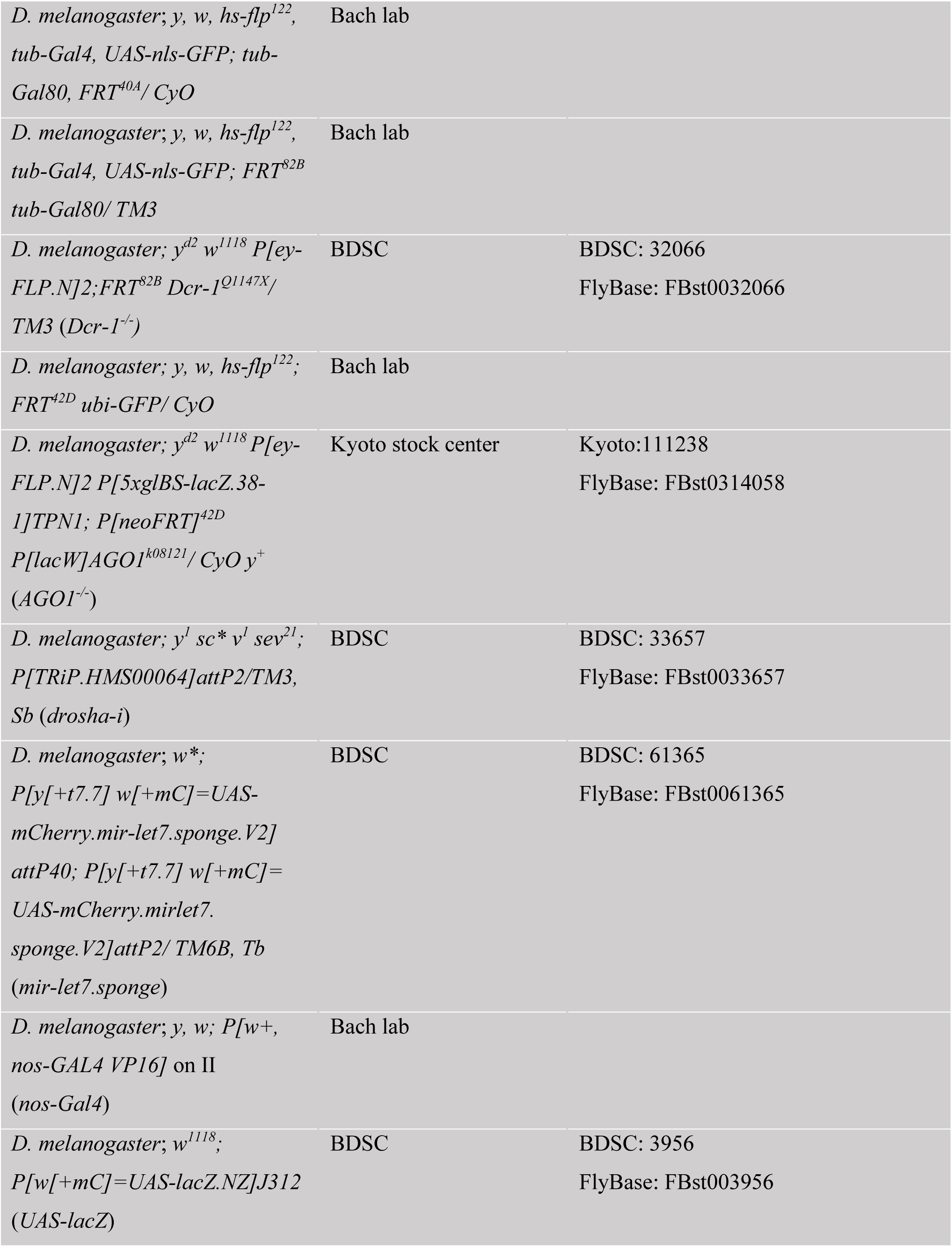
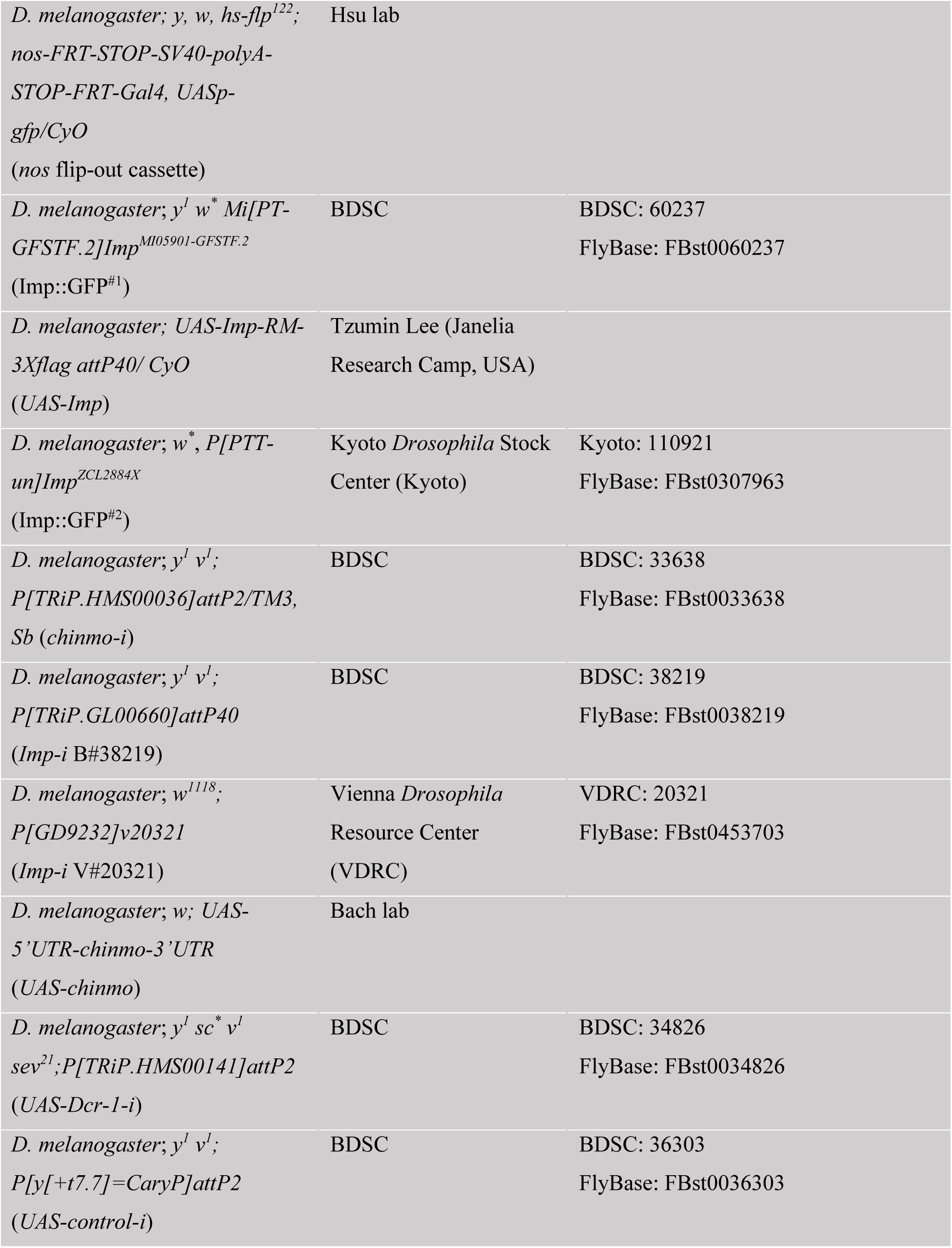
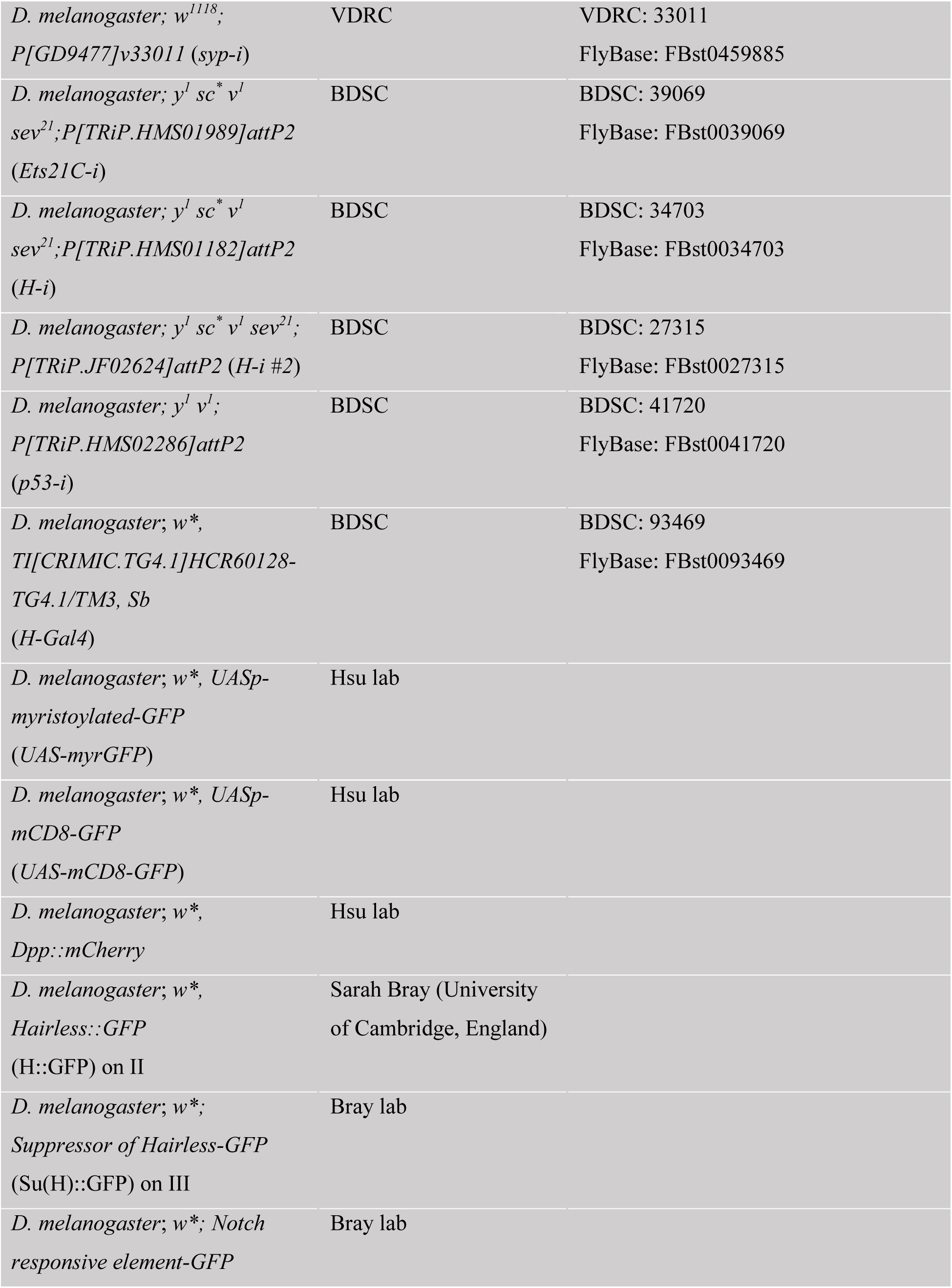
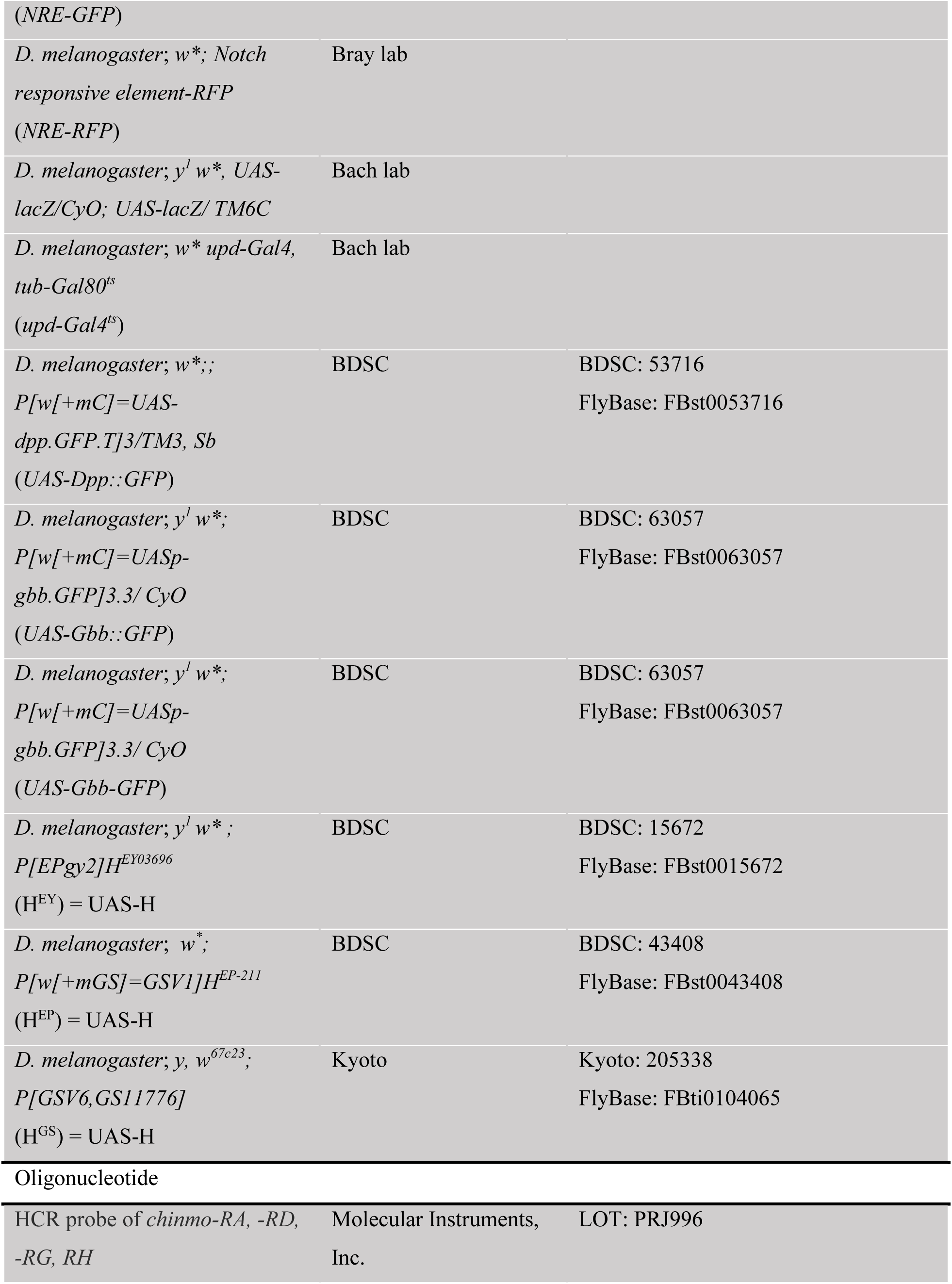
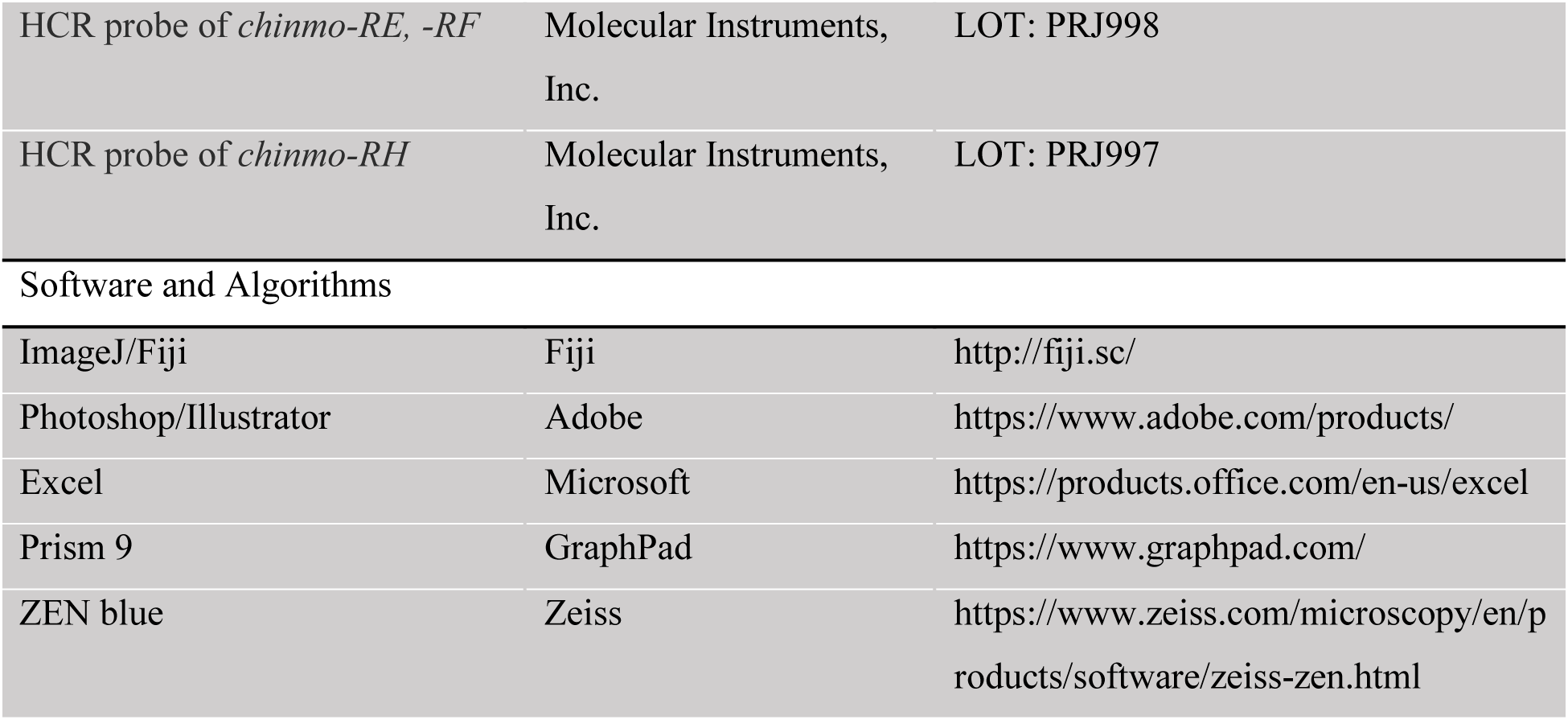

| Reagent or Resource | SOURCE | IDENTIFIER |
| --- | --- | --- |
| Antibodies |  |  |
| Rat polyclonal anti-Chinmo (1:200) | Lesley Weaver (Indiana University Bloomington, USA) |  |
| Guinea pig polyclonal anti-Vasa (1:200) | Bach lab |  |
| Rat polyclonal anti-Imp (1:200) | Claude Desplan (New York University, USA) |  |
| Rabbit polyclonal anti-GFP (1:1000) | Invitrogen | Cat# A-11122, RRID: AB_221569 |
| Rabbit polyclonal anti-Pcan (1:1000) | Stefan Baumgartner (Lund University, Sweden) |  |
| Chicken polyclonal anti-Vasa (1:200) | Prashanth Rangan (Icahn School of Medicine at Mount Sinai, USA) |  |
| Chicken polyclonal anti-GFP (1:1000) | Abcam | Cat# ab13970, RPID: AB_300798 |
| Guinea pig anti-Laminin (1:1000) | T. Volk (Weizmann Institute, Israel) |  |
| Rabbit polyclonal anti-Dg (1:500) | Stefan Baumgartner |  |
| Mouse monoclonal anti-Fasciclin III (Fas3) (1:100) | Developmental Hybridoma Bank (DSHB) | Cat# 7G10, RRID: AB_528238 |
| Mouse anti-Glass bottom boat (Gbb) (1:200) | DSHB | Cat# 3D6-24, RRID: AB_2618111 |
| Mouse monoclonal anti-GFP (GFP) 1:500 | ABclonal | Cat# AE012, RRID: AB_2770402 |
| Cy3-AffiniPure Donkey Anti-Mouse IgG (1:400) | Jackson ImmunoResearch Labs | Cat# 715-165-150, RRID: AB_2340813 |
| Alexa Fluor 488-AffiniPure Donkey Anti-Rabbit IgG (H+L) (1:400) | Jackson ImmunoResearch Labs | Cat# 711-545-152, RRID: AB_2313584 |
| Anti-Dig-POD | Roche | Cat# 11207733910, RRID: AB_514500 |
| Cy3-AffiniPure Donkey Anti-Rabbit IgG (H+L) (1:400) | Jackson ImmunoResearch Labs | Cat# 711-165-152, RRID: AB_2307443 |
| Alexa Fluor 488-AffiniPure Donkey Anti-Rat IgG (H+L) (1:400) | Jackson ImmunoResearch Labs | Cat# 712-545-150, RRID: AB_2340683 |
| Cy3-AffiniPure Donkey Anti-Rat IgG (H+L) (1:400) | Jackson ImmunoResearch Labs | Cat# 712-165-150, RRID: AB_2340666 |
| Cy5-AffiniPure Donkey Anti-Rat IgG (H+L) (1:400) | Jackson ImmunoResearch Labs | Cat# 712-175-150, RRID: AB_2340671 |
| Cy3-AffiniPure Donkey Anti-Chicken IgY (IgG) (H+L) (1:400) | Jackson ImmunoResearch Labs | Cat# 703-165-155, RRID: AB_2340363 |
| Cy5-AffiniPure Donkey Anti-Chicken IgY (IgG) (H+L) (1:400) | Jackson ImmunoResearch Labs | Cat# 703-175-155, RRID: AB_2340365 |
| Cy3-AffiniPure Donkey Anti-Guinea Pig IgG (1:400) | Jackson ImmunoResearch Labs | Cat# 706-165-148, RRID: AB_2340460 |
| Cy5-AffiniPure Donkey Anti-Guinea Pig IgG (H+L) (1:400) | Jackson ImmunoResearch Labs | Cat# 706-175-148, RRID: AB_2340462 |
| Chemicals, Peptides, and Recombinant Proteins |  |  |
| VECTASHIELD Mounting Medium with DAPI | Vector Laboratories | Cat# H-1200, RRID: AB_2336790 |
| ibidi Mounting Medium With DAPI | ibidi | Cat# 50011 |
| Paraformaldehyde, 16% w/v aq. soln., methanol free | Thermo Fisher Scientific | Cat# 43368-9L |
| Digoxigenin-11-UTP | Roche | Cat# 3359247910 |
| Blocking Reagent for RNA-FISH | Roche | Cat# 11096176001 |
| TSA Plus Fluorescence Kits; PerkinElmer | Akoya Biosciences | Cat# NEL745001KT |
| Organisms/Strains |  |  |
| *D. melanogaster*; *w*<sup>1118</sup> | Bach lab |  |
| *D. melanogaster*; *y w* | Bach lab |  |
| *D. melanogaster*; *Oregon-R* | Hwei-Jan Hsu lab<br>(Academia Sinica, ICOB) |  |
| *D. melanogaster*; *UASp-3HA-Stat92E*<sup>FL#5</sup> | Bach lab |  |
| *D. melanogaster*; *P[GawB]tj*<sup>NP1624</sup> (*tj-Gal4*) | Bach lab | FlyBase: FBal0326545 |
| *D. melanogaster*; *Stat92E*<sup>F</sup>, *e* | Bach lab | FlyBase: FBst0024757 |
| *D. melanogaster*; *Stat92E*<sup>85c9</sup>, *e* | Bach lab |  |
| *D. melanogaster*; *y, w, hs-flp*<sup>122</sup>/*Y*; *ubi-GFP, FRT*<sup>40A</sup> | Bach lab |  |
| *D. melanogaster*; *w*; *P[ry[+t7.2]=neoFRT]*<sup>40A</sup> | Bloomington<br>*Drosophila* Stock Center<br>(BDSC) | BDSC: 5074<br>FlyBase: FBst005074 |
| *D. melanogaster*; y, w, *hs-flp*<sup>122</sup>,<br>*tub-Gal4*, *UAS-nls-GFP*; *tub-Gal80*, *FRT*<sup>40A</sup>/ *CyO* | Bach lab |  |
| *D. melanogaster*; y, w, *hs-flp*<sup>122</sup>,<br>*tub-Gal4*, *UAS-nls-GFP*; *FRT*<sup>82B</sup><br>*tub-Gal80*/ *TM3* | Bach lab |  |
| *D. melanogaster*; y<sup>d2</sup> w<sup>1118</sup> *P[ey-FLP.N]2*; *FRT*<sup>82B</sup> *Dcr-1*<sup>Q1147X</sup>/<br>*TM3* (*Dcr-1*<sup>-/-</sup>) | BDSC | BDSC: 32066<br>FlyBase: FBst0032066 |
| *D. melanogaster*; y, w, *hs-flp*<sup>122</sup>;<br>*FRT*<sup>42D</sup> *ubi-GFP*/ *CyO* | Bach lab |  |
| *D. melanogaster*; y<sup>d2</sup> w<sup>1118</sup> *P[ey-FLP.N]2* *P[5xgls-lacZ.38-1]TPN1*; *P[neoFRT]*<sup>42D</sup><br>*P[lacW]AGO1*<sup>k08121</sup>/ *CyO* y<sup>+</sup><br>(*AGO1*<sup>-/-</sup>) | Kyoto stock center | Kyoto:111238<br>FlyBase: FBst0314058 |
| *D. melanogaster*; y<sup>1</sup> *sc* * v<sup>1</sup> *sev* <sup>21</sup>;<br>*P[TRiP.HMS00064]attP2/TM3*,<br>*Sb* (*drosha-i*) | BDSC | BDSC: 33657<br>FlyBase: FBst0033657 |
| *D. melanogaster*; w*;<br>*P[y[+t7.7] w[+mC]=UAS-mCherry.mir-let7.sponge.V2]*<br>*attP40*; *P[y[+t7.7] w[+mC]=UAS-mCherry.mirlet7.sponge.V2]attP2/ TM6B, Tb*<br>(*mir-let7.sponge*) | BDSC | BDSC: 61365<br>FlyBase: FBst0061365 |
| *D. melanogaster*; y, w; *P[w+, nos-GAL4 VP16]* on II<br>(*nos-Gal4*) | Bach lab |  |
| *D. melanogaster*; w<sup>1118</sup>;<br>*P[w[+mC]=UAS-lacZ.NZ]J312*<br>(*UAS-lacZ*) | BDSC | BDSC: 3956<br>FlyBase: FBst003956 |
| *D. melanogaster*; *y, w, hs-flp<sup>122</sup>; nos-FRT-STOP-SV40-polyA-STOP-FRT-Gal4, UASp-gfp/CyO*<br>(*nos* flip-out cassette) | Hsu lab |  |
| *D. melanogaster*; *y<sup>l</sup> w<sup>\*</sup> Mi[PT-GFSTF.2]Imp<sup>MI05901-GFSTF.2</sup>*<br>(Imp::GFP<sup>#1</sup>) | BDSC | BDSC: 60237<br>FlyBase: FBst0060237 |
| *D. melanogaster*; *UAS-Imp-RM-3Xflag attP40/ CyO*<br>(*UAS-Imp*) | Tzumin Lee (Janelia Research Camp, USA) |  |
| *D. melanogaster*; *w<sup>\*</sup>, P[PTT-un]Imp<sup>ZCL2884X</sup>*<br>(Imp::GFP<sup>#2</sup>) | Kyoto *Drosophila* Stock Center (Kyoto) | Kyoto: 110921<br>FlyBase: FBst0307963 |
| *D. melanogaster*; *y<sup>l</sup> v<sup>l</sup>; P[TRiP.HMS00036]attP2/TM3, Sb (chinmo-i)* | BDSC | BDSC: 33638<br>FlyBase: FBst0033638 |
| *D. melanogaster*; *y<sup>l</sup> v<sup>l</sup>; P[TRiP.GL00660]attP40*<br>(*Imp-i* B#38219) | BDSC | BDSC: 38219<br>FlyBase: FBst0038219 |
| *D. melanogaster*; *w<sup>1118</sup>; P[GD9232]v20321*<br>(*Imp-i* V#20321) | Vienna *Drosophila* Resource Center (VDRC) | VDRC: 20321<br>FlyBase: FBst0453703 |
| *D. melanogaster*; *w; UAS-5'UTR-chinmo-3'UTR*<br>(*UAS-chinmo*) | Bach lab |  |
| *D. melanogaster*; *y<sup>l</sup> sc<sup>\*</sup> v<sup>l</sup> sev<sup>21</sup>; P[TRiP.HMS00141]attP2*<br>(*UAS-Dcr-I-i*) | BDSC | BDSC: 34826<br>FlyBase: FBst0034826 |
| *D. melanogaster*; *y<sup>l</sup> v<sup>l</sup>; P[y[+t7.7]=CaryP]attP2*<br>(*UAS-control-i*) | BDSC | BDSC: 36303<br>FlyBase: FBst0036303 |
| *D. melanogaster*; $w^{1118}$;<br>*P*[GD9477]v33011 (*syp-i*) | VDRC | VDRC: 33011<br>FlyBase: FBst0459885 |
| *D. melanogaster*; $y^l sc^* v^l$<br>*sev*<sup>21</sup>; *P*[TRiP.HMS01989]*attP2*<br>(*Ets21C-i*) | BDSC | BDSC: 39069<br>FlyBase: FBst0039069 |
| *D. melanogaster*; $y^l sc^* v^l$<br>*sev*<sup>21</sup>; *P*[TRiP.HMS01182]*attP2*<br>(*H-i*) | BDSC | BDSC: 34703<br>FlyBase: FBst0034703 |
| *D. melanogaster*; $y^l sc^* v^l sev21;P[TRiP.JF02624]attP2 (H-i #2)$ | BDSC | BDSC: 27315<br>FlyBase: FBst0027315 |
| *D. melanogaster*; $y^l v^l$;<br>*P*[TRiP.HMS02286]*attP2*<br>(*p53-i*) | BDSC | BDSC: 41720<br>FlyBase: FBst0041720 |
| *D. melanogaster*; $w^*$,<br>*TI*[CRIMIC.TG4.1]*HCR60128-*<br>*TG4.1/TM3, Sb*<br>(*H-Gal4*) | BDSC | BDSC: 93469<br>FlyBase: FBst0093469 |
| *D. melanogaster*; $w^*$, *UASp-*<br>*myristoylated-GFP*<br>(*UAS-myrGFP*) | Hsu lab | |
| *D. melanogaster*; $w^*$, *UASp-*<br>*mCD8-GFP*<br>(*UAS-mCD8-GFP*) | Hsu lab | |
| *D. melanogaster*; $w^*$,<br>*Dpp::mCherry* | Hsu lab | |
| *D. melanogaster*; $w^*$,<br>*Hairless::GFP*<br>(*H::GFP*) on II | Sarah Bray (University<br>of Cambridge, England) | |
| *D. melanogaster*; $w^*$;<br>*Suppressor of Hairless-GFP*<br>(*Su(H)::GFP*) on III | Bray lab | |
| *D. melanogaster*; $w^*$; *Notch*<br>*responsive element-GFP* | Bray lab | |
| (NRE-GFP) |  |  |
| *D. melanogaster*; *w*<sup>*</sup>; *Notch*<br>*responsive element-RFP*<br>(NRE-RFP) | Bray lab |  |
| *D. melanogaster*; *y*<sup>l</sup> *w*<sup>*</sup>, *UAS-*<br>*lacZ/CyO*; *UAS-lacZ/ TM6C* | Bach lab |  |
| *D. melanogaster*; *w*<sup>*</sup> *upd-Gal4*,<br>*tub-Gal80<sup>ts</sup>*<br>(*upd-Gal4<sup>ts</sup>*) | Bach lab |  |
| *D. melanogaster*; *w*<sup>*</sup>; ;<br>*P[w[+mC]=UAS-*<br>*dpp.GFP.T]3/TM3, Sb*<br>(*UAS-Dpp::GFP*) | BDSC | BDSC: 53716<br>FlyBase: FBst0053716 |
| *D. melanogaster*; *y*<sup>l</sup> *w*<sup>*</sup>;<br>*P[w[+mC]=UASp-*<br>*gbb.GFP]3.3/ CyO*<br>(*UAS-Gbb::GFP*) | BDSC | BDSC: 63057<br>FlyBase: FBst0063057 |
| *D. melanogaster*; *y*<sup>l</sup> *w*<sup>*</sup>;<br>*P[w[+mC]=UASp-*<br>*gbb.GFP]3.3/ CyO*<br>(*UAS-Gbb-GFP*) | BDSC | BDSC: 63057<br>FlyBase: FBst0063057 |
| *D. melanogaster*; *y*<sup>l</sup> *w*<sup>*</sup> ;<br>*P[EPgy2]H<sup>EY03696</sup>*<br>(H<sup>EY</sup>) = UAS-H | BDSC | BDSC: 15672<br>FlyBase: FBst0015672 |
| *D. melanogaster*; *w*<sup>*</sup>;<br>*P[w[+mGS]=GSV1]H<sup>EP-211</sup>*<br>(H<sup>EP</sup>) = UAS-H | BDSC | BDSC: 43408<br>FlyBase: FBst0043408 |
| *D. melanogaster*; *y*, *w*<sup>67c23</sup>;<br>*P[GSV6,GS11776]*<br>(H<sup>GS</sup>) = UAS-H | Kyoto | Kyoto: 205338<br>FlyBase: FBti0104065 |
| Oligonucleotide |  |  |
| HCR probe of *chinmo-RA*, *-RD*,<br>*-RG*, *RH* | Molecular Instruments,<br>Inc. | LOT: PRJ996 |
| HCR probe of *chinmo-RE*, -*RF* | Molecular Instruments, Inc. | LOT: PRJ998 |
| HCR probe of *chinmo-RH* | Molecular Instruments, Inc. | LOT: PRJ997 |
| Software and Algorithms |  |  |
| ImageJ/Fiji | Fiji | http://fiji.sc/ |
| Photoshop/Illustrator | Adobe | https://www.adobe.com/products/ |
| Excel | Microsoft | https://products.office.com/en-us/excel |
| Prism 9 | GraphPad | https://www.graphpad.com/ |
| ZEN blue | Zeiss | https://www.zeiss.com/microscopy/en/products/software/zeiss-zen.html |

### Experimental model and subject details

*Drosophila melanogaster* strains used in this study are listed in the Key Resources Table. Flies were raised at 25°C on standard fly food except *nos* flip-out cassette crosses, which were maintained at 18°C until eclosion, and the adult flies were transferred to 29°C. We used only adult *Drosophila* males in this study and the vials were changed daily. The *Drosophila* stocks used in this study as follows: *nos-Gal4-VP16* ^63^; *UASp-3HA-Stat92E^FL#5^*^64^; *Stat92E^F^* ^65^; *Stat92E^85c9^* ^66^; *tj-Gal4 ^67^*; *UAS-LacZ* (BDSC #3956); *UAS-drosha-i* (BDSC#33657); *UAS-chinmo-i* (BDSC #33638); *UAS-Imp-i* (BDSC #38219); *UAS-Imp-i* (VDRC #20321); *UAS-Dcr1-i* (BDSC #34826); *UAS-control-*i (BDSC #36303); *UAS-syp-i* (VDRC #33011); *UAS-Ets21C-i* (BDSC #39069); *UAS-H-i* (BDSC #34703); *UAS-H-i* (BDSC#27315); *UAS-p53-i* (BDSC #41720); *H-Gal4* (BDSC#93469); *UAS-H^EY^* (BDSC#15672); *UAS-H^EP^* (BDSC#43408); *UAS-H^GS^* (Kyoto#205338); *Imp-GFP^#1^* (BDSC#60237) and *Imp-GFP^#2^* (Kyoto#110921); *UAS-Dpp::GFP* (BDSC#53716); *UAS-Gbb::GFP* (BDSC#63057); *UAS-Imp* (Tzumin Lee lab, University of Michigan, USA); *UAS-5’UTR-chinmo-3’UTR ^68^*; *UASp-NICD and Dpp::mCherry* (Hwei-Jan Hsu lab, Academia Sinica, Taiwan); *NRE-GFP*, *NRE-RFP, Hairless::GFP* and *Su(H)::GFP* (Sarah Bray lab, University of Cambridge, UK); *tub-Gal80^ts^*. For a list of full genotypes by figures, see Table S5.

### Detailed methods

#### RNAi-mediated gene depletion

To identify genes that genetically interact with Imp, we employed an RNA interference (RNAi) approach in *Drosophila*. Male UAS-RNAi transgenic flies (targeting *H*, *Ets21C*, or *p53*) were crossed with virgin females carrying the *nos*-Gal4 driver in combination with the temperature-sensitive *tub-Gal80^ts^*. All crosses were initially maintained at 18°C to minimize leaky expression of the RNAi constructs during development. 2- to 3-day-old adult males of the correct genotype were transferred to fresh vials and shifted to 29°C to induce robust RNAi expression. For the initial screening in Fig. S5A, cohorts of male flies were reared at 29 °C for 14 days prior to dissection, allowing assessment of Imp protein levels. A separate set of males were maintained at 29°C for 2, 28 or 42 days to investigate later-onset phenotypes or age-dependent effects.

### Generation of STAT-deficient GSCs

*Stat92E^F^* is a temperature-sensitive allele, and viable adults carrying *Stat92E^F^* can be obtained if this allele is *in trans* to a hypomorphic allele like *Stat92E^85c9^* and if the cross is reared at the permissive temperature 16°C until eclosion ^65^. When *Stat92E^F^*/*Stat92E^85c9^*adults are shifted to the restrictive temperature 29°C, Stat92E function is lost. *Stat92E*-mutant GSCs lose adhesion to the niche and differentiate. However, *Stat92E*-mutant GSCs can be maintained by somatic support cells (i.e., *tj-Gal4-*positive cells) expressing a full-length Stat92E transgene (i.e., *UASp-3HA-Stat92E^FL#5^*)^36^. We generated *y, w/Y; tj-Gal4/UASp-3HA-Stat92E^FL#5^; Stat92E^F^/Stat92E^85c9^*males or *w/Y; Stat92E^F^ e/+* controls, upshifted them to 29°C for 7 days, and then dissected testes and stained them for Chinmo.

### GSC clonal induction

Negatively-marked GSC clones were generated using the FLP/FRT technique after a single 1-hour heat shock at 37°C in 2-day-old adult males ^69^. Males were returned to 25°C until dissection at 2, 7, or 14 dpci. Positively marked clones were generated in 2-day old adult males by the MARCM technique after a single 1-hour heat shock in Figs. 7 and S1 and *nos* flip-out technique in Figs. 7 and S7, after 30-minute heat shock at 37°C ^54,55^. Males were returned to 25°C until dissection at 2 or 14 dpci. Lineage-wide mis-expression or depletion was achieved using the Gal4/UAS system ^70^. A *Gal80^ts^* transgene was for thermo-sensitive control of Gal4 ^71^.

### Immunofluorescence

Dissections and immunostaining were carried out as previously described ^33^. Briefly, testes were dissected in 1x phosphate buffered saline (PBS), fixed for 30 minutes in 4% paraformaldehyde (PFA) in 1xPBS, washed twice for 0.5 hours at 25°C in 1xPBS with 0.5% Triton X-100, and blocked in PBTB (1xPBS 0.2% Triton X-100 and 1% bovine serum albumin) for 1 hour at 25°C. Primary antibodies were incubated overnight at 4°C. Samples were washed twice for 30 minutes in PBTB and incubated for 2 hours in secondary antibody in PBTB at 25°C and then washed twice for 30 minutes in 1xPBS with 0.2% Triton X-100. Samples were mounted in Vectashield with DAPI (Vector Laboratories) or ibidi mounting Medium with DAPI (ibidi). Confocal images were captured using Zeiss LSM710, LSM900 and LSM980 microscopes with a 63x objective and 10–20 z-stacks (interval 1 μm) using lasers at 405, 488, 561, and 640 nM.

### RNA isolation and quantitative real-time reverse transcription PCR (qRT-PCR)

To measure gene expression in the anterior region of *Drosophila* testes (from GSCs to the spermatogonial stage), 150–200 testes were carefully dissected from adult males using fine steel pins under a stereomicroscope. The dissected tissue was immediately collected into lysis buffer, and total RNA was extracted using the Arcturus PicoPure™ RNA Isolation Kit (Thermo Fisher Scientific), following the manufacturer’s instructions. Next, 0.5 µg of total RNA was reverse-transcribed into complementary DNA (cDNA) using an M-MLV Reverse Transcription Kit (Protech Technology Enterprise Co., Ltd) with oligo(dT) primers. The resultant cDNA was diluted appropriately for downstream applications. Gene expression levels of *Imp* and *H* were normalized against the housekeeping genes *rpl19* and *rpl32*. qPCR reactions were set up using the KAPA SYBR® FAST qPCR Master Mix (Kapa Biosystems) on an ABI Prism™ real-time PCR system (Applied Biosystems). Each sample was run in triplicate to ensure technical reproducibility, and three biological replicates were analyzed per condition. All primer sequences used in this study are listed in Table S6.

Hybridization Chain Reaction - Fluorescent *in Situ* Hybridization (HCR-FISH) and RNA-FISH The HCR RNA probes against different *chinmo* mRNA isoforms, the HCR amplifier, and reagents for the hybridization, wash, amplification buffers were purchased from Molecular Instruments, Inc., HCR-FISH was performed according as previously described ^33^. RNA FISH was adapted from ^72^. Briefly, the immunostaining procedure testes were fixed in 4% PFA in 0.1% Triton X-100 in 1xPBS-DEPC for 30 minutes at 25°C, washed twice with 0.5% Triton X-100 in 1xPBS-DEPC for 30 minutes at 25°C. Samples were blocked in 0.1% Triton X-100 in 1xPBS-DEPC with 50 μg/mL heparin and 250 μg/mL yeast tRNA (PBTH), and then they were then incubated with primary antibodies overnight at 4°C. The next day, the samples were washed twice in PBTH for 30 minutes. Samples were then incubated with fluorescently labeled secondary antibodies in PBTH for 2 hours at 25°C. Samples were washed twice in PBTH for 30 minutes at 25°C. Samples were then dehydrated and rehydrated with a series of ethanol washes (25%, 50%, 75%, 100%) in 1xPBS-DEPC for 10 minutes at 25°C. Samples were treated for 3 minutes with 50 μg/mL Proteinase K, which was then inactivated by washing with 0.2% glycine twice in 1xPBS-DEPC for 5 minutes at 25°C. After Proteinase K treatment, the samples were fixed again in 4% PFA in 1xPBS-DEPC for 30 minutes at 25°C. The samples were pre-hybridized in freshly hybridization buffer+ (HyB^+^) containing 50% formamide, 5xSSC, 50 μg/mL heparin and 250 μg/mL yeast tRNA, 10 μg/mL Salmon sperm DNA with 0.1% Tween 20) for 10 minutes at 25°C and then incubated antisense *Imp* RNA probe in HyB^+^ overnight (12 - 16 hour) at 55°C. The sample was washed by Hyb+, rinsed in 2xSCC, and washed with PBT for 10 minutes. The samples were incubated with an anti-DIG antibody conjugated to horseradish peroxidase (HRP, Roche, 11207733910) at 4°C overnight. Finally, probes were detected using a TSA™ Cy3 kit (NEL704A001KT), following the manufacturer’s instructions. The antisense Dig-labeled *Imp* probe was made using open reading frame (ORF) PCR amplicons from complementary DNA libraries.

### Quantification and statistical analysis

Quantification of Chinmo, Imp, Imp::GFP and H::GFP in GSCs, Dpp::mCherry and Gbb in the GSC niche

Confocal Z-stacks were captured at 1 μm intervals to encompass all niche cells (typically 16–20 slices per testis). Fluorescence intensities of Chinmo, Imp, Imp::GFP, and H::GFP were measured using ImageJ in a single Z-slice taken at the maximal width of each GSC. In clonal experiments, the intensities of Chinmo and Imp in the mutant or knockdown GSCs were normalized to those of a non-mutant neighbor GSC within the same testis. Each data point represents one GSC or testis. For Dpp::mCherry and Gbb, signals were also measured in the z-slice containing the largest area of the niche. The background fluorescence was recorded from the nucleus of a muscle cell and subtracted from each measurement. Each data point here represents one testis.

### Quantification of Dg expression at the GSC-niche interface

We captured confocal Z-stacks (at 1 μm intervals) encompassing all niche cells (typically 16-20 slices). We measured Dg fluorescence intensity by ImageJ in single z slices at the area of maximal contact of a GSC with the niche. The background signal was measured in the nucleus of a gonialblast then subtracted from each GSC-niche measurement. In the mis-expression/ knockdown experiments, each data point represents the intensity of Dg in one GSC. In clonal analyses, the fluorescence intensity of Dg at the GSC-niche interface of the clone was normalized to that of a non-mutant neighbor GSC in the same testis. In the clonal analyses, each data point represents one testis.

### Quantification of *chinmo* and *Imp* mRNAs in the testis

We captured confocal Z-stacks (at 1 μm intervals) encompassing all niche cells and GSCs. We measured the number of fluorescence dots in the group of niche cells and in the single GSC. In niche analyses, each point represents niche cells. In GSC analyses, each point represents a single GSC.

### Quantification of Pcan intensity in the testicular lumen

We captured confocal Z-stacks (at 1 μm intervals) encompassing all niche cells (typically 16-20 slices). We measured the maximum area of Pcan fluorescence intensity by ImageJ in single z slices at the area near the niche. The background signal was measured in the nucleus of GSC and then subtracted from each other in the same testis. Each data point represents a single testis.

### Quantification of fold change in Pcan/Lan-positive Testes

Confocal Z-stacks (1 μm intervals) were captured to encompass all niche cells (typically 16–20 slices) in each testis, and the samples were stained with Pcan or Lan. A niche was counted as “Pcan/Lan-positive” if the ECM protein was detectable adjacent to at least one niche cell in any confocal slice. To quantify the fold change, we first calculated the percentage of Pcan/Lan-positive testes in each experimental condition and then divided that percentage by the corresponding percentage in the control. This ratio reflects the relative increase (or decrease) in Pcan/Lan-positive niches compared to control.

### Statistical analysis

Statistical analyses were performed using two-tailed Student’s t-tests, except for Figs. 3E, 3M, 5M, 6S, 7K, 7O, S1D, S7E and Table S1, which were performed using χ^2^ tests. Data were analyzed by GraphPad Prism and Microsoft Excel. Statistical significance was assumed by *P* ≤ 0.05. Individual *P* values are indicated. Data are represented by the mean and standard error of mean (SEM).

## Supplementary Tables S1-S6

**Table S1:** GSC clone induction rate, related to Figs. 7, S1, S7.

| # | Figure | % testes with at least 1 GSC clone | Time point (dpci) | Average | n | P (statistical test) |
| --- | --- | --- | --- | --- | --- | --- |
| 1 | S1D | control (*) | 2 | 71.43% | 28 |  |
| 2 | S1D | control (*) | 7 | 33.33% | 30 |  |
| 3 | S1D | control (*) | 14 | 16.67% | 24 |  |
| 4 | S1D | AGO1<sup>-/-</sup> (*) | 2 | 52.94% | 34 | 0.171 vs line 1 ($\chi^2$-test) |
| 5 | S1D | AGO1<sup>-/-</sup> (*) | 7 | 46.43% | 56 | 0.406 vs line 2 ($\chi^2$-test) |
| 6 | S1D | AGO1<sup>-/-</sup> (*) | 14 | 00.00% | 14 |  |
| 7 | 7 | control (^) | 2 | 36.00% | 18 |  |
| 8 | 7 | control (^) | 14 | 18.51% | 20 |  |
| 9 | 7 | *Imp-i* (^) | 2 | 35.68% | 26 | 0.826 vs line 7 ($\chi^2$-test) |
| 10 | 7 | *Imp-i* (^) | 14 | 31.81% | 36 | 0.259 vs line 8 ($\chi^2$-test) |
| 11 | S7 | *nos>>control-i* (&) | 2 | 60.00% | 18 |  |
| 12 | S7 | *nos>>control-i* (&) | 14 | 45.65% | 17 |  |
| 13 | S7 | *nos>>Imp-i* (&) | 2 | 58.06% | 36 | 0.877 vs line 11 ($\chi^2$-test) |
| 14 | S7 | *nos>>Imp-i* (&) | 14 | 60.86% | 29 | 0.097 vs line 12 ($\chi^2$-test) |
| 15 | 7 | *nos>>LacZ* (&) | 2 | 63.77% | 44 |  |
| 16 | 7 | *nos>>LacZ* (&) | 14 | 41.89% | 62 |  |
| 17 | 7 | *nos>>H<sup>EP</sup>* (&) | 2 | 46.51% | 20 | 0.073 vs line 15 ($\chi^2$-test) |
| 18 | 7 | *nos>>H<sup>EP</sup>* (&) | 14 | 62.42% | 93 | 0.171 vs line 16 ($\chi^2$-test) |
| 19 | 7 | *nos>>H<sup>GS</sup>* (&) | 2 | 57.78% | 26 | 0.521 vs line 15 ($\chi^2$-test) |
| 20 | 7 | *nos>>H<sup>GS</sup>* (&) | 14 | 39.06% | 25 | 0.406 vs line 16 ($\chi^2$-test) |

\* Denotes GFP-negative GSC clones generated by FLP/FRT
^ Denotes GFP-positive GSC clones generated by MARCM
& Denotes GFP-positive GSC clones generated by *nos* flip-out

**Table S2:** Clone occupancy, GSC number, number of neighbor GSCs, and number of GSC clones (clones induced by MARCM), related to Fig. 7.

|  | Figure | Label | Time point (dpci) | Average | SEM | n | P (statistical test) |
| --- | --- | --- | --- | --- | --- | --- | --- |
|  |  | **Clone occupancy** |  |  |  |  |  |
| **1** | 7P | control | 2 | 23.10% | ±3.35 | 18 |  |
| **2** | 7P | control | 14 | 21.12% | ±1.98 | 20 |  |
| **3** | 7P | *Imp-i* | 2 | 25.24% | ±3.63 | 26 |  |
| **4** | 7P | *Imp-i* | 14 | 44.38% | ±5.38 | 36 | 0.0199 vs line 2 (Student's t-test) |
|  |  | **Avg. number of GSCs** |  |  |  |  |  |
| **5** | 7Q | control | 2 | 6.56 | ±0.32 | 18 |  |
| **6** | 7Q | control | 14 | 6.50 | ±0.43 | 20 |  |
| **7** | 7Q | *Imp-i* | 2 | 6.54 | ±0.38 | 26 |  |
| **8** | 7Q | *Imp-i* | 14 | 5.06 | ±0.37 | 36 | 0.0233 vs line 6 (Student's t-test) |
|  |  | **Avg. number of neighbor GSCs** |  |  |  |  |  |
| **9** | 7R | control | 2 | 5.11 | ±0.43 | 18 |  |
| **10** | 7R | control | 14 | 5.13 | ±0.37 | 20 |  |
| **11** | 7R | *Imp-i* | 2 | 4.96 | ±0.42 | 26 |  |
| **12** | 7R | *Imp-i* | 14 | 2.91 | ±0.33 | 36 | 2.0 x 10<sup>-4</sup> vs line 10 (Student's t-test) |
|  |  | **Avg. number of GSC clones** |  |  |  |  |  |
| **13** | 7S | control | 2 | 1.44 | ±0.17 | 18 |  |
| **14** | 7S | control | 14 | 1.38 | ±0.17 | 20 |  |
| **15** | 7S | *Imp-i* | 2 | 1.58 | ±0.20 | 25 | 0.52 vs line 13 |
| **16** | 7S | *Imp-i* | 14 | 1.92 | ±0.30 | 36 | 0.1191 vs line 14 |

**Table S3:** Clone occupancy, GSC number, number of neighbor GSCs, and number of GSC clones (clones induced by *nos* flip-out), related to Fig. S7.

|  | Figure | Label | Time point (dpci) | Average | SEM | n | P (statistical test) |
| --- | --- | --- | --- | --- | --- | --- | --- |
|  |  | **Clone occupancy** |  |  |  |  |  |
| **1** | S7F | *nos>>control-i* | 2 | 22.21% | ±2.74 | 18 |  |
| **2** | S7F | *nos>>control-i* | 14 | 30.27% | ±1.63 | 17 |  |
| **3** | S7F | *nos>>Imp-i* | 2 | 36.82% | ±3.64 | 36 |  |
| **4** | S7F | *nos>>Imp-i* | 14 | 55.26% | ±4.47 | 29 | 0.003 vs line 18 (Student's t-test) |
|  |  | **Avg. number of GSC** |  |  |  |  |  |
| **5** | S7G | *nos>>control-i* | 2 | 6.67 | ±0.27 | 18 |  |
| **6** | S7G | *nos>>control-i* | 14 | 7.11 | ±0.25 | 17 |  |
| **7** | S7G | *nos>>Imp-i* | 2 | 5.65% | ±0.19 | 36 |  |
| **8** | S7G | *nos>>Imp-i* | 14 | 4.44% | ±0.25 | 29 | 3X10<sup>4</sup> vs line 22 (Student's t-test) |
|  |  | **Avg. number of neighbor GSCs** |  |  |  |  |  |
| **9** | S7H | *nos>>control-i* | 2 | 5.17 | ±0.27 | 18 |  |
| **10** | S7H | *nos>>control-i* | 14 | 3.49 | ±0.26 | 17 |  |
| **11** | S7H | *Imp-i* | 2 | 4.94 | ±0.26 | 36 |  |
| **12** | S7H | *Imp-i* | 14 | 2.07 | ±0.23 | 29 | 2.0 x 10<sup>-4</sup> vs line 10 (Student's t-test) |
|  |  | **Avg. number of GSC clone** |  |  |  |  |  |
| **13** | S7I | *nos>>control-i* | 2 | 1.28 | ±0.13 | 18 |  |
| **14** | S7I | *nos>>control-i* | 14 | 2.16 | ±0.22 | 17 |  |
| **15** | S7I | *nos>>Imp-i* | 2 | 1.72 | ±0.21 | 36 | 0.0851 vs line 13 |
| **16** | S7I | *nos>>Imp-i* | 14 | 2.37 | ±0.23 | 29 | 0.5379 vs line 14 |

**Table S4:**
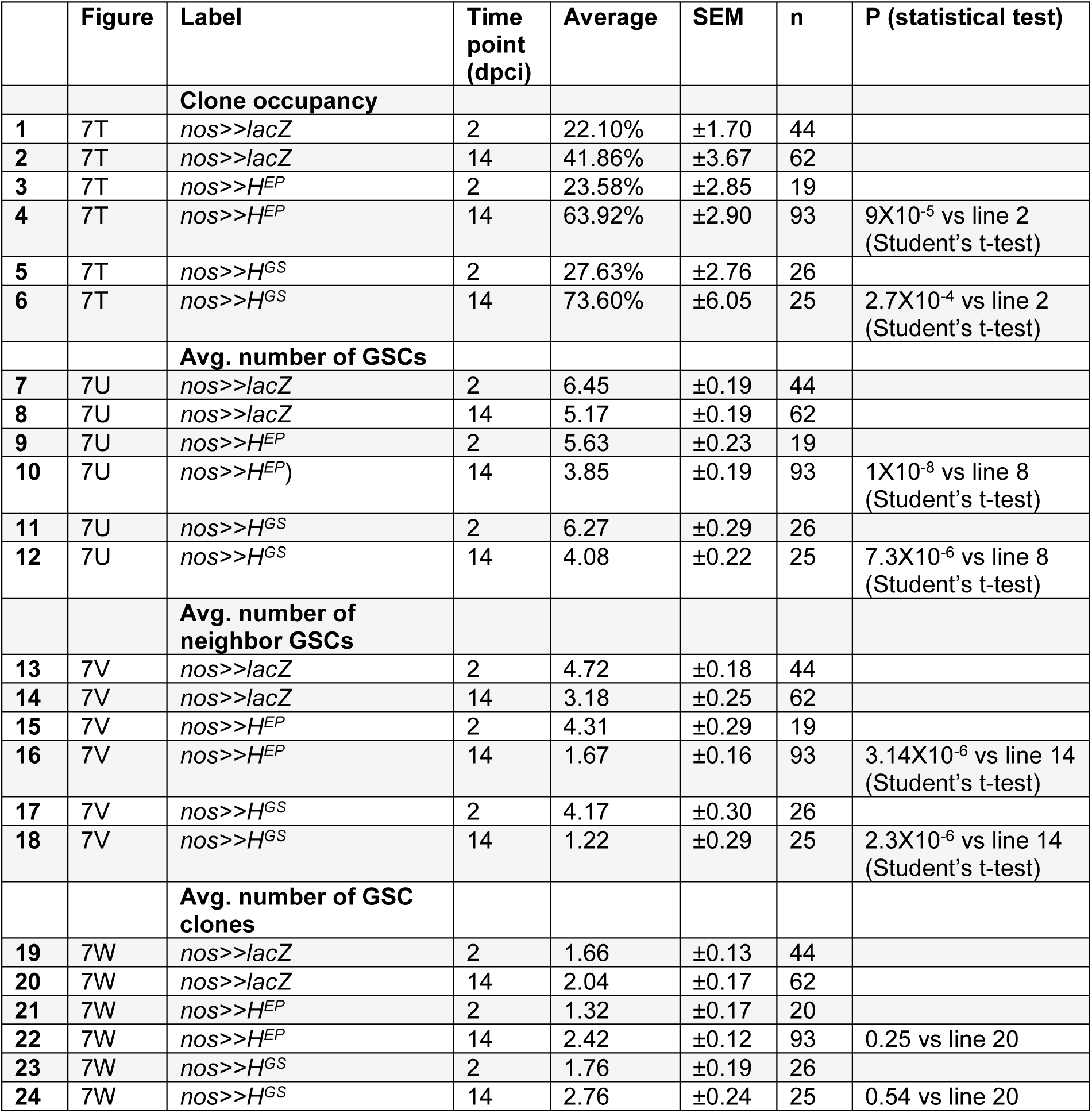
Clone occupancy, GSC number, number of neighbor GSCs, and number of GSC clones (clones induced by *nos* flip-out), related to Fig. 7.

|  | Figure | Label | Time point (dpci) | Average | SEM | n | P (statistical test) |
| --- | --- | --- | --- | --- | --- | --- | --- |
|  |  | **Clone occupancy** |  |  |  |  |  |
| 1 | 7T | *nos>>lacZ* | 2 | 22.10% | ±1.70 | 44 |  |
| 2 | 7T | *nos>>lacZ* | 14 | 41.86% | ±3.67 | 62 |  |
| 3 | 7T | *nos>>H<sup>EP</sup>* | 2 | 23.58% | ±2.85 | 19 |  |
| 4 | 7T | *nos>>H<sup>EP</sup>* | 14 | 63.92% | ±2.90 | 93 | 9X10<sup>-5</sup> vs line 2 (Student's t-test) |
| 5 | 7T | *nos>>H<sup>GS</sup>* | 2 | 27.63% | ±2.76 | 26 |  |
| 6 | 7T | *nos>>H<sup>GS</sup>* | 14 | 73.60% | ±6.05 | 25 | 2.7X10<sup>-4</sup> vs line 2 (Student's t-test) |
|  |  | **Avg. number of GSCs** |  |  |  |  |  |
| 7 | 7U | *nos>>lacZ* | 2 | 6.45 | ±0.19 | 44 |  |
| 8 | 7U | *nos>>lacZ* | 14 | 5.17 | ±0.19 | 62 |  |
| 9 | 7U | *nos>>H<sup>EP</sup>* | 2 | 5.63 | ±0.23 | 19 |  |
| 10 | 7U | *nos>>H<sup>EP</sup>* | 14 | 3.85 | ±0.19 | 93 | 1X10<sup>-8</sup> vs line 8 (Student's t-test) |
| 11 | 7U | *nos>>H<sup>GS</sup>* | 2 | 6.27 | ±0.29 | 26 |  |
| 12 | 7U | *nos>>H<sup>GS</sup>* | 14 | 4.08 | ±0.22 | 25 | 7.3X10<sup>-6</sup> vs line 8 (Student's t-test) |
|  |  | **Avg. number of neighbor GSCs** |  |  |  |  |  |
| 13 | 7V | *nos>>lacZ* | 2 | 4.72 | ±0.18 | 44 |  |
| 14 | 7V | *nos>>lacZ* | 14 | 3.18 | ±0.25 | 62 |  |
| 15 | 7V | *nos>>H<sup>EP</sup>* | 2 | 4.31 | ±0.29 | 19 |  |
| 16 | 7V | *nos>>H<sup>EP</sup>* | 14 | 1.67 | ±0.16 | 93 | 3.14X10<sup>-6</sup> vs line 14 (Student's t-test) |
| 17 | 7V | *nos>>H<sup>GS</sup>* | 2 | 4.17 | ±0.30 | 26 |  |
| 18 | 7V | *nos>>H<sup>GS</sup>* | 14 | 1.22 | ±0.29 | 25 | 2.3X10<sup>-6</sup> vs line 14 (Student's t-test) |
|  |  | **Avg. number of GSC clones** |  |  |  |  |  |
| 19 | 7W | *nos>>lacZ* | 2 | 1.66 | ±0.13 | 44 |  |
| 20 | 7W | *nos>>lacZ* | 14 | 2.04 | ±0.17 | 62 |  |
| 21 | 7W | *nos>>H<sup>EP</sup>* | 2 | 1.32 | ±0.17 | 20 |  |
| 22 | 7W | *nos>>H<sup>EP</sup>* | 14 | 2.42 | ±0.12 | 93 | 0.25 vs line 20 |
| 23 | 7W | *nos>>H<sup>GS</sup>* | 2 | 1.76 | ±0.19 | 26 |  |
| 24 | 7W | *nos>>H<sup>GS</sup>* | 14 | 2.76 | ±0.24 | 25 | 0.54 vs line 20 |

**Table 5:** Genotypes in each figure.

| Figure | Full genotype | Abbreviation used in figure |
| --- | --- | --- |
| 1B-E, G-O | *w*<sup>1118</sup> | WT |
| 1E | *y, w/Y; nos-Gal4/+; UAS-lacZ/+* | *nos>lacZ* |
| 1E | *y, w/Y; nos-Gal4/+; UAS-chimmo*<sup>HMS00036</sup><br>*RNA-i/+* | *nos>chinmo-i* |
| 2A-F, I | *Oregon-R* | WT |
| 2G-I | *y, w* | WT |
| 2J-L | *P{PTT-un}Imp*<sup>ZCL2884X</sup>/*Y* | *Imp::GFP #1* |
| 2M, O, P | *w/Y; nos-Gal4/+; UAS-lacZ/+* | *nos>lacZ* |
| 2N, O, P | *w/Y; nos-Gal4/+; UAS-Imp-HA/+* | *nos>Imp* |
| 3A, C, E, F, H | *w/Y; nos-Gal4/+; UAS-lacZ/+* | *nos>lacZ* |
| 3B, D, E, G, H | *w/Y; nos-Gal4/+; UAS-Imp-HA/+* | *nos>Imp* |
| 3I, M, N, R, S | *w/Y; nos-Gal4/UAS-lacZ; UAS-lacZ/+* | *nos>lacZ, lacZ* |
| 3J, M, O, R, S | *w/Y; nos-Gal4/UAS-lacZ/+; UAS-chimmo*<sup>HMS00036</sup><br>*RNA-i/+* | *nos>lacZ, chinmo-i* |
| 3K, M, P, R, S | *w/Y; nos-Gal4/UAS-Imp-i*<sup>GL00660</sup>; *UAS-lacZ/+* | *nos>Imp-i, lacZ* |
| 3L, M, Q-S | *w/Y; nos-Gal4/UAS-Imp-i*<sup>GL00660</sup>; *UAS-chinmo/+* | *nos>Imp-i, chinmo* |
| 4A, B | *y, w/Y; H-Gal4/+; UASp-myr::GFP/+* | *H>myrGFP* |
| 4C | *y, w/Y; nos-Gal4/+; UASp-myr::GFP/+* | *nos>myrGFP* |
| 4D | *Oregon-R* | WT |
| 4E-H | *w/Y; Hairless::GFP* | *H::GFP* |
| 4I, K-N | *y, w/Y; nos-Gal4/+; UAS-control RNAi/+* | *nos>control-i* |
| 4J-N | *y, w/Y; nos-Gal4/+; UAS-H*<sup>HMS01182</sup><br>*RNAi/+* | *nos>H-i* |
| 5A-C | *y, w/Y; Dpp::mCherry/+* | *Dpp::mCh* |
| 5D-F | *y, w/Y* | WT |
| 5G, J | *upd-Gal4, tubGal80<sup>ts</sup>/Y* | *upd>control* |
| 5K-N | *upd-Gal4, tubGal80<sup>ts</sup>/Y; UAS-mCD8::GFP/+* | *upd>mCD8::GFP* |
| 5K-N | *upd-Gal4, tubGal80<sup>ts</sup>/Y; UAS-dpp::GFP/+* | *upd>Dpp::GFP* |
| 5K-N | *upd-Gal4, tubGal80<sup>ts</sup>/Y; UAS-Gbb::GFP/+* | *upd>Gbb::GFP* |
| 6A, C, D, F, G, I, J | *y, w/Y; nos-Gal4/+; UAS-control RNAi/+* | *nos>control-i* |
| 6B, C, E, F, H, I, J | *y, w/Y; nos-Gal4/+; UAS-H<sup>HMS01182</sup>RNAi/+* | *nos>H-i* |
| 6K, M, N, P, Q, S-W | *y, w/ Y; nos-Gal4/+; UAS-lacZ/ +* | *nos>lacZ* |
| 6L, M, O, P, R, S, U-W | *y, w/ Y; nos-Gal4/+; UAS-H<sup>EY03696</sup>/+* | *nos>H<sup>EY</sup>* |
| 6P, S, V, W | *y, w/ Y; nos-Gal4/+; UAS-H<sup>EP-211</sup>/ +* | *nos>H<sup>EP</sup>* |
| 6P, S, V, W | *y, w/ Y; nos-Gal4/+; UAS-H<sup>GS11776</sup>/ +* | *nos>H<sup>GS</sup>* |
| 7C, L-O, T-W | *y, w, hs-flp/Y; nos-FRT-GAL4-VP16-nos 3'UTR/UASp-gfp; UAS-lacZ/+* | *nos>>lacZ* |
| 7D, L-O, T-W | *y, w, hs-flp/Y; nos-FRT-GAL4-VP16-nos-3'UTR/UASp-gfp; UAS-H<sup>EP-211</sup>/ +* | *nos>>H<sup>EP</sup>* |
| 7E, F, H, J, K, P-S | *y, w, hs-flp<sup>122</sup>, tub-Gal4, UAS-nls-GFP/Y; tub-Gal80, FRT<sup>40A</sup>/FRT<sup>40A</sup>; UAS-Imp<sup>GD9232</sup>RNAi/+* | *Imp-i clone* |
| 7F, G, I, K, P-S | *y, w, hs-flp<sup>122</sup>, tub-Gal4, UAS-nls-GFP/Y; tub-Gal80, FRT<sup>40A</sup>/FRT<sup>40A</sup>* | *control clone* |
| 7N, P-W | *y, w, hs-flp/Y; nos-mCherry-SV40-polyA-FRT-GAL4-VP16-nos 3'UTR/UASp-gfp; UAS-lacZ/+* | *nos>>lacZ* |
| 7L-O, T-W | *y, w, hs-flp/Y; nos-FRT-GAL4-VP16-nos 3'UTR/UASp-gfp; UAS-H<sup>GS11776</sup>/+* | *nos>>H<sup>GS</sup>* |
| S1A, C | *y, w, hs-flp<sup>122</sup>, tub-Gal4, UAS-nls-GFP/Y; FRT<sup>82B</sup>, tub-Gal80/ FRT<sup>82B</sup>, Dcr-1<sup>Q1147XA</sup>* | *Dcr-1<sup>-/-</sup> clone* |
| S1B-D | *y, w, hs-flp<sup>122</sup>/Y; FRT<sup>42D</sup>, ubi-GFP/ FRT<sup>42D</sup>, AGO<sup>1k08121</sup>* | *AGO1<sup>-/-</sup> clone* |
| S1D | *y, w, hs-flp<sup>122</sup>/Y; FRT<sup>42D</sup>, ubi-GFP/FRT<sup>42D</sup>* | *control clone* |
| S1E, F, I | *y, w/Y; nos-Gal4/+; UAS-control-RNAi/+* | *nos>control-i* |
| S1G, I | *y, w/Y; nos-Gal4/+; UAS-drosha<sup>HMS00064</sup> RNAi/+* | *nos>drosha-i* |
| S1H, I | *y, w/Y; nos-Gal4/UAS-mCherry.mir-let7.sponge; UAS-mCherry.mir-let7.sponge/+* | *nos>mir-let7.sponge* |
| S1J, L | *+/Y; Stat92E<sup>F</sup>, e/+* | *control* |
| S1K, L | *y, w/Y; tj-Gal4/UASp-3HA-Stat92E<sup>FL#5</sup>; Stat92E<sup>F</sup>, e/Stat92E<sup>85c9</sup>, ca, e* | *tj>HA-Stat92E; Stat92E<sup>F/85c9</sup>* |
| S2A-C | *w\*/Y; P{PTT-un}Imp<sup>ZCL2884X</sup>* | *Imp::GFP<sup>#2</sup>* |
| S2D, E, G | *y, w/Y; nos-Gal4/+; UAS-lacZ/+* | *nos>lacZ* |
| S2F, G | *y, w/Y; nos-Gal4/+; UAS-Imp-HA/+* | *nos>Imp* |
| S3A, D, E, F, H, I | *y, w/Y; nos-Gal4/+; UAS-control-RNAi/+* | *nos>control-i* |
| S3B, D, H, I | *y, w/Y; nos-Gal4/+; UAS-Imp-i<sup>GL00660</sup>/+* | *nos>Imp-i<sup>B#38219</sup>* |
| S3C, D, G, H, I | *y, w/Y; nos-Gal4/+; UAS-Imp-i<sup>GD9232</sup>/+* | *nos>Imp-i<sup>V#20321</sup>* |
| S4A, D, G, I, J | *y, w/Y; nos-Gal4/+; UAS-control-RNAi/+* | *nos>control-i* |
| S4B, D | *y, w/Y; nos-Gal4/+; UAS-dicer1<sup>HMS00141</sup> RNAi/+* | *nos>Dcr1-i* |
| S4C, D | *y, w/Y; nos-Gal4/+; UAS-drosha<sup>HMS00064</sup> RNAi/+* | *nos>drosha-i* |
| S4E, F | *Oregon-R* | *WT* |
| S4H, I, J | *y, w/Y; nos-Gal4/UAS-syp<sup>GD9477</sup> RNAi* | *nos>Syp-i* |
| S4K, M | *y, w/ Y; nos-Gal4/+; UAS-lacZ/+* | *nos>lacZ* |
| S4L, M | *y, w/ Y; nos-Gal4 /+; UAS-5'UTR-Chinmo-3'UTR/+* | *nos>chinmo* |
| S5A, E, G, H, J, K, M, N, P, Q | *y, w/ Y; nos-Gal4/+; UAS-control-RNAi/+* | *nos>control-i* |
| S5A | *y, w/Y; nos-Gal4/+; UAS-H<sup>HMS01182</sup> RNAi/+* | *nos>H-i* |
| S5A | *y, w/Y; nos-Gal4/+; UAS-Ets21<sup>HMS01989</sup> RNAi/+* | *nos>Ets21-i* |
| S5A | *y, w/Y; nos-Gal4/+; UAS-p53<sup>HMS02286</sup> RNAi/+* | *nos>p53-i* |
| S5B, D | *Imp*<sup>MI05901-GFSTF.2</sup>/Y; nos-Gal4/+; UAS-control-RNAi/+ | *nos>control-i* |
| S5C, D | *Imp*<sup>MI05901-GFSTF.2</sup>/Y; nos-Gal4/+; UAS-*H*<sup>HMS01182</sup> RNAi/+ | *nos>H-i* |
| S5F, G, I, J, L, M, O, P, Q | *y, w/Y; nos-Gal4/+; UAS-H*<sup>JF02624</sup> RNAi/+ | *nos>H-i#2* |
| S6A-C | *w/Y; Suppressor of Hairless (Su(H))::GFP* | *Su(H)::GFP* |
| S6D, F | *w\*/Y; Notch responsive element (NRE)-GFP* | *NRE-GFP* |
| S6E, G | *w\*/Y; Notch responsive element (NRE)-RFP* | *NRE-RFP* |
| S6H | *y, w/Y; nos-Gal4/NRE-GFP; UAS-control-RNAi/+* | *nos>control-i* |
| S6I | *y, w/Y; nos-Gal4/ NRE-GFP; UAS-H*<sup>HMS01182</sup> RNAi/+ | *nos>H-i* |
| S6J | *y, w/Y; nos-Gal4/NRE-RFP; UAS-control-RNAi/+* | *nos>control-i* |
| S6K | *y, w/Y; nos-Gal4/ NRE-RFP; UAS-H*<sup>HMS01182</sup> RNAi/+ | *nos>H-i* |
| S6L, N, O | *y, w/Y; nos-Gal4/ +; UAS-lacZ/+* | *nos>lacZ* |
| S6M-O | *y, w/Y; nos-Gal4/+; UAS-N*<sup>ICD</sup>/ + | *nos>N*<sup>ICD</sup> |
| S7A, C, E-I | *y, w, hs-flp/Y; nos-FRT-gal4-VP16-nos 3'UTR/UASp-GFP; UAS-control-RNAi/+* | *nos>>control-i* |
| S7B, D, E-I | *y, w, hs-flp/Y; nos-FRT-gal4-VP16-nos 3'UTR/UASp-GFP; UAS-Imp-i*<sup>B#38219</sup>/+ | *nos>>Imp-i*<sup>B#38219</sup> |
| Table S1 | *y, w, hs-flp*<sup>122</sup>/Y; *FRT*<sup>42D</sup> *ubi-GFP/FRT*<sup>42D</sup> | control (*) |
|  | *y, w, hs-flp*<sup>122</sup>/Y; *FRT*<sup>42D</sup> *ubi-GFP/FRT*<sup>42D</sup> *AGO1*<sup>k08121</sup> | *AGO1*<sup>-/-</sup> (*) |
|  | *y, w, hs-flp*<sup>122</sup>, *tub-Gal4, UAS-nls-GFP/Y; tub-Gal80, FRT*<sup>40A</sup>/*FRT*<sup>40A</sup> | control (^) |
|  | *y, w, hs-flp*<sup>122</sup>, *tub-Gal4, UAS-nls-GFP/Y; tub-Gal80, FRT*<sup>40A</sup>/*FRT*<sup>40A</sup>; UAS-Imp<sup>GL00660</sup> RNAi/+ | *Imp-i* (^) |
|  | *y, w, hs-flp/Y; nos-FRT-gal4-VP16-nos 3'UTR/UASp-GFP; UAS-control-RNAi/+* | *nos>>control-i (&)* |
|  | *y, w, hs-flp/Y; nos-FRT-gal4-VP16-nos 3'UTR/UASp-GFP; UAS-Imp-i<sup>GL00660</sup>/+* | *nos>>Imp-i (&)* |
|  | *y, w, hs-flp/Y; nos-FRT-gal4-VP16-nos 3'UTR/UASp-GFP; UAS-lacZ/+* | *nos>>LacZ (&)* |
|  | *y, w, hs-flp/Y; nos-FRT-gal4-VP16-nos 3'UTR/UASp-GFP; UAS-H<sup>EP-211</sup>/ +* | *nos>>H<sup>EP</sup> (&)* |
|  | *y, w, hs-flp/Y; nos-FRT-gal4-VP16-nos 3'UTR/UASp-GFP; UAS-H<sup>GS11776</sup>/+* | *nos>>H<sup>GS</sup> (&)* |
| Table S2 | *y, w, hs-flp<sup>122</sup>, tub-Gal4, UAS-nls-GFP/Y; tub-Gal80, FRT<sup>40A</sup>/FRT<sup>40A</sup>* | control |
|  | *y, w, hs-flp<sup>122</sup>, tub-Gal4, UAS-nls-GFP/ Y; tub-Gal80, FRT<sup>40A</sup>/FRT<sup>40A</sup>; UAS-Imp<sup>GL00660</sup> RNAi/+* | *Imp-i* |
| Table S3 | *y, w, hs-flp/Y; nos-FRT-gal4-VP16-nos 3'UTR/UASp-GFP; UAS-control-RNAi/+* | *nos>>control-i* |
|  | *y, w, hs-flp/Y; nos-FRT-gal4-VP16-nos 3'UTR/UASp-GFP; UAS-Imp-i<sup>GL00660</sup>/+* | *nos>>Imp-i* |
| Table S4 | *y, w, hs-flp/Y; nos-FRT-gal4-VP16-nos 3'UTR/UASp-GFP; UAS-lacZ/ +* | *nos>>LacZ* |
|  | *y, w, hs-flp/Y; nos-FRT-gal4-VP16-nos 3'UTR/UASp-GFP; UAS-H<sup>EP-211</sup>/ +* | *nos>>H<sup>EP</sup>* |
|  | *y, w, hs-flp/Y; nos-FRT-gal4-VP16-nos 3'UTR/UASp-GFP; UAS-H<sup>GS11776</sup>/+* | *nos>>H<sup>GS</sup>* |

**Table S6.**
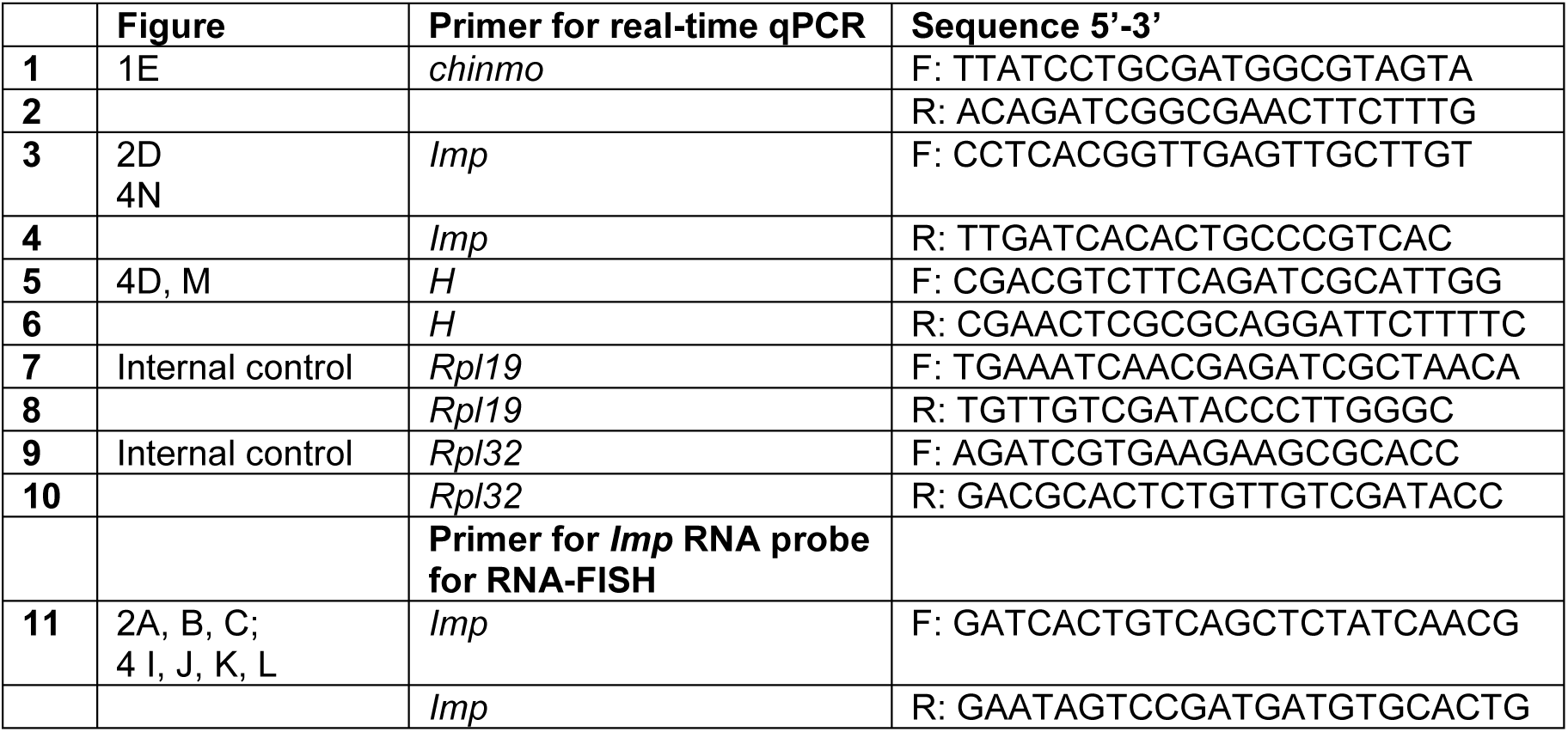
Primers for real-time qRT-PCR and RNA-FISH, related to Figures 1, 2, 5.

## Supplementary figures and legends S1-S7

**Figure S1.**
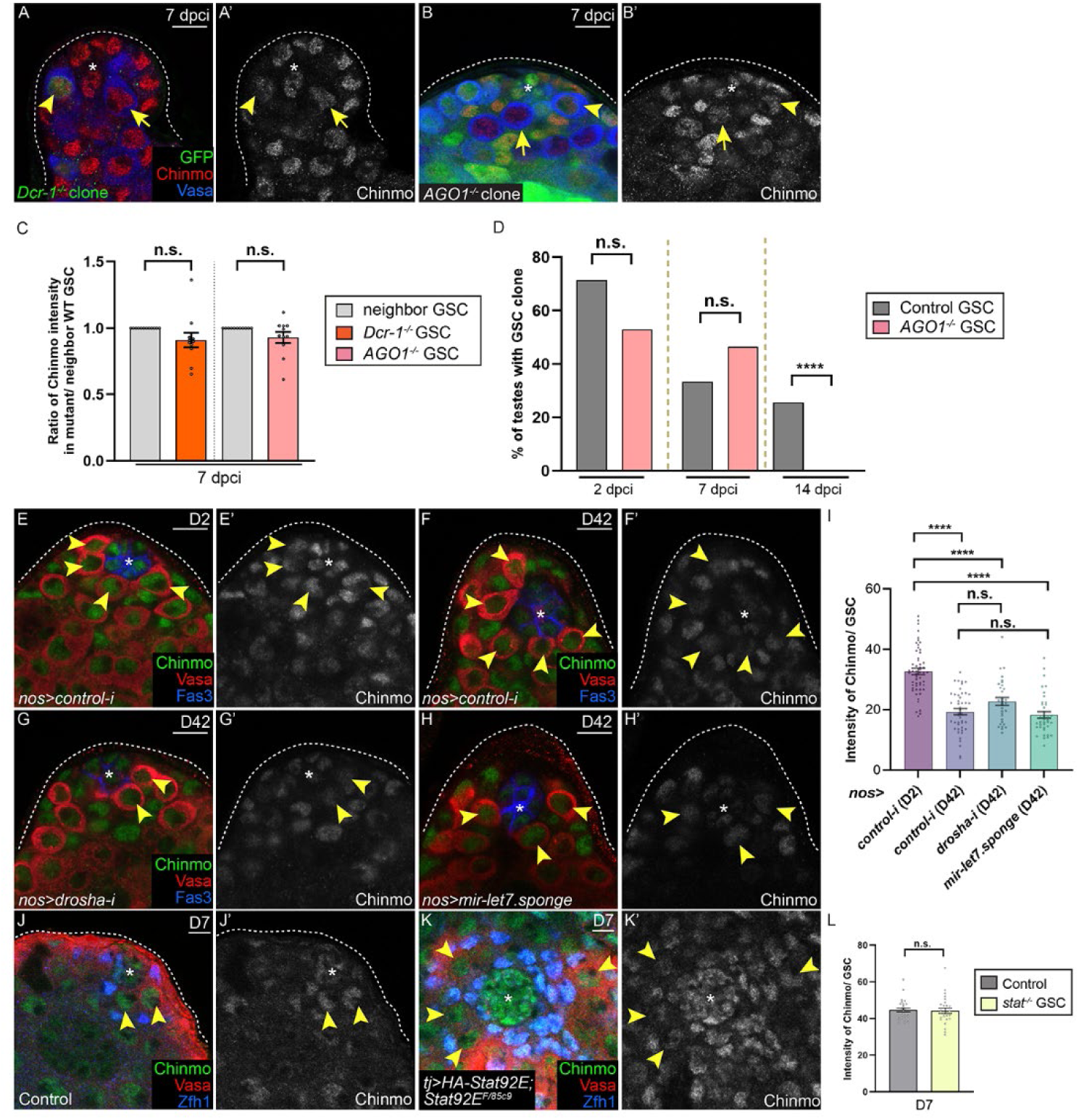
Chinmo is not regulated by the miRNA machinery in GSCs, related to Fig. 1. (A, B) Confocal images of Chinmo (red) in testes with (A) MARCM *Dcr-1^-/-^* GSC clones (A, arrowhead) or (B) GFP-negative *AGO1^-/^*^-^ GSC clones (B, arrowhead) at 14 days post-clone induction (dpci). Arrows indicate non-mutant neighbor GSCs. (C) Graph showing the ratio of Chinmo protein in *Dcr-1^-/-^* (orange bar) or *AGO1^-/^*^-^ GSC clones (pink bar) compared to non-mutant neighbor GSCs (gray bars) per testis. (D) Graph quantifying the percentage of testes harboring control (gray bars) or *AGO1^-/^*^-^ (pink bars) GSC clones at 2, 7, and 14 dpci. (E–I) Confocal images of Chinmo (green) in D2 (E) or D42 (F–H) *nos>control-i* (E, F), *nos>drosha-i* (G), or *nos>mir-let7.sponge* (H) testes, quantified in I. (J-L) Confocal images of Chinmo protein (green) in control (*Stat92E^F^/+*, J) or in STAT-deficient GSCs maintained by STAT-expressing, Tj-positive somatic support cells (*tj-Gal4/UASp-3HA-Stat92E^FL#5^; Stat92E^F/85c9^*, K), quantified in L. Note in K the Vasa-positive GSC-like cells (arrowheads) are located 1 cell diameter’s distance away from the niche. Arrowheads label GSCs in E-H, J and GSC-like cells in K. Asterisks indicate the niche. Vasa marks germ cells and Fas3 marks niche cells. Zfh-1 labels CySCs and their immediate daughter cells. Scale bars = 10 μm. Error bars in C, I, L represent SEM. **** P ≤ 0.0001; n.s. = not significant, as assessed by Student’s t-test and by χ^2^ test in D (see STAR Methods)

**Figure S2.**
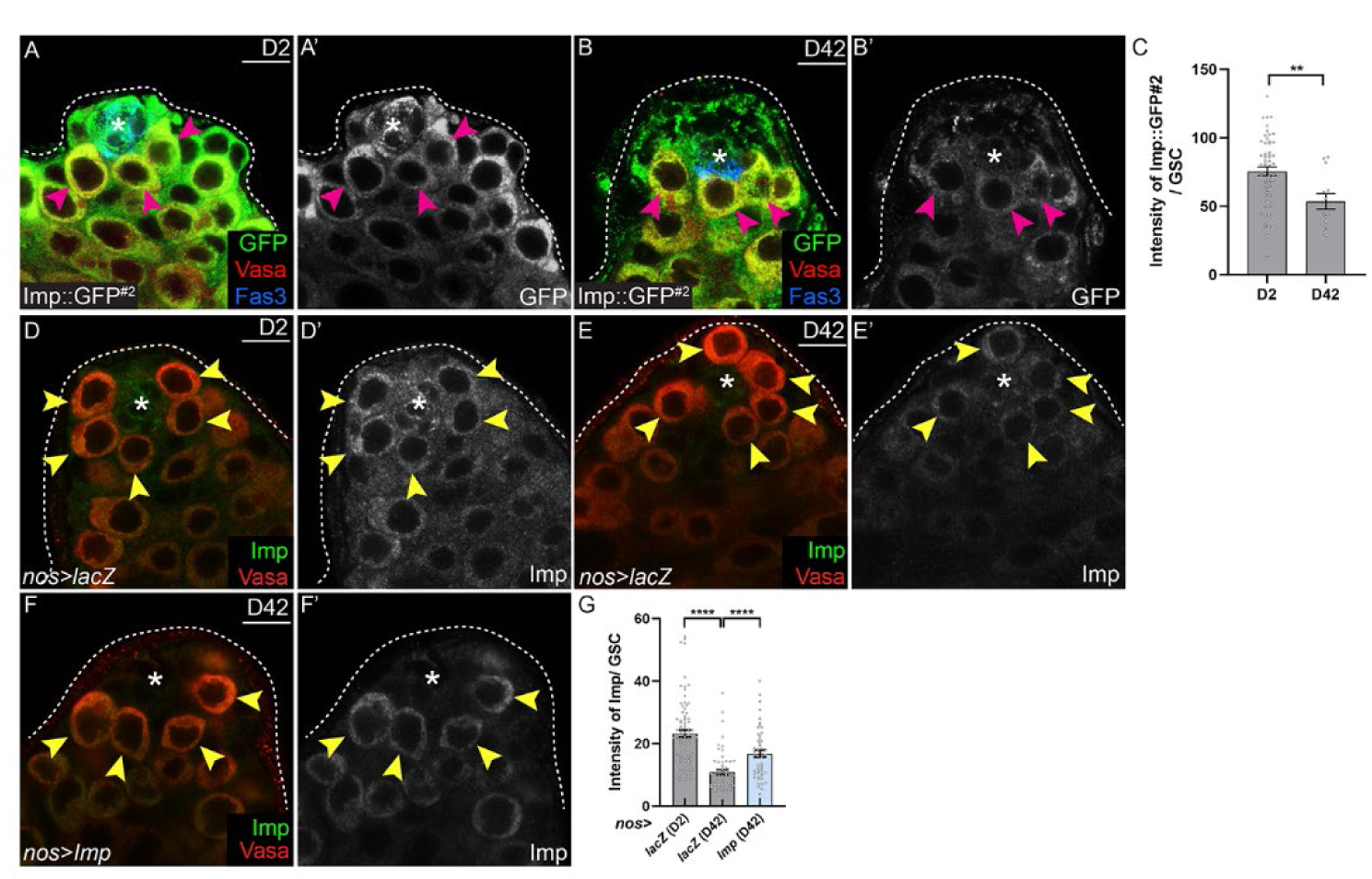
Imp is decreased in GSCs during aging, related to Fig. 2. (A–C) Confocal images of Imp::GFP #2 protein trap (green) in D2 (A) or D28 testes (B), quantified in C. Arrowheads indicate Imp::GFP #2 expression in GSCs. (D–G) Confocal images of Imp protein (green) in D2 and D42 *nos>lacZ* (D, E) or D42 *nos>Imp* (F) testes, quantified in G. Arrowheads mark Imp expression in GSCs. Asterisks indicate niche cells. Vasa marks germ cells and Fas3 labels niche cells. Scale bars = 10 μm. Error bars in C, G represent SEM. **P ≤ 0.01: **** P ≤ 0.0001 by Student’s t-test (see STAR Methods).

**Figure S3.**
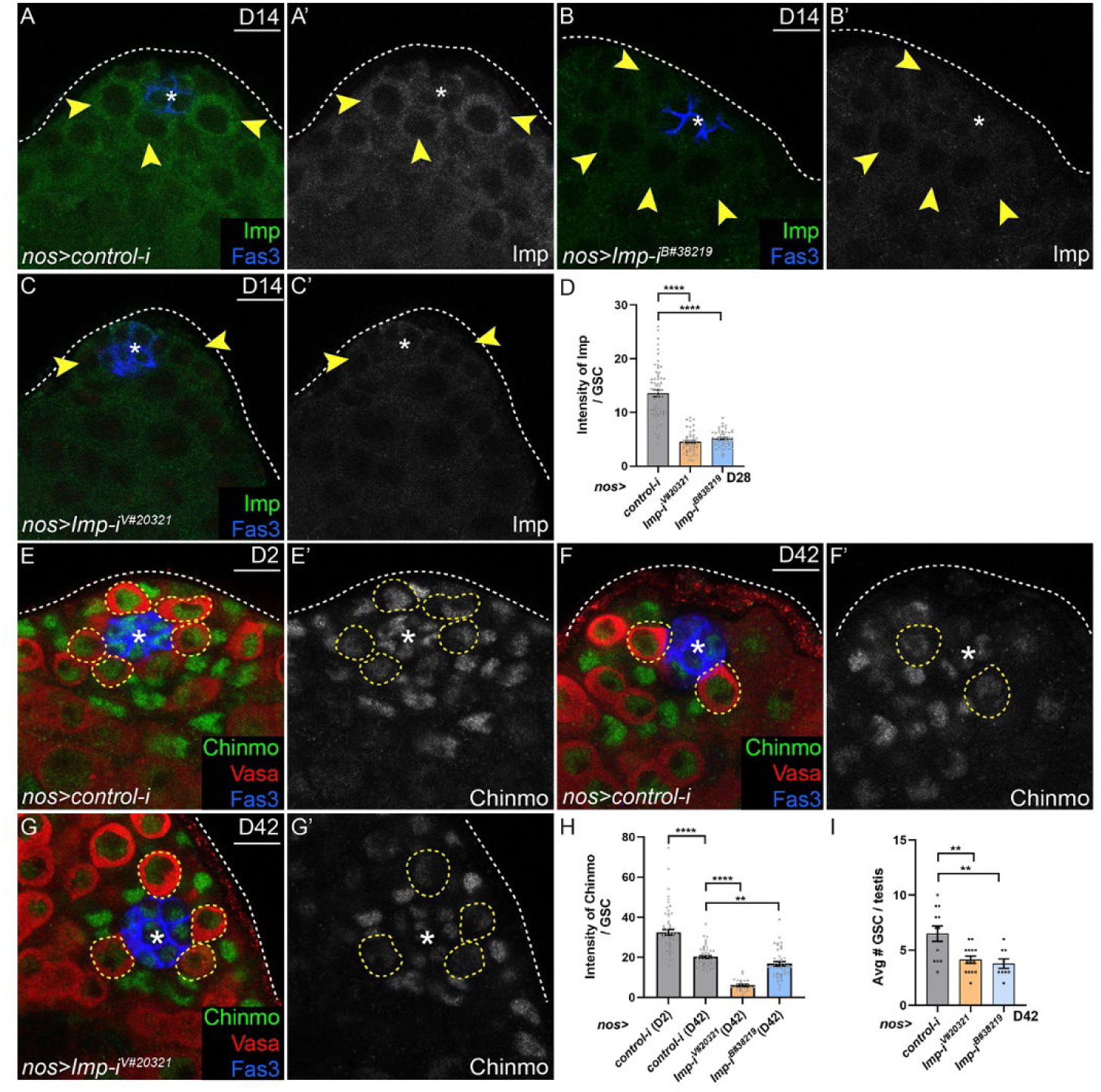
Imp RNAi efficiently depletes Imp, leading to reduced Chinmo and impaired GSC homeostasis, related to Fig. 3. (A–D) Confocal images of Imp protein (green) in D14 *nos>control-i* (A), *nos>Imp-i^B#38219^* (B), or *nos>Imp-i^V#20321^* (C) testes, quantified in D. Arrowheads indicate Imp expression in GSCs. (E–H) Confocal images of Chinmo (green) in D2 or D42 *nos>control-i* testes (E, F), and D42 *nos>Imp-i^V#20321^* (G) testes, quantified in H. Dashed lines indicate GSCs. (I) Graph showing the number of GSCs in D2 or D42 *nos>control-i* (gray bar), D42 *nos>Imp-i^B#38219^*(orange bar), or D42 *nos>Imp-i^V#20321^* (blue bar) testes. Asterisks indicate niche cells. Vasa marks germ cells and Fas3 labels niche cells. Scale bars = 10 μm. Error bars in H, Irepresent SEM. ** P ≤ 0.01; ****; P ≤ 0.0001 by Student’s t-test (see STAR Methods).

**Figure S4.**
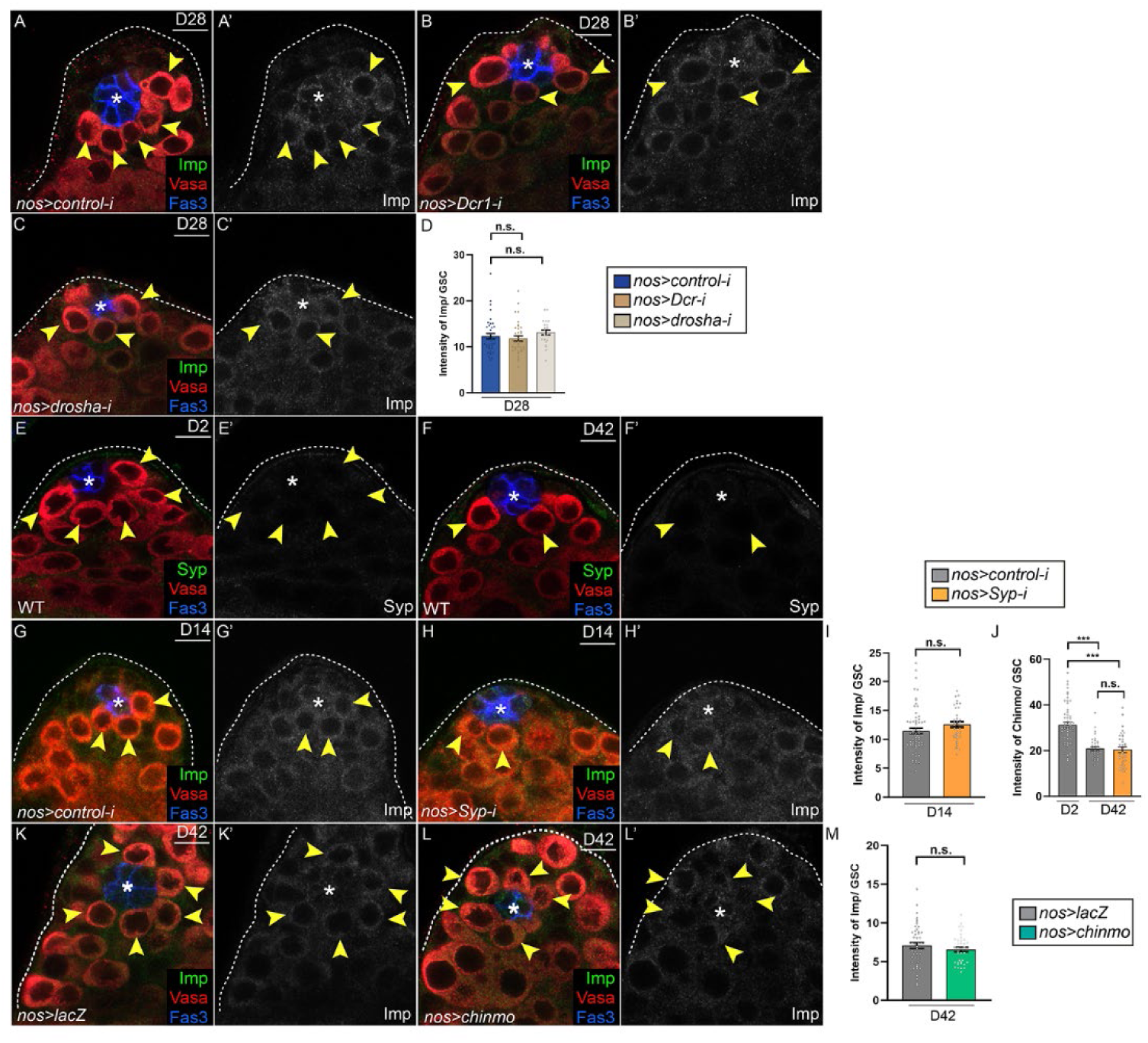
Imp is not regulated by miRNAs, Syp, or Chinmo in GSCs, related to Fig. 4. (A–D) Confocal images of Imp protein (green) in D28 *nos>control-i* (A) or *nos>Dcr-1-i* (B) or *nos>drosha-i* (C) testes, quantified in D. Arrowheads indicate Imp expression in GSCs. (E, F) Confocal images of Syp protein (green) in D2 (E) or D42 (F) testes. Arrowheads mark GSCs. (G-J) Confocal images of Imp protein (green) in D14 *nos>control-i* (G) or *nos>syp-i* (H) testes, quantified in I. Arrowheads indicate Imp expression in GSCs. Graph (J) showing Chinmo protein expression in D2 or D42 *nos>control-i* (gray bars) and D42 *nos>syp-i* (orange bars) testes. (K-M) Confocal images of Imp protein (green) in D42 testes of *nos>control-i* (K) or *nos>chinmo* (L) testes, quantified in M. Arrowheads mark Imp expression in GSCs. Asterisks indicate niche cells. Vasa marks germ cells and Fas3 labels niche cells. Scale bars = 10 μm. Error bars in I, J, M represent SEM. *** P ≤ 0.001; n.s. indicates no significance, as determined by Student’s t-test (see STAR Methods).

**Figure S5.**
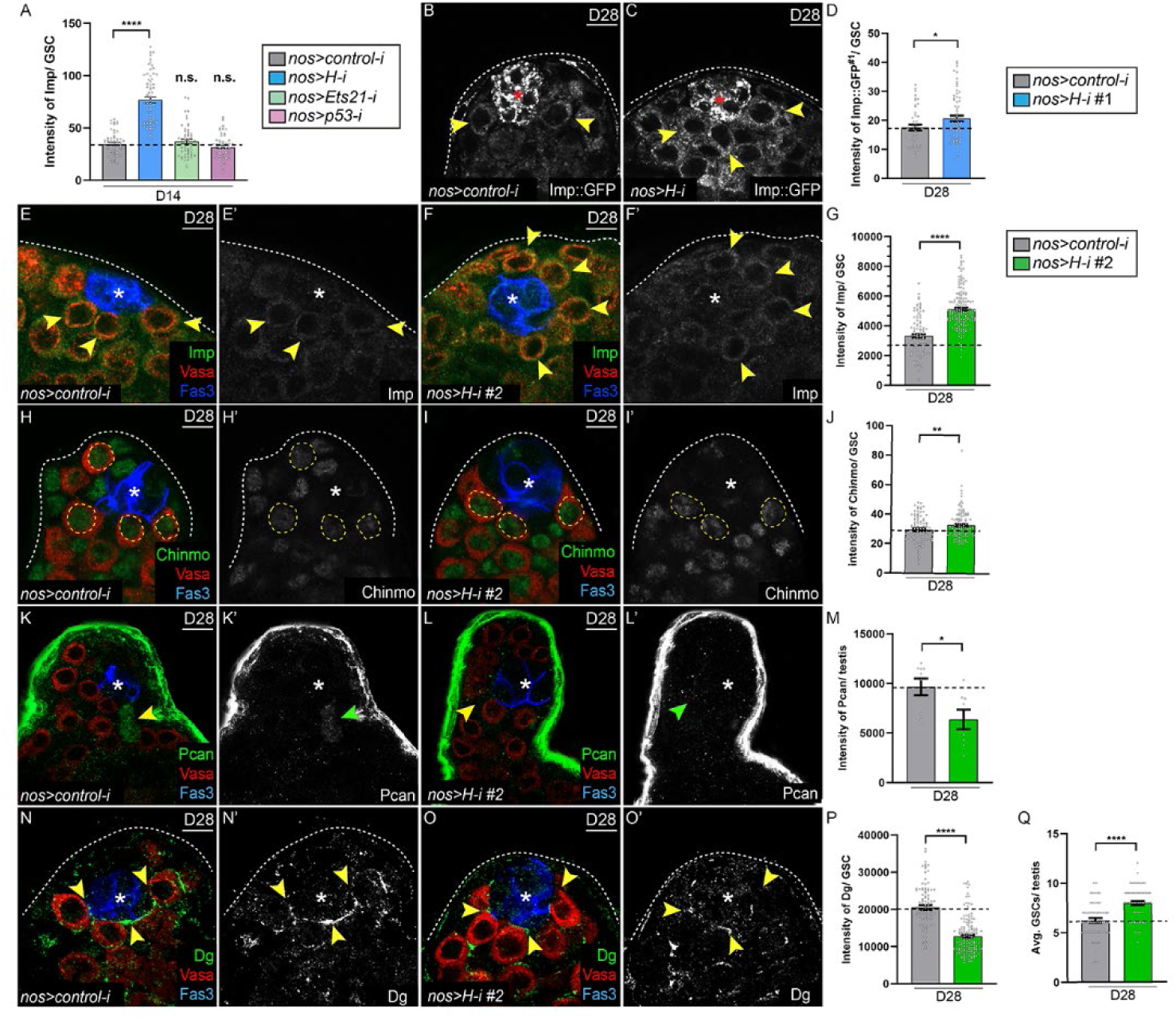
Hairless represses Imp protein in GSCs, related to Fig. 5. (A) Graph showing the intensity of Imp protein in D14 *nos>control-i* (gray bar)*, nos>H-i* (blue bar), *nos>Ets21-i* (green bar), or *nos>p53-i* (purple bar) testes. (B–D) Confocal images of Imp::GFP (gray) in D28 *nos>control-i* (B) or *nos>H-i* (C) testes, quantified in D. Arrowheads indicate Imp::GFP expression in GSCs. (E–G) Confocal images of Imp protein (green) in D28 *nos>control-i* (E) or *nos>H-i#2* (F) testes, quantified in G. Arrowheads mark Imp in GSCs. (H–J) Confocal images of Chinmo protein (green) in D28 *nos>control-i* (H) or *nos>H-i#2* (I) testes, quantified in J. Dashed lines indicate GSCs. (K–M) Confocal images of Pcan protein (green) in D28 *nos>control-i* (K) or *nos>H-i#2* (L) testes, quantified in M. Arrowhead indicates ectopic Pcan surrounding the niche. (N–P) Confocal images of Dg (green) at the GSC–niche interface in D28 *nos>control-i* (N) or *nos>H-i#2* (O) testes, quantified in P. Arrowheads mark Dg at the GSC–niche interface. (Q) Graph of the total number of GSCs in D28 *nos>control-i* (gray bar) or *nos>H-i#2* (green bar) testes. Asterisks indicate niche cells. Vasa marks germ cells and Fas3 labels niche cells. Scale bars = 10 μm. Error bars in A, D, G, J, M, P, Q represent SEM. * P ≤ 0.05; ** P ≤ 0.01; **** P ≤ 0.0001 by Student’s t-test (see STAR Methods).

**Figure S6.**
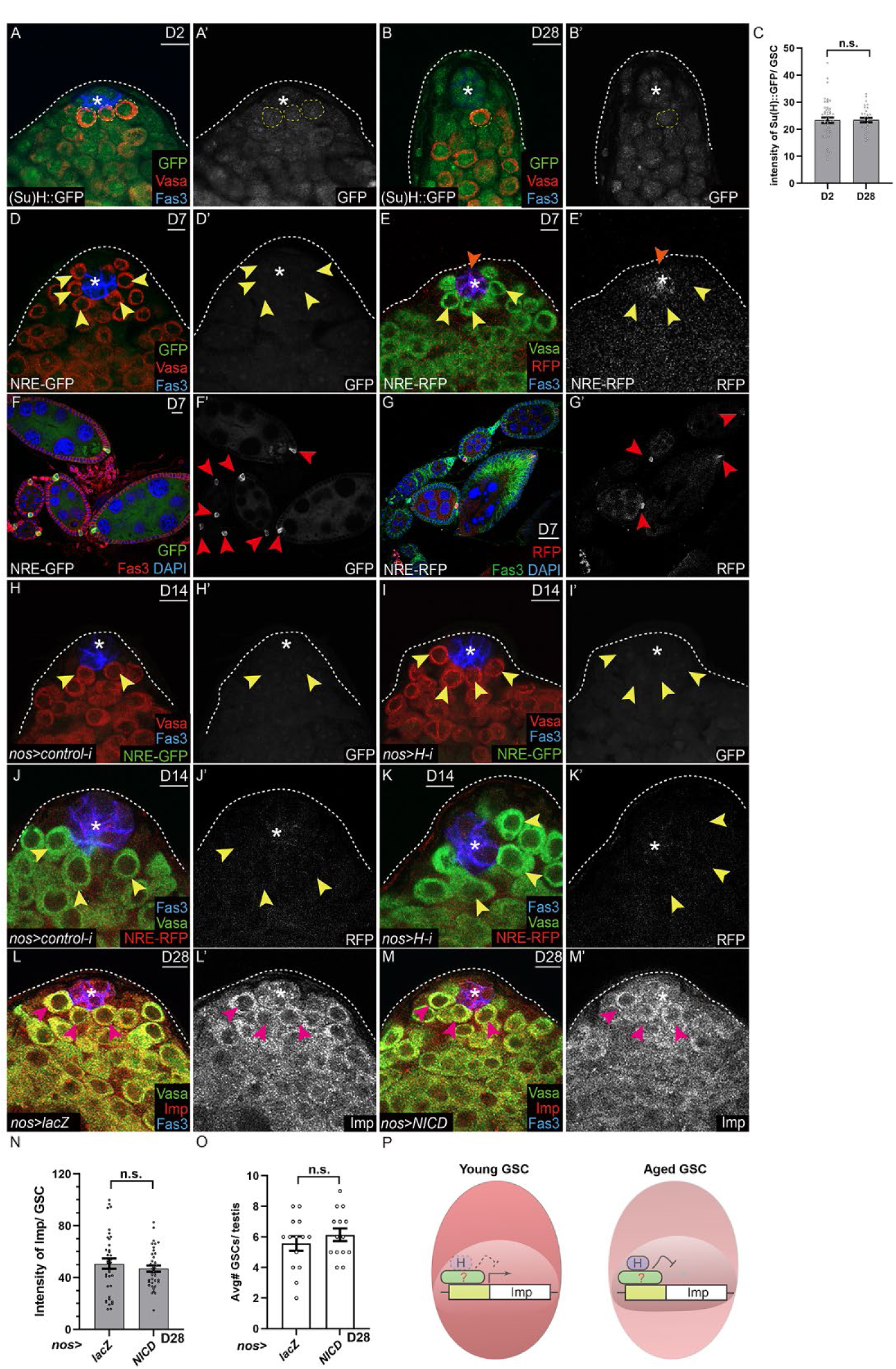
Notch signaling is not observed in adult testes, related to Fig. 5. (A-C) Confocal images of Suppressor of Hairless::GFP ((Su)H::GFP) (green) in D2 (A) or D28 (B) testes, quantified in C. Dashed lines indicate GSCs. (D-G) Confocal images of NRE-GFP (D, F) and NRE-RFP (E, G) in D7 testes (E, F) or in D7 ovaries (F, G). Yellow arrowheads indicate GSCs (D, E), while the orange arrowhead (E) indicates NRE-RFP expression in niche cells. Arrowheads in F, G indicate ovarian polar cells that express NRE-GFP (F) or NRE-RFP (G) reporters. (H-K) Confocal images of NRE-GFP (H, I) and NRE-RFP (J, K) in D14 *nos>control-i* (H, J) or *nos>H-i* (I, K) testes. Arrowheads indicate GSCs. (L-O) Confocal images of Imp protein (red) in D28 *nos>lacZ* (L) or *nos>NICD* (M) testes, quantified in N. Arrowheads indicate GSCs. Graph in O showing the average number of GSCs in D28 *nos>lacZ* or *nos>NICD* testes. (P) Model of H repression of *Imp*. H interacts with an as-yet unknown protein (green shape with question mark) to repress *Imp* gene expression. In young GSCs (left), H (shape outlined with dashed line) is low, permitting robust *Imp* gene expression. At later adult stages (right), H (purple box) increases, leading to repression of the *Imp* gene. H-mediated repression of *Imp* occurs independently of H’s canonical role in the Notch pathway. Asterisks indicate niche cells. Vasa marks germ cells and Fas3 labels niche cells except for G and H, where Fas3 marks ovarian follicle cells. Scale bar = 10 μM Error bars in C, N, O represent SEM. n.s. indicates no significance as assessed by Student’s t-test (see STAR Methods).

**Figure S7.**
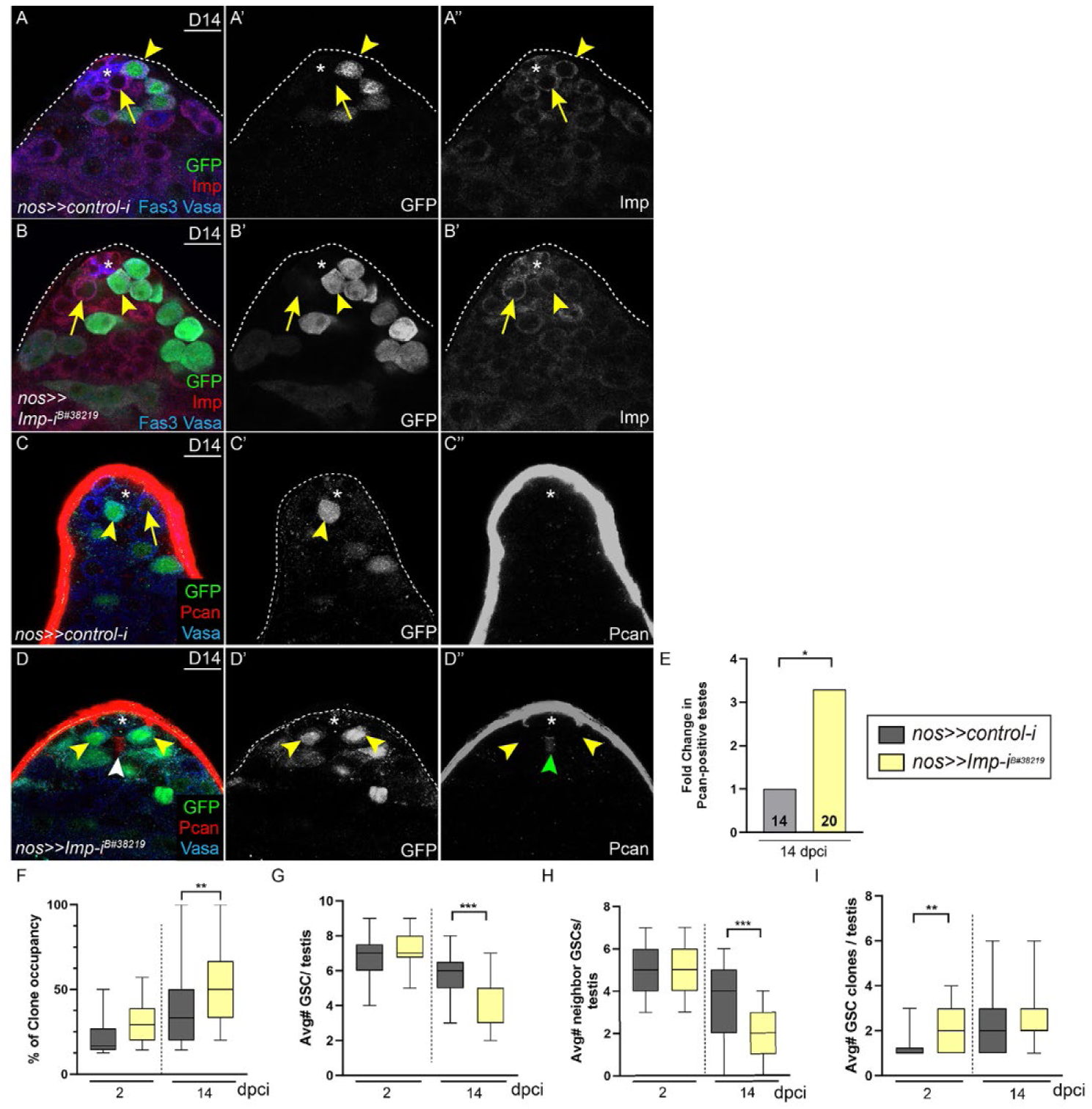
*Imp*-mutant GSCs are more competitive, causing niche remodeling, and evicting non-mutant GSC neighbors, related to Fig. 7. (A-B) Confocal image of Imp protein (red) in testes with *nos>>control-i* GSC clones (A) or *nos>>Imp-i* GSC clones (B) at 14 dpci. Arrowheads indicate the GFP-positive GSC clone. Arrows indicate non-mutant neighboring GSCs. (C-E) Confocal image of Pcan protein (red) in the testis with *nos>>control-i* GSC clones (C) or *nos>>Imp-i* GSC clones (D) at 14 dpci. Arrowheads indicate GFP-positive GSC clones. Magenta arrowhead indicates ectopic Pcan deposition around the niche in testes with *nos>>Imp-i* GSC clones, quantified in E. Sample size for each genotype is indicated in the bar. (F-I) Box and whisker plots showing clone occupancy (F), average total number of GSCs (G), average number of non-mutant GSC neighbors (H), and average number of GSC clones (I) in testes with control *nos>>control-i* GSC clones (gray bars) or *nos>>Imp-i* GSC clones (yellow bars) at 2 and 14 dpci. Asterisks indicated niche cells. Vasa marks germ cells and Fas3 labels niche cells. Scale bar = 10 μM In G-I, error bars represent SEM. * P ≤ 0.05; ** P ≤ 0.01; *** P ≤ 0.001 as assessed by Student’s t-test for F-I and by χ^2^ test for E (see STAR Methods).

## References

1. Morrison, S.J., and Spradling, A.C. (2008). Stem cells and niches: mechanisms that promote stem cell maintenance throughout life. Cell 132, 598–611. 10.1016/j.cell.2008.01.038.

2. Brunet, A., Goodell, M.A., and Rando, T.A. (2023). Ageing and rejuvenation of tissue stem cells and their niches. Nature reviews. Molecular cell biology 24, 45–62. 10.1038/s41580-022-00510-w.

3. Oh, J., Lee, Y.D., and Wagers, A.J. (2014). Stem cell aging: mechanisms, regulators and therapeutic opportunities. Nature medicine 20, 870–880. 10.1038/nm.3651.

4. Harris, I.D., Fronczak, C., Roth, L., and Meacham, R.B. (2011). Fertility and the aging male. Rev Urol 13, e184–190.

5. Khandwala, Y.S., Zhang, C.A., Lu, Y., and Eisenberg, M.L. (2017). The age of fathers in the USA is rising: an analysis of 168 867 480 births from 1972 to 2015. Hum Reprod 32, 2110–2116. 10.1093/humrep/dex267.

6. Johnson, L., Zane, R.S., Petty, C.S., and Neaves, W.B. (1984). Quantification of the human Sertoli cell population: its distribution, relation to germ cell numbers, and age-related decline. Biology of reproduction 31, 785–795. 10.1095/biolreprod31.4.785.

7. Paniagua, R., Nistal, M., Amat, P., Rodriguez, M.C., and Martin, A. (1987). Seminiferous tubule involution in elderly men. Biology of reproduction 36, 939–947. 10.1095/biolreprod36.4.939.

8. Pohl, E., Hoffken, V., Schlatt, S., Kliesch, S., Gromoll, J., and Wistuba, J. (2019). Ageing in men with normal spermatogenesis alters spermatogonial dynamics and nuclear morphology in Sertoli cells. Andrology 7, 827–839. 10.1111/andr.12665.

9. Buchwalter, A., and Hetzer, M.W. (2017). Nucleolar expansion and elevated protein translation in premature aging. Nature communications 8, 328. 10.1038/s41467-017-00322-z.

10. Ryu, B.Y., Orwig, K.E., Oatley, J.M., Avarbock, M.R., and Brinster, R.L. (2006). Effects of aging and niche microenvironment on spermatogonial stem cell self-renewal. Stem cells (Dayton, Ohio) 24, 1505–1511. 10.1634/stemcells.2005-0580.

11. Sato, T., Katagiri, K., Yokonishi, T., Kubota, Y., Inoue, K., Ogonuki, N., Matoba, S., Ogura, A., and Ogawa, T. (2011). In vitro production of fertile sperm from murine spermatogonial stem cell lines. Nature communications 2, 472. 10.1038/ncomms1478.

12. Liggett, L.A., and DeGregori, J. (2017). Changing mutational and adaptive landscapes and the genesis of cancer. Biochim Biophys Acta Rev Cancer 1867, 84–94. 10.1016/j.bbcan.2017.01.005.

13. Florez, M.A., Tran, B.T., Wathan, T.K., DeGregori, J., Pietras, E.M., and King, K.Y. (2022). Clonal hematopoiesis: Mutation-specific adaptation to environmental change. Cell stem cell 29, 882–904. 10.1016/j.stem.2022.05.006.

14. Jaiswal, S., and Ebert, B.L. (2019). Clonal hematopoiesis in human aging and disease. Science (New York, N.Y 366. 10.1126/science.aan4673.

15. Snippert, H.J., Schepers, A.G., van Es, J.H., Simons, B.D., and Clevers, H. (2014). Biased competition between Lgr5 intestinal stem cells driven by oncogenic mutation induces clonal expansion. EMBO reports 15, 62–69. 10.1002/embr.201337799.

16. Snippert, H.J., van der Flier, L.G., Sato, T., van Es, J.H., van den Born, M., Kroon-Veenboer, C., Barker, N., Klein, A.M., van Rheenen, J., Simons, B.D., and Clevers, H. (2010). Intestinal crypt homeostasis results from neutral competition between symmetrically dividing Lgr5 stem cells. Cell 143, 134–144.

17. Lopez-Garcia, C., Klein, A.M., Simons, B.D., and Winton, D.J. (2010). Intestinal stem cell replacement follows a pattern of neutral drift. Science (New York, N.Y 330, 822–825. 10.1126/science.1196236.

18. Amoyel, M., Simons, B.D., and Bach, E.A. (2014). Neutral competition of stem cells is skewed by proliferative changes downstream of Hh and Hpo. The EMBO journal 33, 2295–2313. 10.15252/embj.201387500.

19. Vermeulen, L., Morrissey, E., van der Heijden, M., Nicholson, A.M., Sottoriva, A., Buczacki, S., Kemp, R., Tavare, S., and Winton, D.J. (2013). Defining stem cell dynamics in models of intestinal tumor initiation. Science (New York, N.Y 342, 995–998. 10.1126/science.1243148.

20. van Neerven, S.M., de Groot, N.E., Nijman, L.E., Scicluna, B.P., van Driel, M.S., Lecca, M.C., Warmerdam, D.O., Kakkar, V., Moreno, L.F., Vieira Braga, F.A., et al. (2021). Apc-mutant cells act as supercompetitors in intestinal tumour initiation. Nature 594, 436–441. 10.1038/s41586-021-03558-4.

21. Flanagan, D.J., Pentinmikko, N., Luopajarvi, K., Willis, N.J., Gilroy, K., Raven, A.P., McGarry, L., Englund, J.I., Webb, A.T., Scharaw, S., et al. (2021). NOTUM from Apc-mutant cells biases clonal competition to initiate cancer. Nature 594, 430–435. 10.1038/s41586-021-03525-z.

22. Herms, A., Colom, B., Piedrafita, G., Kalogeropoulou, A., Banerjee, U., King, C., Abby, E., Murai, K., Caseda, I., Fernandez-Antoran, D., et al. (2024). Organismal metabolism regulates the expansion of oncogenic PIK3CA mutant clones in normal esophagus. Nature genetics 56, 2144–2157. 10.1038/s41588-024-01891-8.

23. Colom, B., Herms, A., Hall, M.W.J., Dentro, S.C., King, C., Sood, R.K., Alcolea, M.P., Piedrafita, G., Fernandez-Antoran, D., Ong, S.H., et al. (2021). Mutant clones in normal epithelium outcompete and eliminate emerging tumours. Nature 598, 510–514. 10.1038/s41586-021-03965-7.

24. Martincorena, I., Fowler, J.C., Wabik, A., Lawson, A.R.J., Abascal, F., Hall, M.W.J., Cagan, A., Murai, K., Mahbubani, K., Stratton, M.R., et al. (2018). Somatic mutant clones colonize the human esophagus with age. Science (New York, N.Y 362, 911–917. 10.1126/science.aau3879.

25. Roan, H.Y., Tseng, T.L., and Chen, C.H. (2021). Whole-body clonal mapping identifies giant dominant clones in zebrafish skin epidermis. Development (Cambridge, England) 148. 10.1242/dev.199669.

26. Goriely, A., and Wilkie, A.O. (2012). Paternal age effect mutations and selfish spermatogonial selection: causes and consequences for human disease. Am J Hum Genet 90, 175–200. 10.1016/j.ajhg.2011.12.017.

27. Wood, K.A., and Goriely, A. (2022). The impact of paternal age on new mutations and disease in the next generation. Fertility and sterility 118, 1001–1012. 10.1016/j.fertnstert.2022.10.017.

28. Hodge, R.A., and Bach, E.A. (2024). Mechanisms of Germline Stem Cell Competition across Species. Life (Basel) 14. 10.3390/life14101251.

29. Wallenfang, M.R., Nayak, R., and DiNardo, S. (2006). Dynamics of the male germline stem cell population during aging of Drosophila melanogaster. Aging cell 5, 297–304. 10.1111/j.1474-9726.2006.00221.x.

30. Boyle, M., Wong, C., Rocha, M., and Jones, D.L. (2007). Decline in self-renewal factors contributes to aging of the stem cell niche in the Drosophila testis. Cell stem cell 1, 470–478. 10.1016/j.stem.2007.08.002.

31. Toledano, H., D’Alterio, C., Czech, B., Levine, E., and Jones, D.L. (2012). The let-7-Imp axis regulates ageing of the Drosophila testis stem-cell niche. Nature 485, 605–610. 10.1038/nature11061.

32. Herrera, S.C., Sainz de la Maza, D., Grmai, L., Margolis, S., Plessel, R., Burel, M., O’Connor, M., Amoyel, M., and Bach, E.A. (2021). Proliferative stem cells maintain quiescence of their niche by secreting the Activin inhibitor Follistatin. Dev Cell 56, 2284–2294 e2286. 10.1016/j.devcel.2021.07.010.

33. Tseng, C.Y., Burel, M., Cammer, M., Harsh, S., Flaherty, M.S., Baumgartner, S., and Bach, E.A. (2022). chinmo-mutant spermatogonial stem cells cause mitotic drive by evicting non-mutant neighbors from the niche. Dev Cell 57, 80–94 e87. 10.1016/j.devcel.2021.12.004.

34. Greenspan, L.J., de Cuevas, M., and Matunis, E. (2015). Genetics of gonadal stem cell renewal. Annual review of cell and developmental biology 31, 291–315. 10.1146/annurev-cellbio-100913-013344.

35. Leatherman, J.L., and Dinardo, S. (2008). Zfh-1 controls somatic stem cell self-renewal in the Drosophila testis and nonautonomously influences germline stem cell self-renewal. Cell stem cell 3, 44–54.

36. Leatherman, J.L., and Dinardo, S. (2010). Germline self-renewal requires cyst stem cells and stat regulates niche adhesion in Drosophila testes. Nature cell biology 12, 806–811.

37. Tulina, N., and Matunis, E. (2001). Control of stem cell self-renewal in Drosophila spermatogenesis by JAK-STAT signaling. Science (New York, N.Y 294, 2546–2549.

38. Kiger, A.A., Jones, D.L., Schulz, C., Rogers, M.B., and Fuller, M.T. (2001). Stem cell self-renewal specified by JAK-STAT activation in response to a support cell cue. Science (New York, N.Y 294, 2542–2545.

39. Kawase, E., Wong, M.D., Ding, B.C., and Xie, T. (2004). Gbb/Bmp signaling is essential for maintaining germline stem cells and for repressing bam transcription in the Drosophila testis. Development (Cambridge, England) 131, 1365–1375.

40. Shivdasani, A.A., and Ingham, P.W. (2003). Regulation of stem cell maintenance and transit amplifying cell proliferation by tgf-beta signaling in Drosophila spermatogenesis. Curr Biol 13, 2065–2072.

41. Inaba, M., Buszczak, M., and Yamashita, Y.M. (2015). Nanotubes mediate niche-stem-cell signalling in the Drosophila testis. Nature. 10.1038/nature14602.

42. Wu, Y.C., Chen, C.H., Mercer, A., and Sokol, N.S. (2012). Let-7-complex microRNAs regulate the temporal identity of Drosophila mushroom body neurons via chinmo. Dev Cell 23, 202–209. 10.1016/j.devcel.2012.05.013.

43. Yang, L., Chen, D., Duan, R., Xia, L., Wang, J., Qurashi, A., Jin, P., and Chen, D. (2007). Argonaute 1 regulates the fate of germline stem cells in Drosophila. Development (Cambridge, England) 134, 4265–4272. 10.1242/dev.009159.

44. Flaherty, M.S., Salis, P., Evans, C.J., Ekas, L.A., Marouf, A., Zavadil, J., Banerjee, U., and Bach, E.A. (2010). chinmo is a functional effector of the JAK/STAT pathway that regulates eye development, tumor formation, and stem cell self-renewal in Drosophila. Dev Cell 18, 556–568. 10.1016/j.devcel.2010.02.006.

45. Liu, Z., Yang, C.P., Sugino, K., Fu, C.C., Liu, L.Y., Yao, X., Lee, L.P., and Lee, T. (2015). Opposing intrinsic temporal gradients guide neural stem cell production of varied neuronal fates. Science (New York, N.Y 350, 317–320. 10.1126/science.aad1886.

46. Narbonne-Reveau, K., Lanet, E., Dillard, C., Foppolo, S., Chen, C.H., Parrinello, H., Rialle, S., Sokol, N.S., and Maurange, C. (2016). Neural stem cell-encoded temporal patterning delineates an early window of malignant susceptibility in Drosophila. eLife 5. 10.7554/eLife.13463.

47. Muller, D., Kugler, S.J., Preiss, A., Maier, D., and Nagel, A.C. (2005). Genetic modifier screens on Hairless gain-of-function phenotypes reveal genes involved in cell differentiation, cell growth and apoptosis in Drosophila melanogaster. Genetics 171, 1137–1152. 10.1534/genetics.105.044453.

48. Mundorf, J., Donohoe, C.D., McClure, C.D., Southall, T.D., and Uhlirova, M. (2019). Ets21c Governs Tissue Renewal, Stress Tolerance, and Aging in the Drosophila Intestine. Cell reports 27, 3019–3033 e3015. 10.1016/j.celrep.2019.05.025.

49. Li, H., Janssens, J., De Waegeneer, M., Kolluru, S.S., Davie, K., Gardeux, V., Saelens, W., David, F.P.A., Brbic, M., Spanier, K., et al. (2022). Fly Cell Atlas: A single-nucleus transcriptomic atlas of the adult fruit fly. Science (New York, N.Y 375, eabk2432. 10.1126/science.abk2432.

50. Maier, D. (2006). Hairless: the ignored antagonist of the Notch signalling pathway. Hereditas 143, 212–221. 10.1111/j.2007.0018-0661.01971.x.

51. Nagel, A.C., Krejci, A., Tenin, G., Bravo-Patino, A., Bray, S., Maier, D., and Preiss, A. (2005). Hairless-mediated repression of notch target genes requires the combined activity of Groucho and CtBP corepressors. Molecular and cellular biology 25, 10433–10441. 10.1128/MCB.25.23.10433-10441.2005.

52. Barolo, S., Stone, T., Bang, A.G., and Posakony, J.W. (2002). Default repression and Notch signaling: Hairless acts as an adaptor to recruit the corepressors Groucho and dCtBP to Suppressor of Hairless. Genes & development 16, 1964–1976. 10.1101/gad.987402.

53. Shyu, L.F., Sun, J., Chung, H.M., Huang, Y.C., and Deng, W.M. (2009). Notch signaling and developmental cell-cycle arrest in Drosophila polar follicle cells. Molecular biology of the cell 20, 5064–5073. 10.1091/mbc.e09-01-0004.

54. Lee, T., and Luo, L. (1999). Mosaic analysis with a repressible cell marker for studies of gene function in neuronal morphogenesis. Neuron 22, 451–461.

55. Ma, X., Wang, S., Do, T., Song, X., Inaba, M., Nishimoto, Y., Liu, L.P., Gao, Y., Mao, Y., Li, H., et al. (2014). Piwi is required in multiple cell types to control germline stem cell lineage development in the Drosophila ovary. PloS one 9, e90267. 10.1371/journal.pone.0090267.

56. Zhao, R., Xuan, Y., Li, X., and Xi, R. (2008). Age-related changes of germline stem cell activity, niche signaling activity and egg production in Drosophila. Aging cell 7, 344–354. 10.1111/j.1474-9726.2008.00379.x.

57. Yousef, H., Morgenthaler, A., Schlesinger, C., Bugaj, L., Conboy, I.M., and Schaffer, D.V. (2015). Age-Associated Increase in BMP Signaling Inhibits Hippocampal Neurogenesis. Stem cells (Dayton, Ohio) 33, 1577–1588. 10.1002/stem.1943.

58. Valletta, S., Thomas, A., Meng, Y., Ren, X., Drissen, R., Sengul, H., Di Genua, C., and Nerlov, C. (2020). Micro-environmental sensing by bone marrow stroma identifies IL-6 and TGFbeta1 as regulators of hematopoietic ageing. Nature communications 11, 4075. 10.1038/s41467-020-17942-7.

59. Egerman, M.A., Cadena, S.M., Gilbert, J.A., Meyer, A., Nelson, H.N., Swalley, S.E., Mallozzi, C., Jacobi, C., Jennings, L.L., Clay, I., et al. (2015). GDF11 Increases with Age and Inhibits Skeletal Muscle Regeneration. Cell metabolism 22, 164–174. 10.1016/j.cmet.2015.05.010.

60. Johannes, B., and Preiss, A. (2002). Wing vein formation in Drosophila melanogaster: hairless is involved in the cross-talk between Notch and EGF signaling pathways. Mechanisms of development 115, 3–14. 10.1016/s0925-4773(02)00083-7.

61. Song, X., Wong, M.D., Kawase, E., Xi, R., Ding, B.C., McCarthy, J.J., and Xie, T. (2004). Bmp signals from niche cells directly repress transcription of a differentiation-promoting gene, bag of marbles, in germline stem cells in the Drosophila ovary. Development (Cambridge, England) 131, 1353–1364. 10.1242/dev.01026.

62. Suo, M., Rommelfanger, M.K., Chen, Y., Amro, E.M., Han, B., Chen, Z., Szafranski, K., Chakkarappan, S.R., Boehm, B.O., MacLean, A.L., and Rudolph, K.L. (2022). Age-dependent effects of Igf2bp2 on gene regulation, function, and aging of hematopoietic stem cells in mice. Blood 139, 2653–2665. 10.1182/blood.2021012197.

63. Van Doren, M., Williamson, A.L., and Lehmann, R. (1998). Regulation of zygotic gene expression in Drosophila primordial germ cells. Curr Biol 8, 243–246.

64. Ekas, L.A., Baeg, G.H., Flaherty, M.S., Ayala-Camargo, A., and Bach, E.A. (2006). JAK/STAT signaling promotes regional specification by negatively regulating wingless expression in Drosophila. Development (Cambridge, England) 133, 4721–4729. 10.1242/dev.02675.

65. Baksa, K., Parke, T., Dobens, L.L., and Dearolf, C.R. (2002). The Drosophila STAT protein, stat92E, regulates follicle cell differentiation during oogenesis. Developmental biology 243, 166–175. 10.1006/dbio.2001.0539.

66. Silver, D.L., and Montell, D.J. (2001). Paracrine signaling through the JAK/STAT pathway activates invasive behavior of ovarian epithelial cells in Drosophila. Cell 107, 831–841.

67. Li, M.A., Alls, J.D., Avancini, R.M., Koo, K., and Godt, D. (2003). The large Maf factor Traffic Jam controls gonad morphogenesis in Drosophila. Nature cell biology 5, 994–1000. 10.1038/ncb1058.

68. Zhu, S., Lin, S., Kao, C.F., Awasaki, T., Chiang, A.S., and Lee, T. (2006). Gradients of the Drosophila Chinmo BTB-zinc finger protein govern neuronal temporal identity. Cell 127, 409–422.

69. Xu, T., and Rubin, G.M. (1993). Analysis of genetic mosaics in developing and adult Drosophila tissues. Development (Cambridge, England) 117, 1223–1237.

70. Brand, A.H., and Perrimon, N. (1993). Targeted gene expression as a means of altering cell fates and generating dominant phenotypes. Development (Cambridge, England) 118, 401–415.

71. McGuire, S.E., Mao, Z., and Davis, R.L. (2004). Spatiotemporal gene expression targeting with the TARGET and gene-switch systems in Drosophila. Sci STKE 2004, pl6.

72. Zimmerman, S.G., Peters, N.C., Altaras, A.E., and Berg, C.A. (2013). Optimized RNA ISH, RNA FISH and protein-RNA double labeling (IF/FISH) in Drosophila ovaries. Nature protocols 8, 2158–2179. 10.1038/nprot.2013.136.

